# Assembly and activation of the AAA^+^ motor of the multi-subunit cobaltochelatase

**DOI:** 10.64898/2026.10.09.757910

**Authors:** Ana C. Moura, Benjamin Wiseman, Esra Bozkurt, Evelyne Deery, Richard W. Pickersgill, Amanda A. Brindley, Andrew Hitchcock, Martin Högbom, Martin J. Warren, Dmitry Shvarev

**Affiliations:** Department of Biology/Chemistry - Structural Biology of Photosynthetic Microorganisms, University of Osnabrück, Osnabrück 49076, Germany; Center of Cellular Nanoanalytics Osnabrück (CellNanOs); Osnabrück University, Osnabrück 49076, Germany; Stockholm University, Department of Biochemistry and Biophysics, Stockholm 10691, Sweden; School of Natural Sciences, University of Kent, Canterbury CT2 7NJ, United Kingdom; School of Biological and Behavioural Sciences, Queen Mary University of London, London E1 4NS, United Kingdom; Plants, Photosynthesis and Soil, School of Biosciences, University of Sheffield, Sheffield, UK; Molecular Microbiology: Biochemistry to Disease, School of Biosciences, University of Sheffield, Sheffield, United Kingdom; Science for Life Laboratory (SciLifeLab), Stockholm University, Stockholm, Sweden; Quadram Institute Bioscience, Norwich Research Park, Norwich NR4 7UA, United Kingdom; University of East Anglia, Norwich NR4 7TJ, United Kingdom

## Abstract

Cobalamin is a complex yet essential cobalt-containing cofactor synthesized exclusively by prokaryotes and archaea. Aerobic cobalamin biosynthesis requires the CobNST multi-subunit class I cobaltochelatase for cobalt incorporation. Cobalt chelation is dependent on the energy transfer from the CobST AAA^+^/VWA motor unit to the substrate-binding catalytic unit (CobN). However, the mechanism for assembly and function of CobNST cobaltochelatase remains poorly understood. Here, we used cryo-EM to solve the structures of the AAA^+^ motor CobS, in the absence and presence of ATP and its VWA-partner protein CobT. Our data reveal that the AAA^+^ CobS assembles into various conformations, including hexameric planar rings and different spiral left-handed assemblies: hexamers, double rings and filaments. In the absence of nucleotide, the CobT N-terminal domain interacts with CobS left-handed oligomers, remaining in a half-assembled, nonfunctional motor unit. In the CobST complex, ATP induces conformational rearrangement of CobS from left-handed helical oligomers to a right-handed closed hexamer. The simultaneous repositioning of the CobT VWA domain to the AAA^+^ central pore shifts the motor unit to a functional state, ready-to-drive CobN-catalyzed chelation reaction. These findings define the mechanism for assembly and activation of the class I cobaltochelatase CobST motor.

## Introduction

Cobamides comprise a structurally diverse family of cobalt-containing tetrapyrrole cofactors, of which cobalamin is the best-characterized member ^1^. All cobamides share a contracted corrin macrocycle that coordinates cobalt through four nitrogen atoms and comprise lower and upper ligands occupying the axial coordination sites of the cobalt ion. The structural diversity within cobalamins arises mostly from variation in the lower axial ligand supplied through the nucleotide loop ^1^. The combination of a highly modified tetrapyrrole framework and an organometallic cobalt centre gives cobalamin access to chemistry that is unusual among biological cofactors ^2–4^. The molecule commonly referred to as vitamin B_12_ is cyanocobalamin, in which cyanide occupies the upper ligand of the central cobalt ion, whereas the biologically active methylcobalamin and 5′-deoxyadenosylcobalamin forms contain methyl and adenosyl upper ligands, respectively ^5^. In humans, vitamin B_12_ is required by only two known enzymes, methionine synthase and methylmalonyl-CoA mutase, yet their enzymatic reactions illustrate the remarkable catalytic versatility conferred by the corrinoid cobalt centre ^4^. A deficiency of vitamin B_12_ can have profound metabolic consequences in humans, including megaloblastic anaemia, hyperhomocysteinaemia and elevated methylmalonic acid ^6,7^. Therefore, the unusual catalytic properties of cobalamin ultimately depend on the correct incorporation of cobalt into the corrin macrocycle during its biosynthesis.

The structural complexity of cobalamin is reflected in an elaborate biosynthetic pathway involving more than 30 enzymatic steps and comprising two broadly distinct routes, conventionally referred to as the aerobic and anaerobic pathways ^5,8,9^. Although both pathways ultimately generate the same corrinoid cofactor, they differ in the stage at which cobalt is inserted and in the molecular machinery responsible for metal chelation ^10^. In the anaerobic pathway, characteristic of many archaea and anaerobic bacteria, cobalt is inserted early into sirohydrochlorin by small, ATP-independent class II cobaltochelatases, including CbiK and the CbiX family, to generate cobalt-factor II before corrin-ring contraction and modification ^11–14^. In contrast, the aerobic pathway delays cobalt insertion until after corrin-ring contraction, where cobalt is incorporated into hydrogenobyrinic acid *a,c*-diamide (HBAD) by the ATP-dependent class I cobaltochelatase CobNST to generate cobyrinic acid *a,c*-diamide (CBAD), the first cobalt-containing intermediate of the aerobic pathway ^15–17^. The class I cobaltochelatase was first isolated by Debussche and colleagues from *Pseudomonas denitrificans* cell extracts ^18^ and consists of three proteins with distinct but coordinated functions: the large corrin-binding protein CobN (140 kDa), the ATPase associated with diverse cellular activities (AAA^+^) protein CobS (38 kDa) and the von Willebrand factor type A (VWA)-domain-containing adaptor protein CobT (80 kDa). Although these three proteins are sufficient to catalyse cobalt insertion *in vitro*, a fourth protein, the P-loop GTPase CobW, acts *in vivo* as a cobalt-binding metallochaperone that promotes delivery of cobalt to the CobNST system ^19,20^. The CobN, S and T components share striking functional and architectural similarities with the magnesium chelatase, BchHID/ ChlHID, which catalyses Mg^2+^ insertion during (bacterio)chlorophyll biosynthesis ^5,21^, suggesting an equivalent metal chelation mechanism for class I chelatases across different tetrapyrrole biosynthesis pathways.

In the CobNST complex, CobN binds the tetrapyrrole substrate and contains the site at which cobalt is ultimately inserted ^22^, whereas CobS and CobT assemble into an ATP-dependent motor module ^17^ that is likely to drive the conformational rearrangements required for chelation. The X-ray structure of *Mycobacterium tuberculosis* CobN reveals a six-domain architecture ^22^ resembling that of the homologous cyanobacterial magnesium-chelatase subunit ChlH ^23,24^. A strongly positively charged region at the interface of domains I and II, corresponding to the “head” and “neck” domains of the protein, has been implicated in binding of the negatively charged HBAD substrate ^22^. In contrast, residues implicated in cobalt binding and chelation are located in the more distal body region, centred around domain IV and its interfaces with neighbouring domains ^22^. More recent structural analysis has revealed HBAD in two distinct positions within CobN, demonstrating that the corrin substrate can undergo substantial repositioning within the protein and supporting a pathway of substrate movement from the “head”/”neck” interface towards the catalytic body of CobN [Deery *et. al.*, 2026, JBC, under review]. Together, these observations indicate that cobalt insertion requires an ordered repositioning of HBAD within CobN to bring the corrin macrocycle into a chelation-competent configuration at the domain-IV-centred metal-binding site.

The first structural evidence for the CobST motor revealed a two-tiered, approximately 520-kDa hexameric double-ring assembly formed by the AAA^+^ motor CobS and the adaptor protein CobT ^17^. A similar arrangement was observed by single-particle analysis for the ChlI and ChlD proteins of magnesium chelatase ^25^. CobS structure prediction (Fig. 1A and Supplementary Fig. 1A) shows the presence of pre-sensor 1 insert (PS1i) and helix 2 insert (H2i) sequences in the large AAA^+^ domain, which are associated with substrate recognition and modulation ^26^. PS1i motifs occur in numerous AAA^+^ proteins and are commonly associated with substrate engagement, whereas the H2i β-hairpin is characteristic of clade 6 and clade 7 AAA^+^ proteins and frequently participates in interactions with partner proteins. CobS belongs to the MoxR subfamily of AAA^+^ ATPases, whose members commonly function as molecular chaperones that use ATP hydrolysis to promote cofactor insertion, protein maturation or remodelling of macromolecular complexes ^27,28^. Other well-characterized members include ChlI of magnesium chelatase ^25,29^, the RuBisCO activase CbbQ ^30,31^ and the nitric oxide reductase maturation factor NorQ ^32,33^. MoxR AAA^+^ ATPases are frequently encoded adjacent to partner proteins containing a VWA domain, as is the case for CobS and CobT ^15,16^. In the CbbQO and NorQD systems, the VWA-containing proteins CbbO and NorD interact with the concave side of the AAA^+^ ring, with protruding regions of the partner proteins engaging the central pore of the ATPase ^30,32,33^. VWA domains frequently contain a metal-ion-dependent adhesion site (MIDAS) motif, typically comprising a DXSXS…T…D sequence, which coordinates a divalent metal ion, often Mg^2+^, and can stabilize metal-dependent interactions with partner proteins ^34^.

**Fig. 1.**
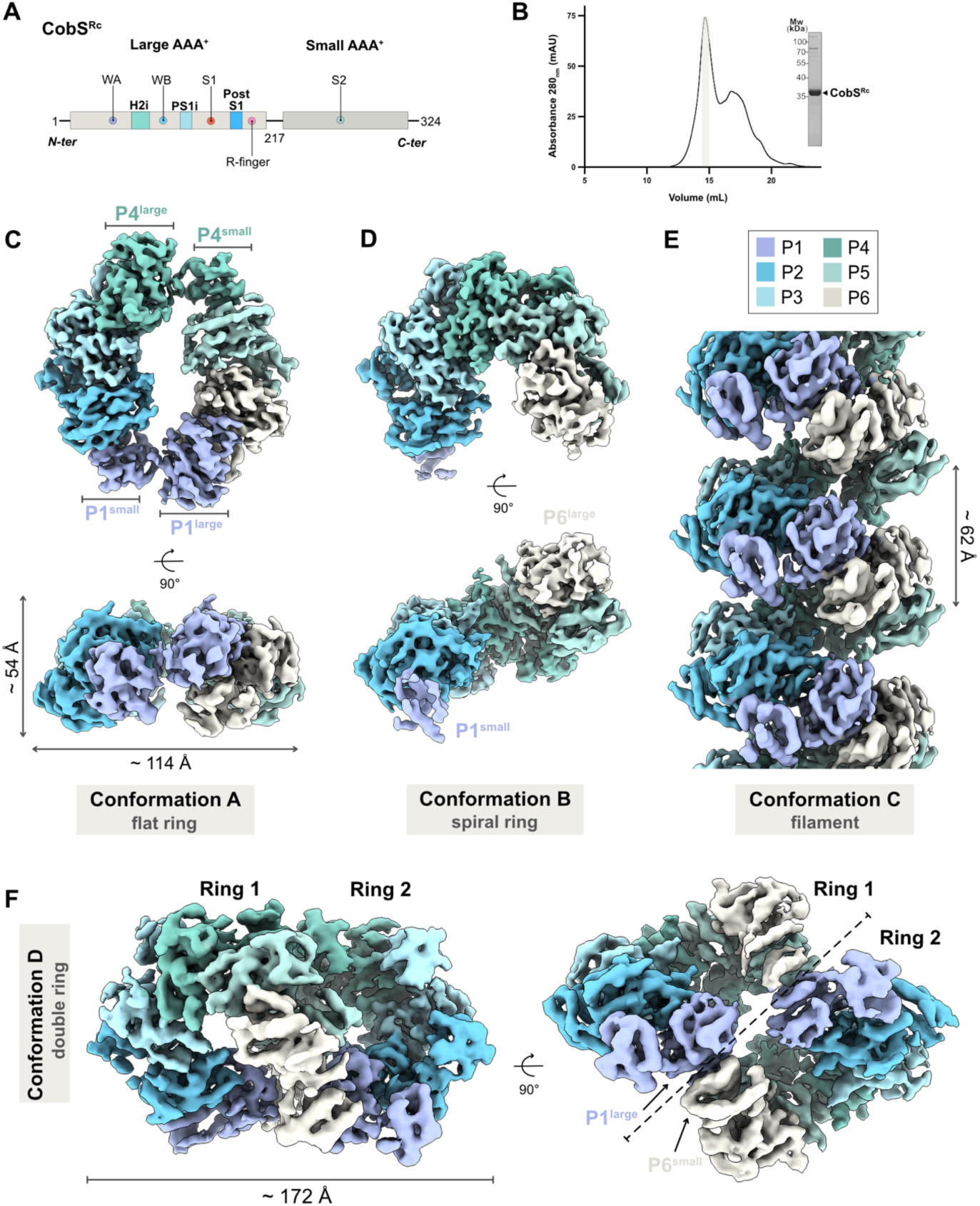
Overview of apo CobS^Rc^ structural diversity. **A.** Schematic representation based on the AlphaFold ^46^ prediction (Supplementary Fig. 1A) of the CobS^Rc^ AAA^+^ ATPase domains (large and small) and motifs. Highlighted motifs: Walker A (WA, Gly70 to Thr75), Walker B (WB, Ala133 to Glu138), sensor 1 (S1, Asn182), arginine finger (R-finger, Arg206) and sensor 2 (S2, Arg267). CobS^Rc^ structural insertions: helix 2 insert (H2i, green), pre-sensor 1 insert (PS1i, light blue) and post-sensor 1 loop insert (Post S1, blue). **B.** Representative size-exclusion chromatogram of CobS^Rc^ (Superose 6 Increase 10/300 column) and respective SDS-PAGE of the main peak fraction. **C.** Cryo-EM map of CobS^Rc^ hexameric flat ring (conformation A), top view (top image) and side view (bottom image). CobS^Rc^ individual subunits colored according to the scheme on the right (P1 to P6). **D.** Cryo-EM map of the partial hexameric spiral open ring (conformation B), views and coloring as in **C. E.** Side view of the cryo-EM map of CobS^Rc^ conformation C from helical refinement (see Supplementary Fig. 2 and Supplementary Fig. 9) showing a left-handed spiral arrangement of subunits. Coloring as in **C. F.** Cryo-EM map of CobS^Rc^ double ring (conformation D) viewed from two different angles, highlighting the interface between the two left-handed spiral rings. Coloring as in **C.**

The presence of analogous AAA^+^/VWA modules in both CobNST and magnesium chelatase therefore points towards a common mechanism, whereby an ATPase motor generates a power stroke, which is then transmitted via the VWA-domain partner to drive a potential rearrangement of the tetrapyrrole-binding catalytic subunit. However, despite extensive functional and structural investigation of magnesium chelatases ^23–25,29,35–40^ and cobaltochelatases ^17,18,20,22^, the molecular events that govern assembly of these complexes and couple ATP binding and hydrolysis to subsequent metal insertion remain unresolved.

Here, we use cryogenic electron microscopy (cryo-EM) to define the structural basis for assembly and nucleotide-dependent remodelling of the CobST motor unit from the purple photosynthetic bacterium *Rhodobacter capsulatus* (*R. capsulatus*) and the pathogenic bacterium *Brucella melitensis* (*B. melitensis*). Structures of the wild-type CobS AAA^+^ ATPase in multiple nucleotide states reveal a remarkable conformational plasticity, including left-handed helical hexamers, extended filaments, double dodecameric rings, and planar C2-symmetric closed hexamers. Structures of the corresponding CobST complexes further show that the N-terminal region of CobT initially associates with left-handed CobS assemblies, followed by engagement of the C-terminal VWA domain within the central pore of the ATPase under turnover conditions. This interaction coincides with a transition of the CobS hexamer from a left-to a right-handed configuration, while ATP hydrolysis likely drives repositioning of the CobT VWA domain within the CobS AAA^+^ ring. Together, these structural and functional data provide a molecular framework for assembly of the CobST motor and reveal how ATP-dependent conformational changes within CobS can be transmitted through CobT as a mechanical power stroke. These findings provide a basis for understanding how the CobST motor may engage and remodel CobN, rendering it chelation-competent and driving cobalt insertion into HBAD.

## Results

### The structural plasticity of the AAA^+^ CobS^Rc^

We overexpressed and purified the C-terminal 6x-His-tagged cobaltochelatase AAA^+^ motor component from *R. capsulatus* in *E*. *coli* (CobS^Rc^-6xHis, hereafter CobS^Rc^, Fig. 1A). CobS^Rc^ eluted unexpectedly early as a broad peak during size-exclusion chromatography (SEC; Fig. 1B), and negative-stain electron microscopy (EM) analysis revealed that CobS assembles into a variety of oligomers (Supplementary Fig. 1B), including ring-like structures commonly observed for other proteins of the AAA^+^ family ^26,41,42^.

To study CobS^Rc^ oligomeric states by cryo-EM, we collected five datasets (see Supplementary Fig. 2, Supplementary Fig. 3, Supplementary Fig. 8 and Supplementary Fig. 9) and solved four conformations: flat ring (3.8 Å), left-handed spiral ring (3.6 Å), left-handed helical filament (3.9 Å, helical rise 10.11 Å, helical twist -61.78°) and double ring (4.1 Å) (Fig. 1). Notably, no nucleotide was observed, indicating that interaction between the CobS^Rc^ protomers does not require the presence of ATP. Interestingly, the CobS^Rc^ hexameric flat ring (conformation A) lacks six-fold or helical symmetry. Instead, it forms a planar “double seam” dimer of CobS^Rc^ trimers with an imperfect C2-symmetry axis crossing the interface of the AAA^+^ large and small domains within the first (P1) and fourth (P4) subunits (Fig. 1C). In this conformation, the small AAA^+^ domains in P1 and P4 are slightly rotated (∼62°, Fig. 2D and Supplementary Movie 1) relative to the small domains of the remaining subunits, allowing the hexamer to maintain a planar configuration (Fig. 1C, 2A). Such an arrangement has been previously reported for other members of the AAA^+^ family, including RavA ^43^ and ClpX ^44,45^. In contrast to the flat ring, the remaining CobS^Rc^ structures (conformations B, C, and D) adopt an open, spiraling subunit organization. In conformation B (spiral ring), the first (P1) and last (P6) subunits of the steeper left-handed spiral assemblies appear flexible, as demonstrated by the absence of the corresponding cryo-EM density (Fig. 1D). The observed filaments (conformation C) varied in length, which is likely concentration-dependent and may be a result of the continuous addition of free CobS^Rc^ subunits to the spiral ring or the stacking of multiple spiral rings (Fig. 1E). Finally, CobS^Rc^ conformation D (spiral double ring) consists of two left-handed spiral hexamers oriented in opposite directions, but remaining connected at an interface between their respective subunits P1 and P6 (Fig. 1F, 2C).

**Fig. 2.**
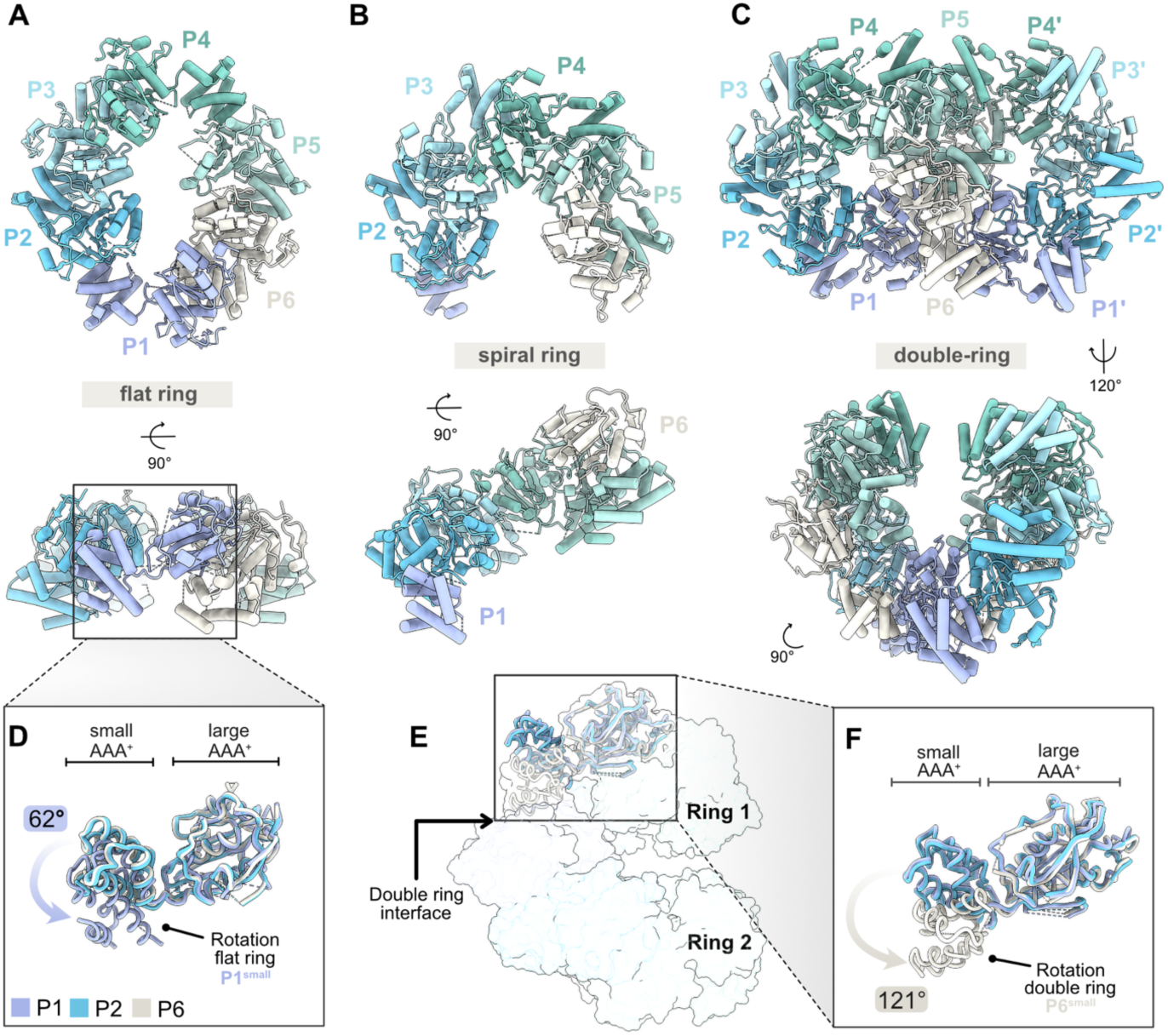
Conformational conversion of apo CobS^Rc^. **A.** Molecular model of the hexameric flat ring (conformation A) represented as cylinders/stubs, top view (top image) and side view (bottom image). CobS^Rc^ individual subunits colored as in Fig. 1 (P1 to P6). **B.** Molecular model of the spiral ring (conformation B), views and coloring as represented in **A. C.** Molecular model of the double ring (conformation D) viewed from two different angles, highlighting the interface between both left-handed spiral rings. Coloring as in **A. D.** Superposition of CobS^Rc^ subunits P1, P2 and P6 from the flat ring (conformation A) represented as licorice/ovals. Rotation of the P1 small AAA^+^ domain relative to the other subunits is highlighted. **E.** Semi-transparent surface representation of the double ring (conformation D) from **C.** displaying the superposition of subunits P1, P2 and P6 at the interface of the rings, represented as licorice/ovals. Colored as in **D.** for the flat ring subunits. **F.** Close-up of the superposition of CobS^Rc^ subunits P1, P2 and P6 from **E.** Rotation of the P6 small AAA^+^ domain relative to the other subunits is highlighted.

The conversion from the planar conformation A to helical conformation B likely results from movement and rotation of the small AAA^+^ domain relative to the large domain within the subunit P4. This conformational change appears to push the small and large AAA^+^ domains of neighboring subunits P5 and P3 in opposite directions. These rigid-body shifts lead to the rearrangement of the planar C2 into a spiral conformation, disrupting interactions between the large AAA^+^ domain of subunit P1 and the small AAA^+^ domain of P6 and destabilizing these domains (see Supplementary Movie 2). At the same time, a new interface is established between the large AAA^+^ domain of P4 and the large AAA^+^ domain of the neighboring P5 (helix 145-156 and PS1i; as present between P2-P3, P3-P4, P5-P6 and P6-P1), enabling the helical architecture of the hexamer (Fig. 2B). However, we cannot exclude the possibility of a reverse sequence of events leading to the transition from the spiral conformation B to the flat conformation A, where, upon flattening the spiral oligomer, an interface appears between the large AAA^+^ domain of P1 and the small AAA^+^ domain of P6, closing the ring (Fig. 2A, 2B). Interestingly, each pseudo-symmetric half of conformation A resembles the left-handed helical arrangement as observed in helical conformations B and C.

Similarly to the transition between conformations A and B, relative movements of the small and large AAA^+^ domains are required for the formation of the double ring conformation D (Fig. 1F, 2C). To accommodate such an unusual conformation for AAA^+^ proteins, the terminal subunit (P6) from each ring undergoes strong conformational rearrangement, in which its small AAA^+^ domain rotates 121° to interact with the large AAA^+^ domain of the first (P1) subunit of the opposing ring, resembling a twisted rope in the shape of the infinity sign (Fig. 1F, 2C – 2F Supplementary Movie 3). In this case, repositioning of the small AAA^+^ domain is more pronounced than during the transitions between conformations A and B (Fig. 2D, 2F). The interactions between all remaining subunits are equivalent to those observed between subunits in the spiral conformations B and C. Interestingly, in our reconstructions, the first ring appeared better resolved than the second ring (Supplementary Fig. 9A; compare the local resolutions of the first and second rings), suggesting substantial flexibility of the double ring CobS^Rc^ arrangement.

CobS^Rc^ exhibits an extremely dynamic conformational range, which cannot be attributed to the protein’s ATPase function, as all the described conformations were observed in the samples lacking added nucleotides. The conformational dynamics of free CobS^Rc^ may arise from spontaneous conformational rearrangements of the terminal subunits in helical conformations B and C; however, the influence of nucleotides present during protein overexpression cannot be excluded. Despite our attempts to prepare cryo-EM grids of the wild-type and the ATPase-impaired Walker B mutant (CobS^Rc-E138Q^) proteins in the presence of nucleotide and magnesium chloride (MgCl_2_), we observed only protein aggregation on the grid. We reason that the high ATPase activity of CobS^Rc^ (Supplementary Fig. 14), together with its high concentration in the sample and the absence of its native proteinaceous substrate (CobT), led to rapid conformational changes, potentially causing protein aggregation.

To obtain nucleotide-bound CobS^Rc^ structures and overcome protein aggregation, we prepared samples of CobS^Rc^ with ATP but without the addition of MgCl_2_, which prevented protein aggregation and allowed cryo-EM imaging (Supplementary Fig. 4 and Supplementary Fig. 10). From this data, we refined CobS^Rc^ in the spiral conformation B and double ring conformation D to 3.6 Å and 3.9 Å, respectively. In both conformations, we observed nucleotide densities in the respective nucleotide-binding pocket between the small and large AAA^+^ domains of subunits P2 to P5 in conformation B and P1 to P5 in conformation D. Owing to the higher flexibility of the double ring conformation D, we were unable to assign the nucleotides in the second ring because of the poor map resolution in this region (see Supplementary Fig. 10B).

### Structural characterization of the nucleotide-free CobST^Rc^ complex

Earlier negative-stain EM of *B. melitensis* CobS and CobT proteins (CobST^Bm^) suggested a larger two-tiered ring structure ^17^. Furthermore, in several functional studies on CobNST cobaltochelatase, the (VWA)-domain-containing adaptor CobT protein and CobS were co-purified ^20,47^. However, the absence of high-resolution structural information prevented an understanding of the molecular interplay between CobS and CobT and how ATP hydrolysis by CobS is coupled to mechanical work by CobT. To investigate the complex between the AAA^+^ CobS^Rc^ and its VWA partner protein CobT^Rc^ (Fig. 3B; hereafter CobST^Rc^), we co-expressed *R. capsulatus* proteins from a pETDuet-1 vector for protein co-purification using a double affinity pulldown approach (see Methods section). In brief, CobS^Rc^ was first captured by its C-terminal 6xHis-tag, followed by a Strep-tag purification to further capture the complex through an N-terminal tag on CobT^Rc^ (Fig. 3A), resulting in an approximate total protein concentration of 1 mg/mL in the elution fraction that was used for cryo-EM grid preparation. Interestingly, within the collected dataset (18,543 movies) for apo CobST^Rc^ (Supplementary Fig. 5), 2D class averages for the CobS^Rc^ double ring (conformation D) decorated by several undefined extra densities were predominant (Fig. 3C). As consensus refinement was insufficient to resolve the nature of the additional densities, we used focused 3D classification with a mask covering this peripheral region. The final 3.8 Å map showed a clear density that we could confidently assign as the CobT^Rc^ N-terminal domain (local resolution between 4.5-6 Å; Fig. 3D, Supplementary Fig.11A, 11C and 11D), attached to subunit P1’ of the CobS^Rc^ double ring. The CobT^Rc^ N-terminal helices α1, α2 and α5 display strong interactions with the small AAA^+^ domain helices α11 and α14 of CobS^Rc^ subunit P1’ on the second ring concave side (Fig. 3E, 3F). The interface appears to be mostly held by CobT^Rc^ residues Glu18, Thr23 and Arg63 with CobS^Rc^ residues Asn259, Thr253 and Arg308. However, no density was detected for the predicted flexible linker and VWA domain of CobT^Rc^ (Fig. 3B, Supplementary Fig. 15). Interestingly, we detected extra densities at the periphery of both CobS^Rc^ rings using a Gaussian-filtered map of the complex. We assigned these densities to five additional CobT^Rc^ N-terminal domains (see Supplementary Fig. 18B). However, the map resolution was insufficient for reliable model building. Interestingly, an additional density is wedged between subunits P1, P1’, P6 and P6’, precisely at the interface between both CobS^Rc^ rings (Supplementary Fig. 18C). We could not assign this density to either CobS^Rc^ or CobT^Rc^ N-terminal domain and speculate that it may originate from a different region of CobT^Rc^.

**Fig. 3.**
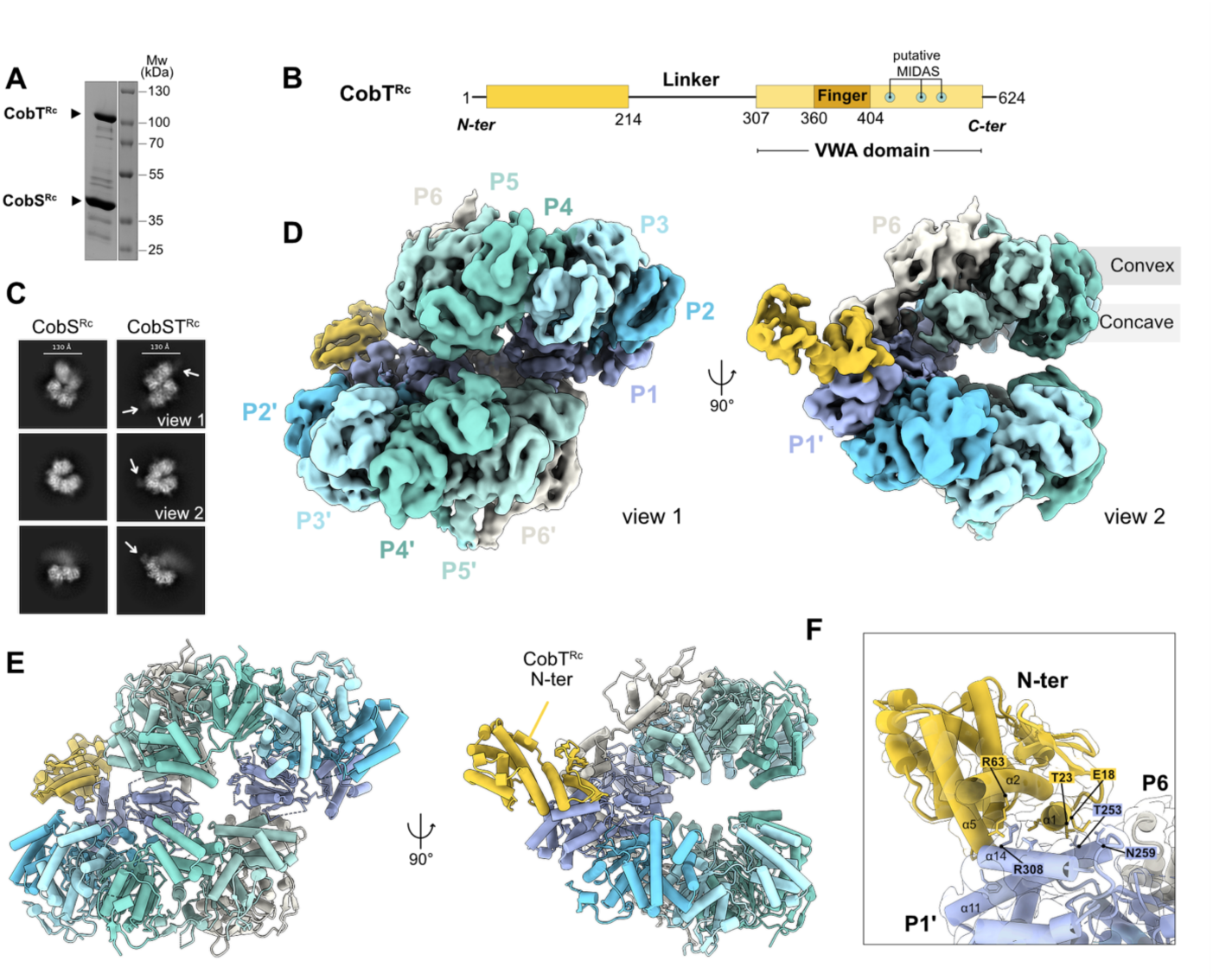
Cryo-EM structure of the nucleotide-free CobST^Rc^ complex. **A.** Sodium dodecyl sulfate-polyacrylamide gel electrophoresis (SDS-PAGE) analysis of the CobST^Rc^ Strep-tag elution fraction confirming the complex pull-down. **B.** Schematic 2D representation of CobT^Rc^ (based on AlphaFold ^46^ from Uniprot entry: D5AQ13) showing the protein’s different domains: N-terminal, predicted flexible linker and von Willebrand factor type A (VWA) domain with predicted finger region and putative Metal Ion-Dependent Adhesion Site (MIDAS: Asp410, Ser412, Ser414, Asn506 and Asp534; see Supplementary Fig. 15) motif. **C.** Representative 2D class averages of the CobS^Rc^ and CobST^Rc^ double ring showing extra density in the CobST^Rc^ sample (white arrows). **D.** Unsharpened cryo-EM map of CobST^Rc^ in the absence of nucleotides (global resolution: 3.8 Å, Supplementary Fig. 11C). Side view 1 (left) and side view 2 (right) as in the 2D class averages in **C. E.** Model of CobS^Rc^ in complex with one CobT^Rc^ N-terminal molecule (based on the local refinement map, see Supplementary Fig. 11D). Views as in **C. F.** Close-up of the interacting residues within α11 and α14 of the CobS^Rc^ P1’ small AAA^+^ domain and α1, α2 and α5 of CobT^Rc^ N-terminal (fitted within the local refinement map global resolution: 3.6 Å, see Supplementary Fig. 11D).

### The CobST^Rc^ complex under turnover conditions

To capture the active CobST^Rc^ complex under turnover conditions, we supplemented the purified CobST^Rc^ complex sample with 0.5 mM ATP:MgCl_2_ and immediately applied it to cryo-EM grids for vitrification. Comparable to the CobST^Rc^ apo dataset, we determined different left-handed ring conformations (Supplementary Fig. 6 and Supplementary Fig. 16), including nucleotide-bound double ring and several CobS^Rc^ spiral rings bound to a variable number of CobT^Rc^ (N-terminal) molecules (Supplementary Fig. 16B). Notably, in the 2D class averages, we identified a fraction of particles representing a single hexameric ring with extra density protruding from the ring’s central pore (Fig. 4A). Such an arrangement has been observed by EM analysis of other MoxR AAA^+^ proteins in complex with their VWA binding partners ^30,32,33^.

**Fig. 4.**
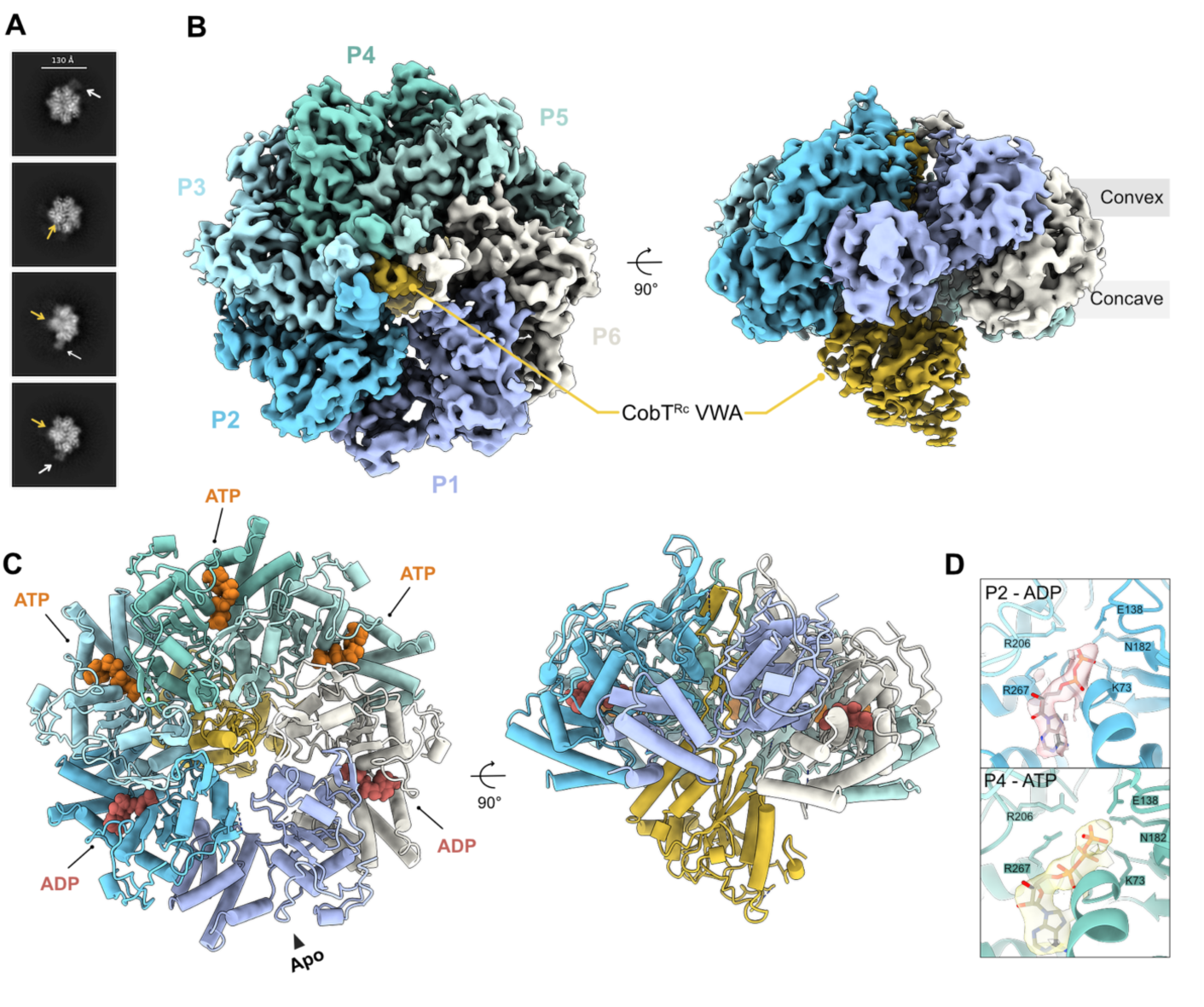
Cryo-EM structure of the active CobST^Rc^ complex. **A.** Representative 2D class averages of the CobS^Rc^ hexamer showing density for the CobT^Rc^ N-terminal (white arrows) and C-terminal (yellow arrows) domains. **B.** Cryo-EM map (sharpened) at global resolution of 3.3 Å of the active CobST^Rc^ (see Supplementary Fig. 11E), with only the CobT^Rc^ C-terminal residues His299 to Phe607 resolved. Top view (left) and side view (right) (threshold ∼0.03). **C.** CobST^Rc^ molecular model represented as cylinders/stubs, with the nucleotides bound to the CobS^Rc^ subunits shown as spheres. **D.** Representative nucleotide density for fitted ATP (P4) and ADP (P2) and the AAA^+^ residues involved in the ATP hydrolysis reaction (threshold ∼0.429, density highlighted around the nucleotide with a range of 2 Å).

To counter the apparent preferred orientation during 3D reconstruction, we applied several rounds of ab initio and heterogeneous refinement (Supplementary Fig. 6) and obtained a final map at a global resolution of ∼3.3 Å (∼46 K particles; Fig. 4B, Supplementary Fig. 11E and Supplementary Fig. 11G). In contrast to other CobS^Rc^ left-handed spiral conformations, the hexamer resembles an asymmetric closed ring, where four of the AAA^+^ subunits adopt a spiral right-handed arrangement (P2 to P5), while the remaining two (P1 and P6) stay in a more planar conformation, allowing the ring to fully close (see Supplementary Fig. 17). In this conformation, a globular density is seen through the AAA^+^ ring’s concave side, corresponding to the previously absent CobT^Rc^ C-terminal VWA domain (Fig. 4B, 4C). Contrary to the CobST^Rc^ conformations described earlier (Fig. 3, Supplementary Fig. 5), the CobT^Rc^ N-terminal domain was not apparent in this map. However, after Gaussian-filtering the active CobST^Rc^ map, we observed six repetitive densities interacting with the small AAA^+^ domains of CobS^Rc^, matching the CobT^Rc^ N-terminal domains previously identified in the CobST^Rc^ apo maps (Supplementary Fig. 18B).

To further resolve a possible conformational heterogeneity of CobT^Rc^ within our active CobST^Rc^ consensus map, we 3D classified the particles (∼46 K) into four different classes using a focused mask around the CobT^Rc^ region (see Supplementary Fig. 6 and Supplementary Fig. 19A). The obtained maps ranged from a global resolution of 3.7 to 4 Å and displayed different occupancies for the CobT^Rc^ domains (Supplementary Fig. 19B). While class 0 (∼15 K particles, 34.4%), class 1 (∼11 K particles, 24.5%), and class 3 (∼8 K particles, 17.5%) showed a partial resolved CobT^Rc^ C-terminal VWA as the consensus map, class 2 (10 K particles, 23.5%) also demonstrated clear density for N-terminal CobT^Rc^ domain interacting with CobS^Rc^ subunit P6 small domain. Although we did not observe substantial differences in the position of the VWA domain between the classes, the different occupancy suggests flexibility of CobT^Rc^ within the active CobST^Rc^ complex under turnover conditions.

We could confidently assign an ATP density at the Walker A motif of the right-handed spiraling CobS^Rc^ subunits P3, P4 and P5, and ADP to subunits P2 and P6 (Fig. 4C, 4D). In our consensus map, subunit P1 lacks nucleotide density, is more flexible, and is shifted relative to the other subunits, generating a “seam”. Such a state is in agreement with the hand-over-hand mechanism for substrate engagement and translocation suggested for other members of the AAA^+^ family ^41,48–51^. Furthermore, ADP-bound and apo subunits are less well-resolved, indicating an ATP-hydrolysis-induced flexibility of the ring.

The CobT^Rc^ VWA globular domain is oriented towards CobS^Rc^ ATP-bound subunits, while a finger region (approximately residues Ala362 to Thr403) extends from the globular VWA domain through the AAA^+^ pore (Fig. 4B, 4C, Fig. 5). At the top of the concave side of the CobS^Rc^ ring, this finger region appears to be in close contact with subunits P3 and P4 (Fig. 5A, panel 1). Inside the AAA^+^ pore, the finger loops twist and turn (Fig. 5A, panels 2, and 3), until reaching the end of the pore channel (Fig. 5A, panel 4). The CobT^Rc^ finger tip forms a hydrophobic α-helix (α5, residues Leu379 to Ala384) that interacts with the CobS^Rc^ H2i β-hairpin inserts, especially through residues Ile108 and Glu115 (P3 and P4) (Fig. 5B, 5C). The VWA domain of CobT^Rc^ also appears to maintain proximity to the P4 subunit post-sensor 1 loops region through the conserved residues Arg488 and Arg489 (α8; Fig. 5B, 5D and Supplementary Fig. 20). The proximity of CobT^Rc^ to subunits P2, P3 and P4 stabilizes their previously unresolved AAA^+^ post-sensor 1 loops (Fig. 5D). In the more distant and less stable AAA^+^ subunits P1, P5 and P6, these loops are mostly flexible and lack clear density for model building. The region of the CobT^Rc^ VWA domain located opposite to the finger and the CobS^Rc^ ring concave side, which is predicted to contain the MIDAS motif (Supplementary Fig. 15), was less resolved in our map, potentially due to its higher flexibility in the absence of its putative interaction partner CobN, precluding confident structure modelling and assignment of density corresponding to a metal ion density.

**Fig. 5.**
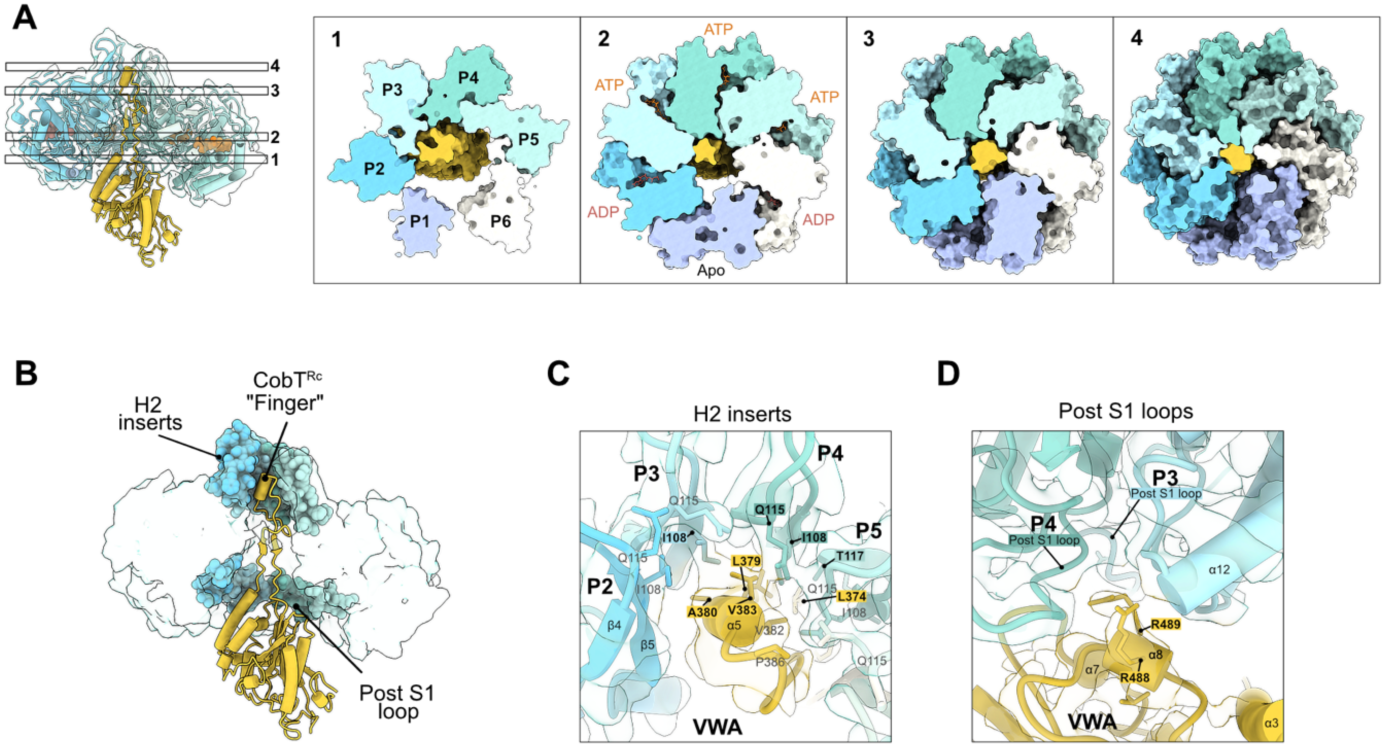
Overview of CobT^Rc^ C-terminal VWA domain interactions with CobS^Rc^. **A.** Top view slices (1 to 4) of the CobST^Rc^ model in surface representation, showing CobS^Rc^ subunit interactions with the CobT^Rc^ C-terminal finger region (colored gold) across its central pore. **B.** Side view slice of the CobST^Rc^ complex with CobS^Rc^ H2i and post-S1 loop insertions highlighted as spheres. CobT^Rc^ is represented as cylinders/stubs, and CobS^Rc^ as a molmap (generated in ChimeraX ^52^ software, resolution 6 Å). **C.** Close-up view of CobST^Rc^ finger tip, displaying CobT^Rc^ finger α-helix (residues Leu379 to Ala384) interactions with CobS^Rc^ H2i β-strands residues (Ile108, Gln115 and Thr117) within P3 to P5 subunits. **D.** Close-up view of CobS^RC^ P3 and P4 post-sensor 1 loops (residues Val184 to Ile199) and CobT^Rc^ VWA α-helix residues Arg488 and Arg489.

Overall, our structure for the active CobST^Rc^ complex is comparable to the cryo-EM structure of the *Paracoccus denitrificans* (*P*. *denitrificans*) Walker B NorQ mutant in complex with the VWA domain of NorD (PDB: 9QH6) ^33^. Intriguingly, while we observe the seam subunit flanked by ADP-bound AAA^+^ subunits (Fig. 4B, 4C), in the NorQD complex, the seam subunit is positioned between ATP- and ADP-bound subunits. Moreover, the *P*. *denitrificans* NorD protein displays an extended finger region with a longer α-helix that inclines towards the side of the ring (Supplementary Fig. 21).

### The CobST^Bm^ complex in the post-hydrolysis left-handed open state

To further gain insights into the cobaltochelatase complex and its reaction mechanism, we expressed and purified *B*. *melitensis* proteins CobN and CobST in *E*. *coli*. We set up an overnight reaction at 4 °C containing the three proteins in the presence of 2 mM ATP, 5 mM MgCl_2_, 0.5 mM Co^2+^, and 150 µM of the substrate hydrogenobyrinic acid *a*,*c*-diamide (HBAD; purified as in Widner *et*. *al*., 2016 ^53^). From this reaction, we collected a cryo-EM dataset that surprisingly did not reveal a fully assembled CobNST complex. Instead, most of the particles corresponded to a spiral CobS^Bm^ binding a variable number of CobT^Bm^ N-terminal molecules (Supplementary Fig. 7, Supplementary Fig. 13 and Fig. 6), as observed for CobST^Rc^. We obtained a final 3.2 Å map of CobST^Bm^ with five N-terminal CobT^Bm^ domains bound and one CobT^Bm^ VWA globular region facing the pore on the ring’s concave side (Fig. 6A, Supplementary Fig. 12). The interaction between CobT^Bm^ N-terminal and CobS^Bm^ small domain (Fig. 6D) is equivalent to the one observed for CobST^Rc^ (Fig. 3F), being held by a conserved arginine residue within CobT α2 (Supplementary Fig. 20). Due to the lack of attached CobN^Bm^ to the complex and the presence of ADP density within the Walker A motifs (P1 to P5, Fig. 6B, BC), we interpret this conformation as a post-reaction state, where the termination of the ATP hydrolysis cycle triggered the disassembly of the CobNST^Bm^ complex.

**Fig. 6.**
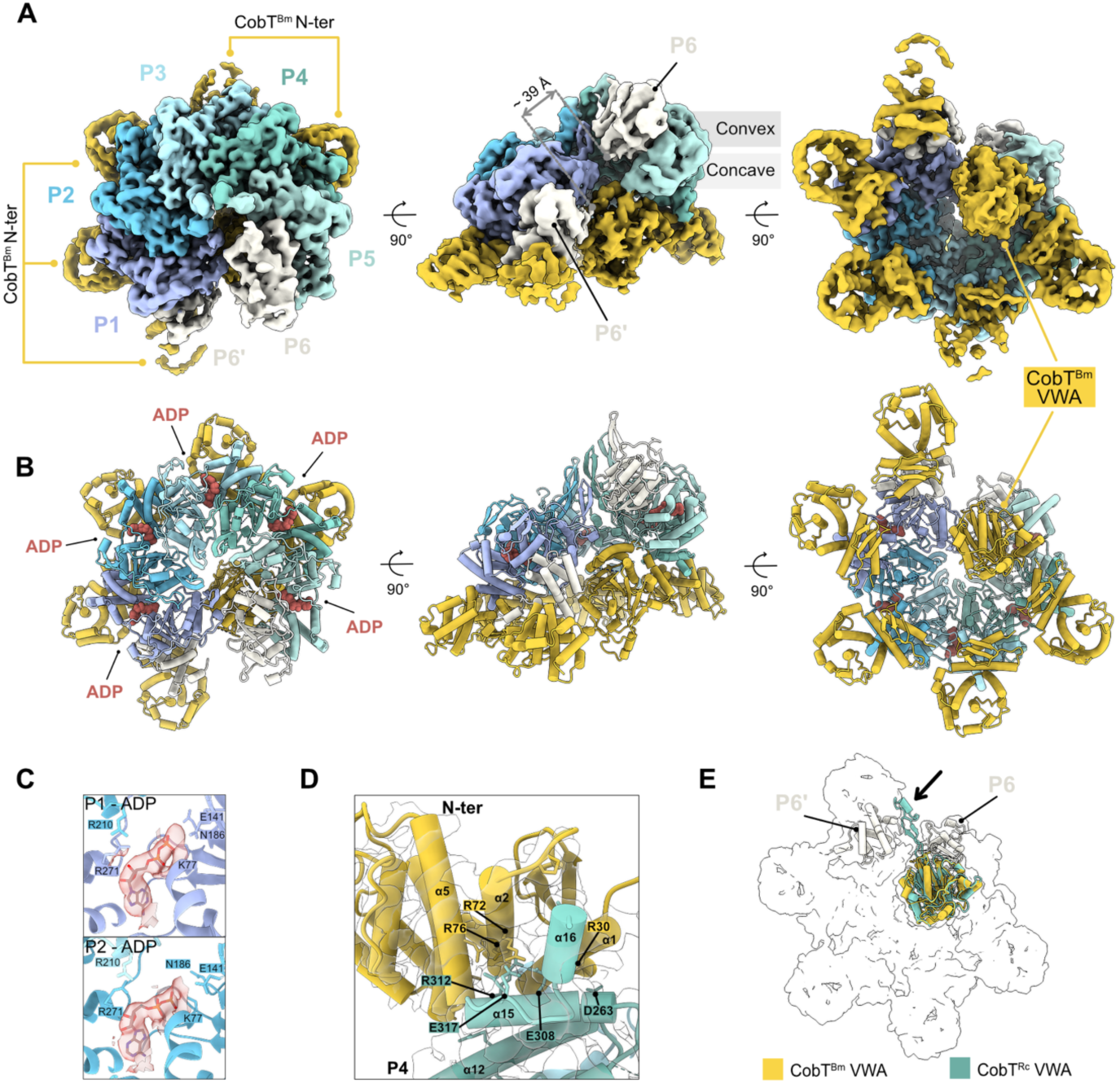
Cryo-EM structure of the post-hydrolysis CobST^Bm^ complex. **A.** Unsharpened cryo-EM map of the spiral left-handed CobS^Bm^ ring in complex with five molecules of CobT^Bm^ (Supplementary Fig. 12, global resolution 3.2 Å, threshold ∼ 0.0922). CobS^Bm^ subunits labelled and colored as in Fig.1. Extra small AAA^+^ domain highlighted as P6’. Top view (left), side view (middle) and bottom view (right). **B.** CobST^Bm^ molecular model represented as cylinders/stubs, with the nucleotides bound to the CobS^Rc^ subunits shown as spheres. Views as in **A. C.** Representative nucleotide density for fitted ADP (P1 and P2) and the AAA^+^ residues involved in the ATP hydrolysis reaction (sharpened map, threshold ∼0.217, density highlighted around the nucleotide with a range of 3 Å). **D.** Close-up of the interacting residues within P4 CobS^Bm^ small AAA^+^ domain helices α15 and α16 and CobT^Bm^ N-terminal helices α1, α2 and α5 (fitted within the sharpened map global resolution: 3.2 Å). **E.** Superimposition of CobT^Rc^ (from the active CobST^Rc^ structure) in CobT^Bm^ globular VWA domain, showing the possible orientation of the unresolved CobT^Bm^ finger region. The overall CobST^Bm^ model is represented as a transparent surface, highlighting the CobS^Bm^ helix terminal and subunits P6 and P6’.

In contrast to the hexameric closed ring observed for the nucleotide-bound active CobST^Rc^ (Fig. 5), the ADP-bound CobST^Bm^ complex adopts a steeper left-handed arrangement of the AAA^+^ subunits as observed for CobS^Rc^ spiral conformations. Moreover, to our surprise, CobS^Bm^ spiral ring shows a perfectly defined density for the subunit P1, which was not fully resolved in our structure of the spiral hexameric ring for the apo CobS^Rc^ (see Fig. 1D, conformation B). Additionally, we observed a density for an extra small AAA^+^ domain of a CobS^Bm^ molecule next to the P1 subunit (Fig. 6A, middle, P6’) that, being at a distance of approximately 39 Å from the large domain of the upper subunit P6 in the spiral ring, cannot be assigned as the P6 small AAA^+^ domain. In this post-hydrolysis state structure, the finger region of CobT^Bm^ VWA is displaced from the ring’s central pore and is not resolved in the map, probably due to its flexibility. Nevertheless, when superimposing the CobT^Rc^ from our active

CobST^Rc^ structure with CobT^Bm^ in this putative post-hydrolysis CobST^Bm^ complex, the finger region is oriented towards the opening of CobS^Bm^ spiral ring between the P6 and P6’ subunits’ large and small domains, respectively (see Fig. 6E). A morph generated by superimposing the active CobST^Rc^ and CobST^Bm^ post-hydrolysis structures suggests the conformational changes within the CobS ring drive its transition from a right-handed AAA^+^ subunit arrangement back to a left-handed one, rotating the VWA domain around the opening AAA^+^ pore and moving the finger region of CobT away from the pore toward the ring opening (Supplementary movie 4).

## Discussion

The ATP-dependent multi-subunit class I chelatases are fundamental to the biosynthesis of essential tetrapyrrole compounds, including chlorophyll and cobalamin (vitamin B_12_) ^5^. The function of this class of chelatases ultimately depends on the interplay between a large tetrapyrrole-binding catalytic subunit and an ATP-dependent motor unit, raising the question: why is an ATP motor unit required, and what mechanistic basis links energy from ATP hydrolysis to metal chelation? The inherent flexibility and transient nature of class I chelatases have hindered their structure determination and, therefore, a full mechanistic understanding. Based on our structural analysis of the *R*. *capsulatus* and *B*. *melitensis* motor component CobS, which was reconstituted with and without its binding partner CobT, and investigated in the absence of nucleotides or under turnover conditions, we propose a model for the assembly and activation of the CobST cobaltochelatase motor unit (Fig. 7).

**Fig. 7.**
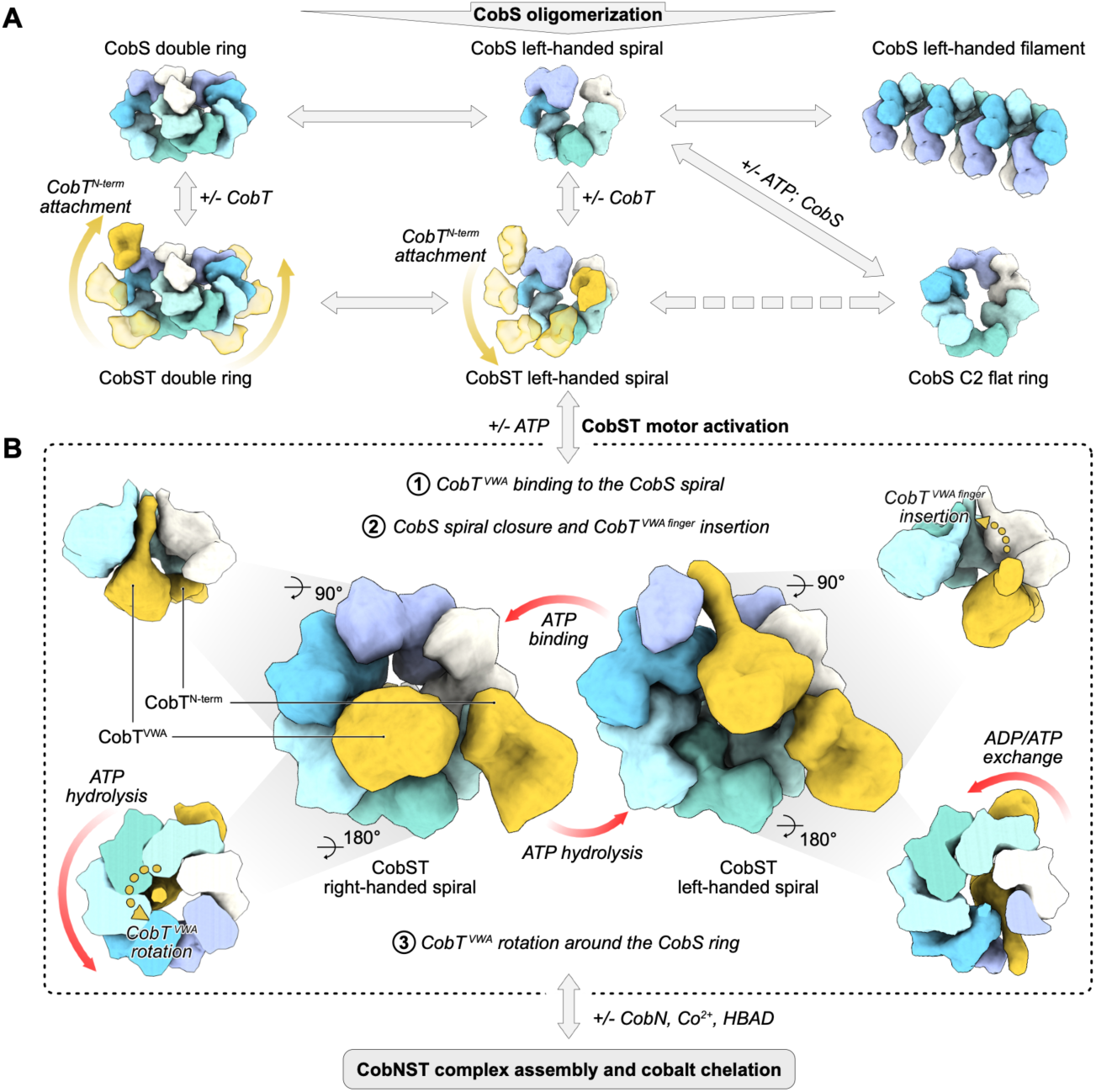
Proposed mechanism for the assembly and activation of the CobST motor unit of the multi-subunit cobaltochelatase CobNST. **A.** CobS oligomerizes independently of the presence of nucleotide into hexameric C2 flat rings and steeper left-handed spiral oligomers. The exchange between the different conformations is likely mediated by the dynamic AAA^+^ small domain. Multiple CobT units interact with CobS oligomers through CobT N-terminal domain, forming an inactive CobST complex represented by single- or double-ring left-handed spiral assemblies. **B.** ATP binding rearranges the left-handed spiral CobS subunits, shifting each subunit of CobS and simultaneously bringing the CobT VWA domain in proximity to the ring’s concave side. During subunit rearrangement, the CobT finger region is pushed into the central pore and held in place by CobS helix 2 and post-sensor 1 loop inserts. The repositioning of CobT and the concurrent closure of the ring to a right-handed semi-spiral conformation activates the cobaltochelatase motor component for the chelation reaction.

Our structural data on CobS show that the AAA^+^ assembles into different conformations, including flat C2 (conformation A), left-handed spiral (conformation B) hexamers and higher-order oligomers such as double spiral rings (conformation D) and short filaments (conformation C; Fig. 1 and Fig. 2). Similar conformations have been previously reported for other members of the AAA^+^ family. The C2 flat hexameric ring with two “seams” was previously observed with partially or fully occupied nucleotide-binding pockets for ClpX ^44,45^, RavA ^43^, and FtsH ^54^ proteins and has been suggested as a potential intermediate conformation of the proteins’ AAA^+^ cycle. Left-handed steeper spiral hexamers have been extensively observed for diverse AAA^+^ proteins, including the substrate-free, inactive state of the Lon AAA^+^ Protease ^49^. Moreover, filament formation was observed for AAA^+^ proteins ^55,56^ and, more recently, for the *Synechocysti*s sp. PCC 6803 magnesium chelatase ChlI AAA^+^ motor ^39^. Intriguingly, the observed CobS double ring appears to be a more unusual conformation, analogous to a previously reported cryo-EM structure of AAA^+^ RuvB in complex with dsDNA ^57^. The transitions between different observed CobS conformations are directly linked to the rotation of the small AAA^+^ domain of the protein (Fig. 2D, 2F), which could be driven by ATP hydrolysis ^44,45^. Interestingly, even in the absence of CobT, CobS retains ATPase activity (Supplementary Fig. 14), suggesting that precise regulation may be required to prevent futile ATP hydrolysis cycles *in vivo*. Several enzymes were reported to form filaments that serve as storage depots or regulate enzyme activity ^58^, suggesting a potential regulatory function of CobS filaments. However, further studies will be needed to test this hypothesis.

Based on our cryo-EM data for the nucleotide-free CobS in the presence of CobT, the N-terminal domain of CobT attaches to the periphery of the CobS left-handed spiral double rings (Fig. 3). The CobT C-terminal VWA domain remains detached and flexible, connected to the N-terminal domain only through the predicted long linker (Supplementary Fig. 15A). As cobalt chelation is likely dependent on VWA-mediated interaction of the CobST motor unit with the catalytic subunit CobN, these conformations are unlikely to drive the chelation reaction. When analyzing the nucleotide-free double-ring CobST cryo-EM map, we identified an unassigned density at the interface between the CobS rings, which may correspond to part of the CobT VWA domain or linker (Supplementary Fig. 18C). We speculate that the CobS double rings observed for the nucleotide-free CobST may represent an intermediate conformation that serves as a scaffold for proper CobT anchoring and positioning on CobS. Interestingly, it was recently shown that ChlD protein, the magnesium chelatase component analogous to CobT, promotes filament disassembly and subsequent activation of its AAA^+^ partner protein ChlI in the presence of ATP ^39^, suggesting that CobT may similarly promote disassembly of CobS oligomers.

Under turnover conditions, CobT and ATP cause a rearrangement of the CobS AAA^+^ ring from a steeper left-handed spiral double ring (Fig. 3) to a close spiral hexameric right-handed conformation (Fig. 4), which is reminiscent of other AAA^+^ complexes when engaging substrate binding and translocation ^49,59,60^. Our structure of the active CobST complex agrees with the arrangement of the VWA finger-threading through the AAA^+^ pore observed in the other MoxR-like AAA^+^/VWA complex NorQD (Supplementary Fig. 21) ^33^. Moreover, CobS structural inserts helix 2 β-hairpin insert (H2i) and pre-sensor 1 insert (PS1i) are essential for the AAA^+^ interaction with the CobT VWA partner protein (Fig. 5), as previously observed for the CbbQO ^30^ and NorQD complexes ^33^, suggesting a similar subunit interaction and mechanism for MoxR AAA+/VWA complexes. We propose that the CobT- and ATP-induced activation of CobS proceeds through three sequential steps. First, higher-order CobS oligomers disassemble and CobT N-terminal domain(s) attach to CobS left-handed spiral complexes. Second, the CobT VWA finger region is repositioned to interact with the H2i inserts within the central pore of the AAA^+^ hexamer, coupled to nucleotide-induced rearrangement of the CobS subunits. Third, the CobS ring closes into an active right-handed conformation, threading the CobT finger region through its central pore (Fig. 7B).

We propose that the ATP binding- and hydrolysis-induced shifts between CobS subunits are likely transmitted to the CobT finger (Supplementary Fig. 19, see results of 3D classification suggesting flexibility of the CobT VWA in the CobST complex under turnover conditions), which sequentially engages the H2i β-hairpin inserts of the ATP-bound CobS protomers. Similar mechanisms have been described for Lon and ClpX AAA^+^ proteins, which thread their substrates through the central pore during ATP hydrolysis ^48–50^. Likewise, stepwise ATP hydrolysis around the CobS AAA^+^ ring may result in displacement of the respective CobS protomer, accompanied by disengagement of the CobT finger and its subsequent handover to the H2i β-hairpin of the next ATP-bound CobS protomer. Further analogies to the mechanism suggested for NorQD ^32,33^ support that the sequential ATP hydrolysis around the AAA^+^ ring presumably results in a rotational movement of the CobT finger region, which is then transmitted to the MIDAS motif within the globular VWA domain. Although in other AAA^+^ proteins, ATP hydrolysis is proposed to proceed unidirectionally around the ring ^41^, we observed in our active CobST structure ADP molecules bound to the subunits on both sides of the seam subunit, challenging this model. A possible explanation can be the averaging of a heterogeneous particle population comprising different nucleotide-bound states, allowing for alternative nucleotide assignments to individual CobS subunits.

Interpolation between our active, right-handed CobST^Rc^ and post-hydrolysis, left-handed CobST^Bm^ structures supports an ATP-hydrolysis-dependent rotational movement of the CobT VWA globular domain (see Supplementary Movie 4). This rotation of the CobT VWA domain presumably transmits the ATP-hydrolysis-generated power stroke to the CobN catalytic subunit of cobaltochelatase (Supplementary Fig. 22), inducing the required conformational changes to converge HBAD and Co^2+^ within the CobN active site. We speculate that within the full active cobaltochelatase complex, the peripherally attached N-terminal domains of CobT (Supplementary Fig. 18, Fig. 6, Supplementary Fig. 22) may act as a scaffold by interacting with CobN and supporting its proper functional positioning. During the preparation of this manuscript, another study on the partial structure of *B*. *melitensis* CobNST became available ^61^. The authors’ findings further support the observed higher-order oligomers for CobS AAA^+^ and show that the C-terminal fragment of catalytic subunit CobN interacts with the CobT VWA and N-terminal domains within an AMP-PNP-trapped complex.

Finally, based on our post-hydrolysis structure of the CobST^Bm^ complex and 3D variability analysis (Supplementary Movie 5), we propose that the completion of ATP hydrolysis in all six CobS subunits results in opening of the CobS ring, its transition back to the left-handed spiral conformation, and release of the CobT VWA finger (Fig. 7B). This rearrangement presumably inactivates the motor of cobaltochelatase and primes it for a new reactivation upon binding of new ATP molecules.

While our attempts to assemble the full CobNST cobaltochelatase were unsuccessful, potentially owing to its high activity and tendency to disassemble after completion of the chelation reaction, our structures enable us to propose a mechanism for the assembly and activation of the CobST motor unit and the transmission of the power stroke to CobN. Our data extends the understanding on the general mechanism governing chaperone functions of MoxR AAA^+^/VWA complexes and provides a foundation for the future characterization of the overall multi-subunit cobaltochelatase.

## Methods

### Cloning

#### Cloning of CobS^Rc^, CobST^Rc^, CobN^Bm^ and CobST^Bm^ (co-)expression constructs

Primers and double-stranded DNA fragments (dsDNA) used in this study were purchased from Biolegio or Invitrogen (UK) and BioCat GmbH, respectively, and are listed in Supplementary Table 1. All PCR amplifications were performed using Q5-High-Fidelity DNA Polymerase (NEB) and the cloning steps were carried out in *E. coli* JM109 (Promega) or DH5α. Final constructs were validated by Sanger sequencing (Microsynth or GATC Biotech, Germany).

For overproduction of CobS^Rc^, the *cobS* gene from *R. capsulatus* SB1003 (Uniprot entry: D5AQ14) was amplified without its stop codon and ligated into the NdeI and XhoI sites of pET-21a(+) (Novagen) using T4 DNA ligase (NEB, UK), generating the plasmid pET21a-*cobS*-6xHis. CobS^Rc^ mutants (CobS^Rc-E138Q^, CobS^Rc-K73A^) were generated by site-directed mutagenesis using the Q5-High-Fidelity DNA Polymerase (NEB) following the manufacturer’s instructions. For co-expression of *cobS^Rc^* and *cobT*^Rc^ (Uniprot entry: D5AQ13), the genes were amplified and cloned into pETDuet-1 (Novagen) using HiFi Gibson assembly master mix (NEB), generating plasmid pETDuet-1-*cobS*-6xHis-StrepII-*cobT.* The *cobS* gene was cloned into the NdeI/XhoI sites of multiple cloning site two (MCS II) such that it was in frame with the sequence encoding a C-terminal 3C cleavage site and a 6xHis-tag. The *cobT* gene was subsequently cloned into the NcoI/NotI sites of MCS I such that it was in frame with the sequence encoding an N-terminal Twin-Strep-tag and 3C cleavage site.

The *cobN^Bm^* (Uniprot entry: C0RJR8), *cobS^Bm^*(Uniprot entry: C0RFR3) and *cobT^Bm^* (Uniprot entry: C0RFR2) genes from *Brucella melitensis* were cloned as previously detailed in Lundqvist *et*. *al*., 2009 ^17^. For the *cobST^Bm^* co-expression construct, the *cobS^Bm^* gene fragment was digested with NdeI and BamHI from a pET14b-*cobS^Bm^* plasmid and ligated into a digested pET3a (Novagen) using T4 DNA ligase (Promega, UK). The *cobT^Bm^* gene was digested from pET14b-*cobT^Bm^*with XbaI and HindIII and ligated into pET3a-*cobS^Bm^* digested with SpeI and HindIII, creating the final pET3a-*cobS^Bm^*-his-*cobT^Bm^*construct. Both *cobT^Bm^* and *cobN^Bm^* were cloned with a sequence encoding an N-terminal thrombin cleavage site followed by a 6xHis-tag.

### Protein Expression and Purification

#### CobS^Rc^ and CobS^Rc^ mutants Purification

For CobS^Rc^ expression, pET21a-*cobS*-6xHis was transformed into competent *E. coli* strains BL21(DE3) or LOBSTR(DE3) cells. Transformed cells were grown in Luria-Bertani medium (LB) with 100 µg/mL carbenicillin at 37 °C with shaking until an OD_600nm_ of 0.6. Cultures were transferred to 16 °C and protein expression was induced with 0.4 mM Isopropyl-β-D-thiogalactopyranosid (IPTG) overnight. Cells were collected at 6000 *g* for 30 min at 4 °C, flash frozen in liquid nitrogen and stored at - 80 °C. For purification, pellets were thawed on ice and resuspended in 20 mM Tris-HCl pH 7.5, 500 mM NaCl, 5 mM imidazole, 10% glycerol (lysis buffer) supplemented with one tablet of cOmplete EDTA-free Protease Inhibitor Cocktail (Roche), 0.5 mg/mL lysozyme, 0.05 mg/mL DNase I and 5 mM MgCl_2_ and briefly incubated at 4 °C with rotation. Cells were then further sonicated (Branson Ultrasonics Sonifier SFX250) in an ice bath at 25% amplitude with 5 sec on / 15 sec off for 2 min. The soluble fraction was collected through centrifugation at 17000 *g* for 20 min at 4 °C and incubated with HisPur^TM^ Ni-NTA Resin (Thermo Fisher Scientific). The column beads were washed with 20mM Tris-HCl pH 7.5, 500 mM NaCl, 5 mM imidazole, 5% glycerol (wash 1) and 20 mM Tris-HCl pH 7.5, 500 mM NaCl, 20 mM Imidazole, 5% glycerol (wash 2). His-tagged CobS was eluted with 20 mM Tris-HCl pH 7.5, 500 mM NaCl, 250 mM imidazole, 5% glycerol and dialyzed overnight at 4 °C against 20 mM Tris-HCl pH 7.5, 200 mM NaCl. The dialyzed sample was concentrated using a Vivaspin® centrifugal concentrator (10 kDa MWCO; Sartorius) at 2164 *g* at 4 °C and then applied to a Superose 6 Increase 10/300 column (Cytiva) for further size-exclusion chromatography (SEC). Protein purity was assessed by sodium dodecyl sulfate-polyacrylamide gel electrophoresis (SDS-PAGE) and protein concentration was determined by absorbance at 280nm. Concentrated protein was directly used for other analyses or aliquoted and stored at -80 °C until further experiments.

#### CobST^Rc^ Purification

The co-expression of tagged CobS^Rc^ and CobT^Rc^ was achieved using pETDuet-1-*cobS*-6xHis-StrepII-*cobT* in *E. coli* strain BL21(DE3) and purified by double affinity purification. Protein expression was carried out as described above for CobS^Rc^. Co-expression cell pellets were thawed on ice and resuspended in 50 mM Tris-HCl pH 7.4, 300 mM NaCl, 20 mM imidazole (binding buffer) supplemented with one tablet of cOmplete EDTA-free Protease Inhibitor Cocktail (Roche) and 0.05 mg/mL DNase I. Cells were lysed by sonication for 3 min at 20% amplitude and with 5 sec on / 20 sec off pulses. The CobST^Rc^ complex was first captured through the CobS^Rc^ C-terminal 6x-His-tag by Ni-NTA affinity purification, similarly to CobS^Rc^ purification, with the exception that the beads were extensively washed with binding buffer (wash 1) and wash buffer (50 mM Tris-HCl pH 7.4, 300 mM NaCl, 50 mM imidazole) (wash 2) before elution with buffer containing 400 mM imidazole. Eluted fractions containing the complex were directly loaded to a gravity column containing Strep-Tactin XT 4Flow high-capacity resin (IBA) for the capture of the CobST^Rc^ complex through CobT^Rc^ N-terminal Twin-Strep-tag. The resin was washed with 50 mM Tris-HCl pH 7.4, 300 mM NaCl (CobST^Rc^ Buffer), and the complex was eluted in the same buffer containing 50 mM biotin. Fractions containing CobST^Rc^ were buffer exchanged to the same buffer without biotin during concentration using a Vivaspin® 500 centrifugal concentrator (10 kDa MWCO; Sartorius) at 4 °C, 8000 *g*. The final complex concentration was estimated by SDS-PAGE using bovine serum albumin (BSA) standards as controls. Concentrated protein was directly used for other analyses or aliquoted and stored at -80 °C until further experiments.

#### CobN^Bm^ and CobST^Bm^ Purification

For *B.* CobN^Bm^ and CobST^Bm^ expression, pET3a containing N-terminal-His-tagged *cobN^Bm^* or pET3a containing both *cobS^Bm^* and N-terminal-His-tagged *cobT^Bm^*was transformed into competent *E. coli* strain BL21 star (DE3) pLysS T7P DNAJ cells. Transformed cells were grown in LB supplemented with 50 μM carbenicillin at 37 °C with shaking until an OD_600nm_ of 0.6-0.7. Bacteria cultures were transferred to 18 °C and protein expression was induced with 0.4 mM IPTG overnight. For purification, cells were harvested via centrifugation at 6000 g for 15 min, and pellets were resuspended in 20mM Tris-HCl pH 8, 500 mM NaCl (lysis buffer *^Bm^*). Cells were then sonicated using a large sonication probe in an ice bath at 40% amplitude with 30 sec on / 30 sec off for a total of 8 minutes. The lysate was centrifuged at 16000 x g for 40 minutes at 4 °C to pellet cell debris and incubated with 3 mL Ni-NTA resin (Qiagen) for 60 minutes at 4 °C. The column beads were collected with a gravity-flow column and washed with lysis buffer *Bm* containing either 0 (wash 1), 10 (wash 2) or 25 mM imidazole (wash 3). His-tagged CobST^Bm^ was co-eluted with lysis buffer*^Bm^* with either 250 (elution 1) or 400 mM imidazole (elution 2). The eluted sample was concentrated using a centrifugal concentrator (50 kDa MWCO; Amicon) at 4000 *g* at 4 °C and then applied to a Superdex 200 column (Cytiva) equilibrated with 25 mM Tris-HCl pH 8.0, 100 mM NaCl at 4 °C. Protein purity was assessed by SDS-PAGE and protein concentrations determined by absorbance at 280nm, and the purified proteins were subjected to cryo-EM analysis. Concentrated protein was directly used for other analyses or aliquoted and stored at -80 °C until further experiments.

### ATPase Activity Assay

The ATPase activity of CobS^Rc^ and mutants was measured using an enzyme-coupled assay, with an ATP-regenerating system, quantifying NADH oxidation as a result of ATP hydrolysis^62^. Before the measurement, the reaction cocktail mixture (50 mM Tris-HCl pH 7.5, 12 mM MgSO_4_, 3 mM phosphoenolpyruvate, 0.1 mM EGTA, 0.05 mg/mL pyruvate kinase, 0.05 mg/mL lactate dehydrogenase, 0.3 mM NADH, 10 mM ATP) was shortly equilibrated at 37 °C. A volume of 190 µL reaction cocktail was distributed to a 96-well plate, and the reaction was started with the addition of 10 µL of a 0.025 mg/mL protein stock (0.25 µg of protein in the final reaction). Reactions were performed in triplicate, and the measurements were followed by the decrease of NADH absorbance at 340 nm (NAD^+^ generation) for 2 h using a microplate reader (Infinite 200 Pro, Tecan) pre-equilibrated at 37 °C. The ATP hydrolysis rates were calculated based on the rate of decrease in NADH absorbance within its linear range. CobS^Rc^ protein amount for each sample was estimated by SDS-PAGE using bovine serum albumin (BSA) standards as controls.

### Negative-Stain Electron Microscopy

A 3 µL volume of purified CobS^Rc^ (0.04 mg/mL) was applied to freshly glow-discharged (using a PELCO easiGlow) in-house plastic-supported carbon-coated Gilder G-400 Cu grids (EMS; Electron Microscopy Sciences) ^63^. After 30 sec to 1 min of incubation on the grid, the excess sample was blotted using filter paper, and the grid was immediately stained three times with a 2% (w/v) uranyl formate solution and left to dry. Data collection was carried out manually on a JEM-2100Plus transmission electron microscope (JEOL) operated at an acceleration voltage of 200 kV and equipped with an XAROSA CMOS 20-megapixel camera (EMSIS) at a nominal magnification of 30k (3.12 Å per pixel). Negative-stain EM data were processed in cryoSPARC v4 ^64^.

### Cryo-EM sample preparation and data acquisition

#### CobS^Rc^ and CobST^Rc^

For cryo-EM grid preparation, 3 µL of protein (concentration between 0.7-1 mg/mL) was applied to freshly glow-discharged (using a PELCO easiGlow, 15mA, 120 seconds) C-flat (R1.2/1.3 300 mesh) Cu grids (EMS; Electron Microscopy Sciences) and immediately plunge-frozen in liquid ethane using a Vitrobot Mark IV (Thermo Fisher Scientific) set to 4 °C and 100% humidity. Cryo-EM data acquisition (cryo-EM platform - CellNanOs, Osnabrück, Germany) was performed automatically using EPU software (Thermo Fisher Scientific) on a Glacios cryo-transmission electron microscope operated at 200 kV and equipped with a Selectris energy filter and a Falcon 4i detector (Thermo Fisher Scientific). All images were collected in electron event representation (EER) mode at 165k nominal magnification (0.68 Å per pixel) with a defocus range from -0.8 µm to -1.8 µm and an electron dose of 50 e^-^/Å2.

For the CobS^Rc^-ATP (no MgCl_2_) reaction, 1 mM ATP was incubated on ice for 15 min with CobS^Rc^ (at a final concentration of 0.75 mg/mL). For the CobST^Rc^ - ATP/MgCl_2_ reaction, 0.5mM of an ATP:MgCl_2_ mixture was added to the protein (final volume of 4 µL and CobST^Rc^ concentration of approximately 0.6 mg/mL) and 3 µL of the reaction sample was immediately applied to the grid for plunge-freezing.

#### CobNST^Bm^

For cryo-EM grid preparation, 3 µL of sample (2 mg/mL final protein concentration diluted from a mixture containing 2.2 mg CobN^Bm^, 9.9 mg CobST^Bm^, 5 mM MgCl_2_, 2 mM ATP, 150 µM HBAD, 0.5 mM Co^2+^) was applied to freshly glow-discharged (20 mA for 60 s using a GloQube glow discharge system) C-flat R 2/1 mesh Cu 300 holy carbon grids, incubated for 2 s, blotted for 2 s, and plunge-frozen in liquid ethane using a Vitrobot Mark IV (Thermo Fisher Scientific) set to 4 °C and 100% humidity. Cryo-EM data acquisition (cryo-EM Swedish National Facility, Stockholm, Sweden) was performed automatically using EPU software (Thermo Fisher Scientific) on a 300-kV Titan Krios cryo-transmission electron microscope (Thermo Fisher Scientific) at a nominal magnification of 105k, corresponding to a calibrated pixel size of 0.84 Å. Exposures of 2 seconds were dose-fractionated into 40 frames (with a dose rate of 17.2 electrons/pixel/sec, resulting in a total dose of 47.9 e^-^/Å^2^. Detailed data collection parameters can be found in Supplementary Table 3.

### Cryo-EM data processing

#### CobS^Rc^ and CobST^Rc^

Cryo-EM data were pre-processed in cryoSPARC Live and further processed in cryoSPARC v4 and v5 ^64^. The data processing workflows are summarized in Supplementary Fig. 2 to Supplementary Fig. 6. Briefly, the collected movies were subjected to patch motion correction (EER number of fractions = 50, EER upsampling factor 1), patch-based contrast transfer function (CTF) estimation, micrograph denoiser and later curated to a CTF cut-off of 5 Å. In the case of CobS^Rc^ apo data, six datasets were collected and pre-processed similarly and later combined into two main micrograph sets (set A (five datasets, sample concentration approximately between 0.9 -1 mg/mL): 19352 micrographs and set B (one dataset, sample concentration approximately 0.75 mg/mL): 7843 micrographs). For the remaining samples, only a single dataset per sample was collected (see Supplementary Fig. 4 to Supplementary Fig. 6 for more details). Particles were picked using blob and template pickers, merged, and duplicated particles were removed. Remaining particles were initially extracted with a box size of 432 pix binned 4x (Fourier-cropped, resulting pixel size 2.72 Å) and further cleaned up in consecutive two-dimensional (2D) classification rounds to remove junk or wrong particle images. The best 2D classes were selected for ab initio reconstructions with multiple classes. For CobS^Rc^ Apo set A, 2D classes corresponding to filament particles were selected for subsequent filament tracing, resulting in an increased number of picked filament particles that were further cleaned by 2D and 3D classifications and used for further Non-uniform (NU)-refinement ^65^ or Helical refinement ^66^ (Supplementary Fig. 2).

The particles corresponding to the remaining (non-filament) conformations of CobS^Rc^ were processed similarly with slight variations (see Supplementary Fig. 2 for more details). Briefly, pre-selected particles were used for further rounds of 2D classifications and multiple-class ab initio 3D reconstructions, followed by Heterogeneous and NU-refinements, generating a polished group of particles for each conformation. All polished particle sets for each of the conformations were re-extracted using a full box size (432 pix or 450 pix, pixel size 293.76 Å and 306 Å, respectively) and used for final rounds of NU-refinement, local refinement, reference-based motion correction (RBMC) and global or local CTF refinement for further improvement of map quality.

The CobST^Rc^ datasets were processed similarly to the CobS^Rc^ non-filament conformations workflow with a few differences (see Supplementary Fig. 5 and Supplementary Fig. 6 for more details). For the CobST^Rc^ apo dataset, to separate particles corresponding to the CobS^Rc^ double ring decorated with extra densities from the CobS^Rc^ double ring alone, focused 3D classification (2 classes) was applied using a soft mask covering the best-resolved of these densities (CobT^Rc^ N-terminal), followed by local refinement to improve this region’s resolution further. For the CobST^Rc^ under turnover conditions dataset (ATP/MgCl_2_), we subjected the final consensus map particles (∼46 K; see Supplementary Fig. 6) to focused 3D classification with a mask around both the C-terminal and N-terminal (visible in a Gaussian-filtered map; see Supplementary Fig. 19).

#### Cob(N)ST^Bm^

Motion correction was performed using PatchMotion and contrast transfer function (CTF) parameters were calculated from averaged movies using PatchCTF within cryoSPARC v4 ^64^. Initial particles were picked using unbiased blob picking. The number of particles was then reduced by 2D and 3D classifications. Initial maps were calculated *ab initio* and the final maps were refined using non-uniform refinement ^65^ without the application of symmetry (C1) using cryoSPARC v4 (Supplementary Fig. 7). Detailed image processing parameters can be found in Supplementary Table 3. A final 3.2 Å resolution density map was used for *de novo* model building of *B. melitensis* CobST. To identify a possible discrete heterogeneity within the consensus map obtained for CobST^Bm^, 3D Variability analysis ^67^ was performed using a filter resolution of 4 Å, and solving for three components.

### Model building and refinement

Model building of *R*. *capsulatus* proteins was performed using AlphaFold 3 (AF) ^46^ predictions of monomeric CobS^Rc^ and CobT^Rc^ (Uniprot entries: D5AQ14 (CobS^Rc^), D5AQ13 (CobT^Rc^)) as initial models. The AF models were rigid-body fitted into the cryo-EM densities using the “Fit in Map” tool in UCSF ChimeraX ^52^ and further combined and manually adjusted in Coot ^68^ and Isolde ^69^. For CobST^Bm^ (Uniprot entries: C0RFR3 (CobS^Bm^), C0RFR2 (CobT^Bm^)), an initial full-length CobST structure was generated using the AlphaFold web server ^46^. The top predicted model was initially fitted into the EM density using ChimeraX ^52^.

The CobS(T) models for *R*. *capsulatus* and *B*. *melitensis* were further refined through interactive rounds of real-space refinement ^70^ in Phenix ^71^ and subsequent manual adjustments in Coot ^68^. Final model validation was done using MolProbity ^72^ within Phenix ^71^. Maps and models were inspected, visualized and figures were generated in UCSF ChimeraX ^52^ or PyMOL (The PyMOL Molecular Graphics System, Version 3.0 Schrödinger, LLC).

## Data Availability

*R*. *capsulatus* and *B. melitensis* CobS and CobST maps and models have been deposited in the Electron Microscopy Data Bank and Protein Data Bank under accession codes EMD-xxxx and yyyy respectively.

## Acknowledgments

DS acknowledges the support by the German Research Foundation (DFG; Emmy Noether grant 537976353, SFB 1557), the Osnabrück University’s Central Research Pool, and the Department of Biology’s early-career fellowship at Osnabrück University. DS and AM thank Prof. Arne Moeller for his support, carefully reading the manuscript and providing access to the cryo-EM platform at the CellNanOs center at Osnabrück University (DFG grant INST190/196-1 FUGG). DS and AM further thank Kilian Schnelle and Mateusz Nowacki for computational and wet lab support, respectively. BW, EB, MH were supported by the Swedish Research Council (2021-03992) and the Knut and Alice Wallenberg Foundation (2023.201). AH acknowledges the support of a Royal Society University Research Fellowship (URF\R1\191548 and URF\R\241006). The authors also gratefully acknowledge the support of the Biotechnology and Biological Sciences Research Council (BBSRC); this research was funded by the BBSRC Institute Strategic Programme Food Microbiome and Health BB/X011054/1 and its constituent project BBS/E/F/000PR13630 as well as project grants BB/X001946/1 and BB/Y008456/1. RWP was supported by UKRI (BBSRC) under Grant BB/X001946/1.

## Author contributions

AM, AH, MH, MW and DS designed the study. AM, ED, and AB cloned the constructs. AM purified, biochemically analyzed and performed cryo-EM on the *R*. *capsulatus* CobS and CobST complexes. BW and EB purified and performed cryo-EM on the *B*. *melitensis* CobST complexes. ED provided HBAD. AM, BW, EB and DS analyzed and interpreted the structures with input from AH, RP, MH and MW. The supervision of the project was done by MH, MW and DS. Funding acquisition was done by AH, MH, MW and DS. AM and DS wrote the manuscript with input from all authors.

## References

(1) Sokolovskaya, O. M.; Shelton, A. N.; Taga, M. E. Sharing Vitamins: Cobamides Unveil Microbial Interactions. Science 2020, 369 (6499), eaba0165. 10.1126/science.aba0165

(2) Banerjee, R.; Ragsdale, S. W. The Many Faces of Vitamin B12 : Catalysis by Cobalamin-Dependent Enzymes. Annu. Rev. Biochem. 2003, 72 (1), 209–247. 10.1146/annurev.biochem.72.121801.161828

(3) Kieninger, C.; Deery, E.; Lawrence, A. D.; Podewitz, M.; Wurst, K.; Nemoto-Smith, E.; Widner, F. J.; Baker, J. A.; Jockusch, S.; Kreutz, C. R.; Liedl, K. R.; Gruber, K.; Warren, M. J.; Kräutler, B. The Hydrogenobyric Acid Structure Reveals the Corrin Ligand as an Entatic State Module Empowering B12 Cofactors for Catalysis. Angew Chem Int Ed 2019, 58 (31), 10756–10760. 10.1002/anie.201904713

(4) Kräutler, B. Biochemistry of B12-Cofactors in Human Metabolism. In Water Soluble Vitamins; Stanger, O. Ed.; Subcellular Biochemistry; Springer Netherlands: Dordrecht, 2012; Vol. 56, pp 323–346. 10.1007/978-94-007-2199-9_17

(5) Bryant, D. A.; Hunter, C. N.; Warren, M. J. Biosynthesis of the Modified Tetrapyrroles—the Pigments of Life. Journal of Biological Chemistry 2020, 295 (20), 6888–6925. 10.1074/jbc.REV120.006194

(6) Stabler, S. P.; Allen, R. H. VITAMIN B12 DEFICIENCY AS A WORLDWIDE PROBLEM. Annu. Rev. Nutr. 2004, 24 (1), 299–326. 10.1146/annurev.nutr.24.012003.132440

(7) Smith, A. D.; Warren, M. J.; Refsum, H. Vitamin B12. In Advances in Food and Nutrition Research; Elsevier, 2018; Vol. 83, pp 215–279. 10.1016/bs.afnr.2017.11.005

(8) Osman, D.; Cooke, A.; Young, T. R.; Deery, E.; Robinson, N. J.; Warren, M. J. The Requirement for Cobalt in Vitamin B12: A Paradigm for Protein Metalation. Biochimica et Biophysica Acta (BBA) - Molecular Cell Research 2021, 1868 (1), 118896. 10.1016/j.bbamcr.2020.118896

(9) Warren, M. J.; Raux, E.; Schubert, H. L.; Escalante-Semerena, J. C. The Biosynthesis of Adenosylcobalamin (Vitamin B12). Nat. Prod. Rep. 2002, 19 (4), 390–412. 10.1039/b108967f

(10) Raux, E.; Schubert, H. L.; Roper, J. M.; Wilson, K. S.; Warren, M. J. Vitamin B12: Insights into Biosynthesis’s Mount Improbable. Bioorganic Chemistry 1999, 27 (2), 100–118. 10.1006/bioo.1998.1125

(11) Romão, C. V.; Ladakis, D.; Lobo, S. A. L.; Carrondo, M. A.; Brindley, A. A.; Deery, E.; Matias, P. M.; Pickersgill, R. W.; Saraiva, L. M.; Warren, M. J. Evolution in a Family of Chelatases Facilitated by the Introduction of Active Site Asymmetry and Protein Oligomerization. Proc. Natl. Acad. Sci. U.S.A. 2011, 108 (1), 97–102. 10.1073/pnas.1014298108

(12) Leech, H. K.; Raux, E.; McLean, K. J.; Munro, A. W.; Robinson, N. J.; Borrelly, G. P. M.; Malten, M.; Jahn, D.; Rigby, S. E. J.; Heathcote, P.; Warren, M. J. Characterization of the Cobaltochelatase CbiXL. Journal of Biological Chemistry 2003, 278 (43), 41900–41907. 10.1074/jbc.M306112200

(13) Brindley, A. A.; Raux, E.; Leech, H. K.; Schubert, H. L.; Warren, M. J. A Story of Chelatase Evolution. Journal of Biological Chemistry 2003, 278 (25), 22388–22395. 10.1074/jbc.M302468200

(14) Schubert, H. L.; Raux, E.; Wilson, K. S.; Warren, M. J. Common Chelatase Design in the Branched Tetrapyrrole Pathways of Heme and Anaerobic Cobalamin Synthesis. Biochemistry 1999, 38 (33), 10660–10669. 10.1021/bi9906773

(15) Warren, M. J.; Deery, E. Vitamin B12 (Cobalamin) Biosynthesis in the Purple Bacteria. In The Purple Phototrophic Bacteria; Hunter, C. N., Daldal, F., Thurnauer, M. C., Beatty, J. T., Eds.; Govindjee, Series Ed.; Advances in Photosynthesis and Respiration; Springer Netherlands: Dordrecht, 2009; Vol. 28, pp 81–95. 10.1007/978-1-4020-8815-5_5

(16) McGoldrick’, H.; Deery, E.; Warren, M.; Heathcote, P. Cobalamin (Vitamin B J Biosynthesis in Rhodobacter Capsulatus. Biochemical Society Transactions 2002, 30.

(17) Lundqvist, J.; Elmlund, D.; Heldt, D.; Deery, E.; Söderberg, C. A. G.; Hansson, M.; Warren, M.; Al-Karadaghi, S. The AAA+ Motor Complex of Subunits CobS and CobT of Cobaltochelatase Visualized by Single Particle Electron Microscopy. Journal of Structural Biology 2009, 167 (3), 227–234. 10.1016/j.jsb.2009.06.013

(18) Debussche, L.; Couder, M.; Thibaut, D.; Cameron, B.; Crouzet, J.; Blanche, F. Assay, Purification, and Characterization of Cobaltochelatase, a Unique Complex Enzyme Catalyzing Cobalt Insertion in Hydrogenobyrinic Acid a,c-Diamide during Coenzyme B12 Biosynthesis in Pseudomonas Denitrificans. J Bacteriol 1992, 174 (22), 7445–7451. 10.1128/jb.174.22.7445-7451.1992

(19) Young, T. R.; Martini, M. A.; Foster, A. W.; Glasfeld, A.; Osman, D.; Morton, R. J.; Deery, E.; Warren, M. J.; Robinson, N. J. Calculating Metalation in Cells Reveals CobW Acquires CoII for Vitamin B12 Biosynthesis While Related Proteins Prefer ZnII. Nat Commun 2021, 12 (1), 1195. 10.1038/s41467-021-21479-8

(20) Young, T. R.; Deery, E.; Foster, A. W.; Martini, M. A.; Osman, D.; Warren, M. J.; Robinson, N. J. Two Distinct Thermodynamic Gradients for Cellular Metalation of Vitamin B12. JACS Au 2023, 3 (5), 1472–1483. 10.1021/jacsau.3c00119

(21) Hunter, C. N.; Morey-Burrows, F. S. The Structural Chemistry and Biosynthesis of Chlorophylls. RSC Chem. Biol. 2026, 7 (7), 1134–1166. 10.1039/D6CB00082G

(22) Zhang, J.; Yuan, H.; Wang, X.; Dai, H.; Zhang, M.; Liu, L. Crystal Structure of the Large Subunit of Cobaltochelatase from *MYCOBACTERIUM TUBERCULOSIS* . Proteins 2021, 89 (4), 462–467. 10.1002/prot.26023

(23) Adams, N. B. P.; Bisson, C.; Brindley, A. A.; Farmer, D. A.; Davison, P. A.; Reid, J. D.; Hunter, C. N. The Active Site of Magnesium Chelatase. Nat. Plants 2020, 6 (12), 1491– 1502. 10.1038/s41477-020-00806-9

(24) Chen, X.; Pu, H.; Fang, Y.; Wang, X.; Zhao, S.; Lin, Y.; Zhang, M.; Dai, H.-E.; Gong, W.; Liu, L. Crystal Structure of the Catalytic Subunit of Magnesium Chelatase. Nature Plants 2015, 1 (9), 15125. 10.1038/nplants.2015.125

(25) Lundqvist, J.; Elmlund, H.; Wulff, R. P.; Berglund, L.; Elmlund, D.; Emanuelsson, C.; Hebert, H.; Willows, R. D.; Hansson, M.; Lindahl, M.; Al-Karadaghi, S. ATP-Induced Conformational Dynamics in the AAA+ Motor Unit of Magnesium Chelatase. Structure 2010, 18 (3), 354–365. 10.1016/j.str.2010.01.001

(26) Jessop, M.; Felix, J.; Gutsche, I. AAA+ ATPases: Structural Insertions under the Magnifying Glass. Current Opinion in Structural Biology 2021, 66, 119–128. 10.1016/j.sbi.2020.10.027

(27) Snider, J.; Houry, W. A. MoxR AAA+ ATPases: A Novel Family of Molecular Chaperones? Journal of Structural Biology 2006, 156 (1), 200–209. 10.1016/j.jsb.2006.02.009

(28) Wong, K. S.; Houry, W. A. Novel Structural and Functional Insights into the MoxR Family of AAA+ ATPases. Journal of Structural Biology 2012, 179 (2), 211–221. 10.1016/j.jsb.2012.03.010

(29) Shvarev, D.; Scholz, A. I.; Moeller, A. Conformational Variability of Cyanobacterial ChlI, the AAA+ Motor of Magnesium Chelatase Involved in Chlorophyll Biosynthesis. mBio 2023, 14 (5), e01893–23. 10.1128/mbio.01893-23

(30) Tsai, Y.-C. C.; Ye, F.; Liew, L.; Liu, D.; Bhushan, S.; Gao, Y.-G.; Mueller-Cajar, O. Insights into the Mechanism and Regulation of the CbbQO-Type Rubisco Activase, a MoxR AAA+ ATPase. Proc. Natl. Acad. Sci. U.S.A. 2020, 117 (1), 381–387. 10.1073/pnas.1911123117

(31) Sutter, M.; Roberts, E. W.; Gonzalez, R. C.; Bates, C.; Dawoud, S.; Landry, K.; Cannon, G. C.; Heinhorst, S.; Kerfeld, C. A. Structural Characterization of a Newly Identified Component of α-Carboxysomes: The AAA+ Domain Protein CsoCbbQ. Sci Rep 2015, 5 (1), 16243. 10.1038/srep16243

(32) Kahle, M.; Appelgren, S.; Elofsson, A.; Carroni, M.; Ädelroth, P. Insights into the Structure-Function Relationship of the NorQ/NorD Chaperones from Paracoccus Denitrificans Reveal Shared Principles of Interacting MoxR AAA+/VWA Domain Proteins. BMC Biol 2023, 21 (1), 47. 10.1186/s12915-023-01546-w

(33) Kahle, M.; Appelgren, S.; König, F.; Carroni, M.; Ädelroth, P.; Wendler, P. NorQD AAA+ Complex Drives Metal Insertion by a Twisting Mechanism. Nat Commun 2026, 17 (1), 3032. 10.1038/s41467-026-71044-4

(34) Whittaker, C. A.; Hynes, R. O. Distribution and Evolution of von Willebrand/Integrin A Domains: Widely Dispersed Domains with Roles in Cell Adhesion and Elsewhere. MBoC 2002, 13 (10), 3369–3387. 10.1091/mbc.e02-05-0259

(35) Jensen, P. E.; Gibson, L. C. D.; Hunter, C. N. ATPase Activity Associated with the Magnesium-Protoporphyrin IX Chelatase Enzyme of Synechocystis PCC6803 : Evidence for ATP Hydrolysis during Mg2+ Insertion, and the MgATP-Dependent Interaction of the ChlI and ChlD Subunits. 1999.

(36) Jensen, P. E.; Gibson, L. C. D.; Henningsen, K. W.; Hunter, C. N. Expression of the chlI, chlD, and chlH Genes from the Cyanobacterium Synechocystis PCC6803 in Escherichia Coli and Demonstration That the Three Cognate Proteins Are Required for Magnesium-Protoporphyrin Chelatase Activity. Journal of Biological Chemistry 1996, 271 (28), 16662–16667. 10.1074/jbc.271.28.16662

(37) Reid, J. D.; Siebert, C. A.; Bullough, P. A.; Hunter, C. N. The ATPase Activity of the ChlI Subunit of Magnesium Chelatase and Formation of a Heptameric AAA^+^ Ring. Biochemistry 2003, 42 (22), 6912–6920. 10.1021/bi034082q

(38) Farmer, D. A.; Brindley, A. A.; Hitchcock, A.; Jackson, P. J.; Johnson, B.; Dickman, M. J.; Hunter, C. N.; Reid, J. D.; Adams, N. B. P. The ChlD Subunit Links the Motor and Porphyrin Binding Subunits of Magnesium Chelatase. Biochemical Journal 2019, 476 (13), 1875–1887. 10.1042/BCJ20190095

(39) Łata, N.; Hałys, Ł.; Sendorek, P.; Pintscher, S.; Indyka, P.; Rawski, M.; Gabruk, M. Filament Formation by ChlI Challenges the Current View of Magnesium Chelatase Architecture.

(40) Gao, Y.; Wang, Y.; Wang, X.; Liu, L. Hexameric Structure of the ATPase Motor Subunit of Magnesium Chelatase in Chlorophyll Biosynthesis. Protein Science 2020, 29 (4), 1026–1032. 10.1002/pro.3816

(41) Puchades, C.; Sandate, C. R.; Lander, G. C. The Molecular Principles Governing the Activity and Functional Diversity of AAA+ Proteins. Nat Rev Mol Cell Biol 2020, 21 (1), 43–58. 10.1038/s41580-019-0183-6

(42) Wendler, P.; Ciniawsky, S.; Kock, M.; Kube, S. Structure and Function of the AAA+ Nucleotide Binding Pocket. Biochimica et Biophysica Acta (BBA) - Molecular Cell Research 2012, 1823 (1), 2–14. 10.1016/j.bbamcr.2011.06.014

(43) Jessop, M.; Arragain, B.; Miras, R.; Fraudeau, A.; Huard, K.; Bacia-Verloop, M.; Catty, P.; Felix, J.; Malet, H.; Gutsche, I. Structural Insights into ATP Hydrolysis by the MoxR ATPase RavA and the LdcI-RavA Cage-like Complex. Commun Biol 2020, 3 (1), 46. 10.1038/s42003-020-0772-0

(44) Glynn, S. E.; Martin, A.; Nager, A. R.; Baker, T. A.; Sauer, R. T. Structures of Asymmetric ClpX Hexamers Reveal Nucleotide-Dependent Motions in a AAA+ Protein-Unfolding Machine. Cell 2009, 139 (4), 744–756. 10.1016/j.cell.2009.09.034

(45) Stinson, B. M.; Nager, A. R.; Glynn, S. E.; Schmitz, K. R.; Baker, T. A.; Sauer, R. T. Nucleotide Binding and Conformational Switching in the Hexameric Ring of a AAA+ Machine. Cell 2013, 153 (3), 628–639. 10.1016/j.cell.2013.03.029

(46) Abramson, J.; Adler, J.; Dunger, J.; Evans, R.; Green, T.; Pritzel, A.; Ronneberger, O.; Willmore, L.; Ballard, A. J.; Bambrick, J.; Bodenstein, S. W.; Evans, D. A.; Hung, C.-C.; O’Neill, M.; Reiman, D.; Tunyasuvunakool, K.; Wu, Z.; Žemgulytė, A.; Arvaniti, E.; Beattie, C.; Bertolli, O.; Bridgland, A.; Cherepanov, A.; Congreve, M.; Cowen-Rivers, A. I.; Cowie, A.; Figurnov, M.; Fuchs, F. B.; Gladman, H.; Jain, R.; Khan, Y. A.; Low, C. M. R.; Perlin, K.; Potapenko, A.; Savy, P.; Singh, S.; Stecula, A.; Thillaisundaram, A.; Tong, C.; Yakneen, S.; Zhong, E. D.; Zielinski, M.; Žídek, A.; Bapst, V.; Kohli, P.; Jaderberg, M.; Hassabis, D.; Jumper, J. M. Accurate Structure Prediction of Biomolecular Interactions with AlphaFold 3. Nature 2024, 630 (8016), 493–500. 10.1038/s41586-024-07487-w

(47) Fang, H.; Li, D.; Kang, J.; Jiang, P.; Sun, J.; Zhang, D. Metabolic Engineering of Escherichia Coli for de Novo Biosynthesis of Vitamin B12. Nat Commun 2018, 9 (1), 4917. 10.1038/s41467-018-07412-6

(48) Ripstein, Z. A.; Vahidi, S.; Houry, W. A.; Rubinstein, J. L.; Kay, L. E. A Processive Rotary Mechanism Couples Substrate Unfolding and Proteolysis in the ClpXP Degradation Machinery. eLife 2020, 9, e52158. 10.7554/eLife.52158

(49) Shin, M.; Puchades, C.; Asmita, A.; Puri, N.; Adjei, E.; Wiseman, R. L.; Karzai, A. W.; Lander, G. C. Structural Basis for Distinct Operational Modes and Protease Activation in AAA+ Protease Lon. Sci. Adv. 2020, 6 (21), eaba8404. 10.1126/sciadv.aba8404

(50) Fei, X.; Bell, T. A.; Jenni, S.; Stinson, B. M.; Baker, T. A.; Harrison, S. C.; Sauer, R. T. Structures of the ATP-Fueled ClpXP Proteolytic Machine Bound to Protein Substrate. eLife 2020, 9, e52774. 10.7554/eLife.52774

(51) Chen, W.; Yang, J.; Lander, G. C. Cryo-EM Structures of Human ClpXP Reveal Mechanisms of Assembly and Proteolytic Activation. Molecular Biology November 13, 2024. 10.1101/2024.11.12.623337

(52) Pettersen, E. F.; Goddard, T. D.; Huang, C. C.; Meng, E. C.; Couch, G. S.; Croll, T. I.; Morris, J. H.; Ferrin, T. E. UCSF CHIMERAX : Structure Visualization for Researchers, Educators, and Developers. Protein Science 2021, 30 (1), 70–82. 10.1002/pro.3943

(53) Widner, F. J.; Lawrence, A. D.; Deery, E.; Heldt, D.; Frank, S.; Gruber, K.; Wurst, K.; Warren, M. J.; Kräutler, B. Total Synthesis, Structure, and Biological Activity of Adenosylrhodibalamin, the Non-Natural Rhodium Homologue of Coenzyme B12. Angew Chem Int Ed 2016, 55 (37), 11281–11286. 10.1002/anie.201603738

(54) Bieniossek, C.; Schalch, T.; Bumann, M.; Meister, M.; Meier, R.; Baumann, U. The Molecular Architecture of the Metalloprotease FtsH. Proc. Natl. Acad. Sci. U.S.A. 2006, 103 (9), 3066–3071. 10.1073/pnas.0600031103

(55) Mizuno, N.; Dramićanin, M.; Mizuuchi, M.; Adam, J.; Wang, Y.; Han, Y.-W.; Yang, W.; Steven, A. C.; Mizuuchi, K.; Ramón-Maiques, S. MuB Is an AAA+ ATPase That Forms Helical Filaments to Control Target Selection for DNA Transposition. Proc. Natl. Acad. Sci. U.S.A. 2013, 110 (27). 10.1073/pnas.1309499110

(56) Demircioglu, F. E.; Zheng, W.; McQuown, A. J.; Maier, N. K.; Watson, N.; Cheeseman, I. M.; Denic, V.; Egelman, E. H.; Schwartz, T. U. The AAA + ATPase TorsinA Polymerizes into Hollow Helical Tubes with 8.5 Subunits per Turn. Nat Commun 2019, 10 (1), 3262. 10.1038/s41467-019-11194-w

(57) Rish, A. D.; Shen, Z.; Chen, Z.; Zhang, N.; Zheng, Q.; Fu, T.-M. Molecular Mechanisms of Holliday Junction Branch Migration Catalyzed by an Asymmetric RuvB Hexamer. Nat Commun 2023, 14 (1), 3549. 10.1038/s41467-023-39250-6

(58) Park, C. K.; Horton, N. C. Structures, Functions, and Mechanisms of Filament Forming Enzymes: A Renaissance of Enzyme Filamentation. Biophys Rev 2019, 11 (6), 927–994. 10.1007/s12551-019-00602-6

(59) Gates, S. N.; Yokom, A. L.; Lin, J.; Jackrel, M. E.; Rizo, A. N.; Kendsersky, N. M.; Buell, C. E.; Sweeny, E. A.; Mack, K. L.; Chuang, E.; Torrente, M. P.; Su, M.; Shorter, J.; Southworth, D. R. Ratchet-like Polypeptide Translocation Mechanism of the AAA+ Disaggregase Hsp104. Science 2017, 357 (6348), 273–279. 10.1126/science.aan1052

(60) De La Peña, A. H.; Goodall, E. A.; Gates, S. N.; Lander, G. C.; Martin, A. Substrate-Engaged 26 *S* Proteasome Structures Reveal Mechanisms for ATP-Hydrolysis–Driven Translocation. Science 2018, 362 (6418), eaav0725. 10.1126/science.aav0725

(61) Zhou, Y.; Yuan, H.; Wu, Y.; Wang, J.; Chen, H.; Yao, L.; Wang, M.; Wang, J.; He, C.; Chen, X.; Liu, L. Assembly of the ATP-Driven Cobalt Chelatase.

(62) Urbatsch, I. L.; Sankaran, B.; Weber, J.; Senior, A. E. P-Glycoprotein Is Stably Inhibited by Vanadate-Induced Trapping of Nucleotide at a Single Catalytic Site. Journal of Biological Chemistry 1995, 270 (33), 19383–19390. 10.1074/jbc.270.33.19383

(63) Januliene, D.; Moeller, A. Single-Particle Cryo-EM of Membrane Proteins. In Structure and Function of Membrane Proteins; Schmidt-Krey, I., Gumbart, J. C., Eds.; Methods in Molecular Biology; Springer US: New York, NY, 2021; Vol. 2302, pp 153–178. 10.1007/978-1-0716-1394-8_9

(64) Punjani, A.; Rubinstein, J. L.; Fleet, D. J.; Brubaker, M. A. cryoSPARC: Algorithms for Rapid Unsupervised Cryo-EM Structure Determination. Nat Methods 2017, 14 (3), 290–296. 10.1038/nmeth.4169

(65) Punjani, A.; Zhang, H.; Fleet, D. J. Non-Uniform Refinement: Adaptive Regularization Improves Single-Particle Cryo-EM Reconstruction. Nat Methods 2020, 17 (12), 1214–1221. 10.1038/s41592-020-00990-8

(66) Egelman, E. H. Reconstruction of Helical Filaments and Tubes. In Methods in Enzymology; Elsevier, 2010; Vol. 482, pp 167–183. 10.1016/S0076-6879(10)82006-3

(67) Punjani, A.; Fleet, D. J. 3D Variability Analysis: Resolving Continuous Flexibility and Discrete Heterogeneity from Single Particle Cryo-EM. Journal of Structural Biology 2021, 213 (2), 107702. 10.1016/j.jsb.2021.107702

(68) Emsley, P.; Lohkamp, B.; Scott, W. G.; Cowtan, K. Features and Development of Coot. Acta Crystallogr D Biol Crystallogr 2010, 66 (4), 486–501. 10.1107/S0907444910007493

(69) Croll, T. I. *ISOLDE* : A Physically Realistic Environment for Model Building into Low-Resolution Electron-Density Maps. Acta Crystallogr D Struct Biol 2018, 74 (6), 519–530. 10.1107/S2059798318002425

(70) Afonine, P. V.; Poon, B. K.; Read, R. J.; Sobolev, O. V.; Terwilliger, T. C.; Urzhumtsev, A.; Adams, P. D. Real-Space Refinement in Phenix for Cryo-EM and Crystallography.

(71) Liebschner, D.; Afonine, P. V.; Baker, M. L.; Bunkóczi, G.; Chen, V. B.; Croll, T. I.; Hintze, B.; Hung, L.-W.; Jain, S.; McCoy, A. J.; Moriarty, N. W.; Oeffner, R. D.; Poon, B. K.; Prisant, M. G.; Read, R. J.; Richardson, J. S.; Richardson, D. C.; Sammito, M. D.; Sobolev, O. V.; Stockwell, D. H.; Terwilliger, T. C.; Urzhumtsev, A. G.; Videau, L. L.; Williams, C. J.; Adams, P. D. Macromolecular Structure Determination Using X-Rays, Neutrons and Electrons: Recent Developments in Phenix. Acta Crystallogr D Struct Biol 2019, 75 (10), 861–877. 10.1107/S2059798319011471

(72) Davis, I. W.; Leaver-Fay, A.; Chen, V. B.; Block, J. N.; Kapral, G. J.; Wang, X.; Murray, L. W.; Arendall, W. B.; Snoeyink, J.; Richardson, J. S.; Richardson, D. C. MolProbity: All-Atom Contacts and Structure Validation for Proteins and Nucleic Acids. Nucleic Acids Research 2007, 35 (Web Server), W375–W383. 10.1093/nar/gkm216

