## Supplementary Data for "Assembly and activation of the AAA^+^ motor of the multi-subunit cobaltochelatase"

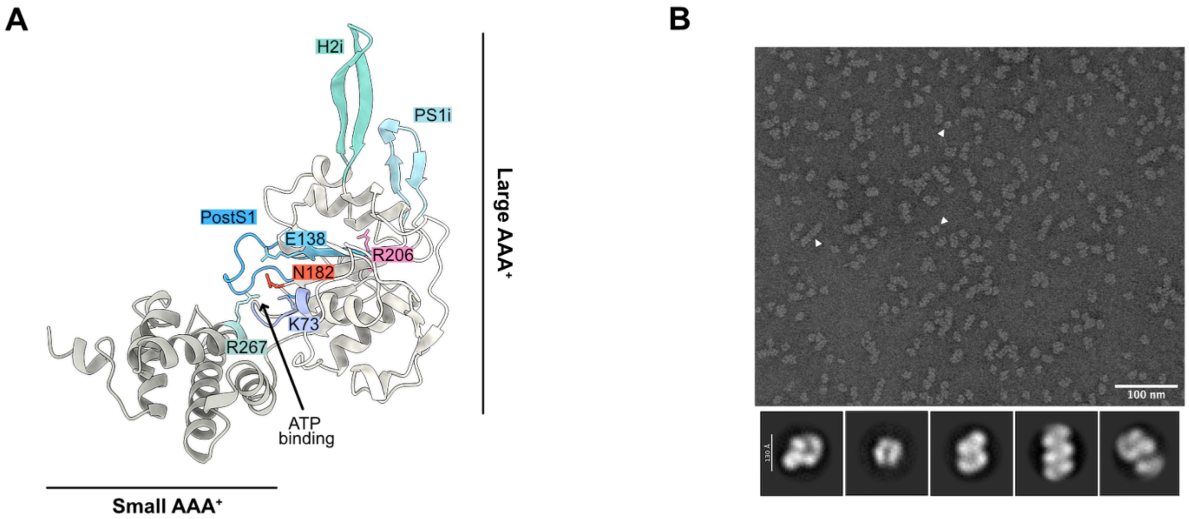


**Supplementary Fig. 1. Characterization of CobS$^{Rc}$. A.** AlphaFold [1] (Uniprot entry: D5AQ14) prediction of the CobS$^{Rc}$ AAA$^{+}$ ATPase domains (large and small AAA$^{+}$) and motifs. Highlighted motifs: Walker A (WA, Gly70 to Thr75), Walker B (WB, Ala133 to Glu138), sensor 1 (S1, Asn182), arginine finger (R-finger, Arg206) and sensor 2 (S2, Arg267). CobS$^{Rc}$ structural insertions: helix 2 insert (H2i, green), pre-sensor 1 insert (PS1i, light blue) and post-sensor 1 loop insert (Post S1, blue). **B.** Negative-stain EM micrograph and 2D class averages of purified CobS$^{Rc}$ showing different oligomers (white arrows).

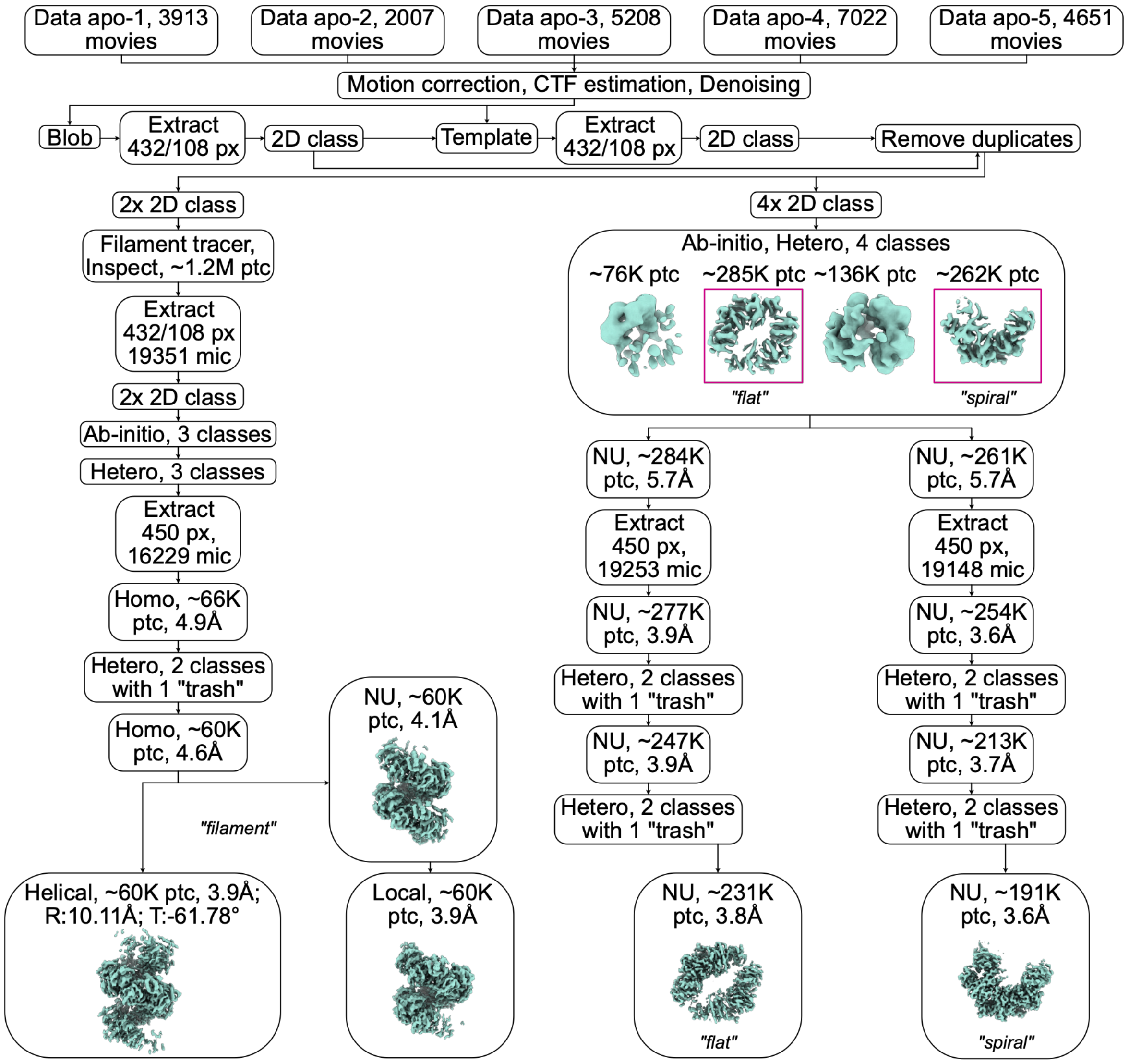


**Supplementary Fig 2. Cryo-EM processing workflow for the CobS$^{Rc}$ apo sample at 0.9-1 mg/ml.** For Extract jobs, box size (full/Fourier-cropped) in pixels (px) and micrograph (mic) number are indicated. For Hetero-, Homo-, NU-, Local-, and Helical refinement jobs, the approximate number of used particles (ptc) and achieved resolution are indicated. Purple frames indicate Hetero-refinement reconstructions used for downstream processing. CTF, contrast transfer function.

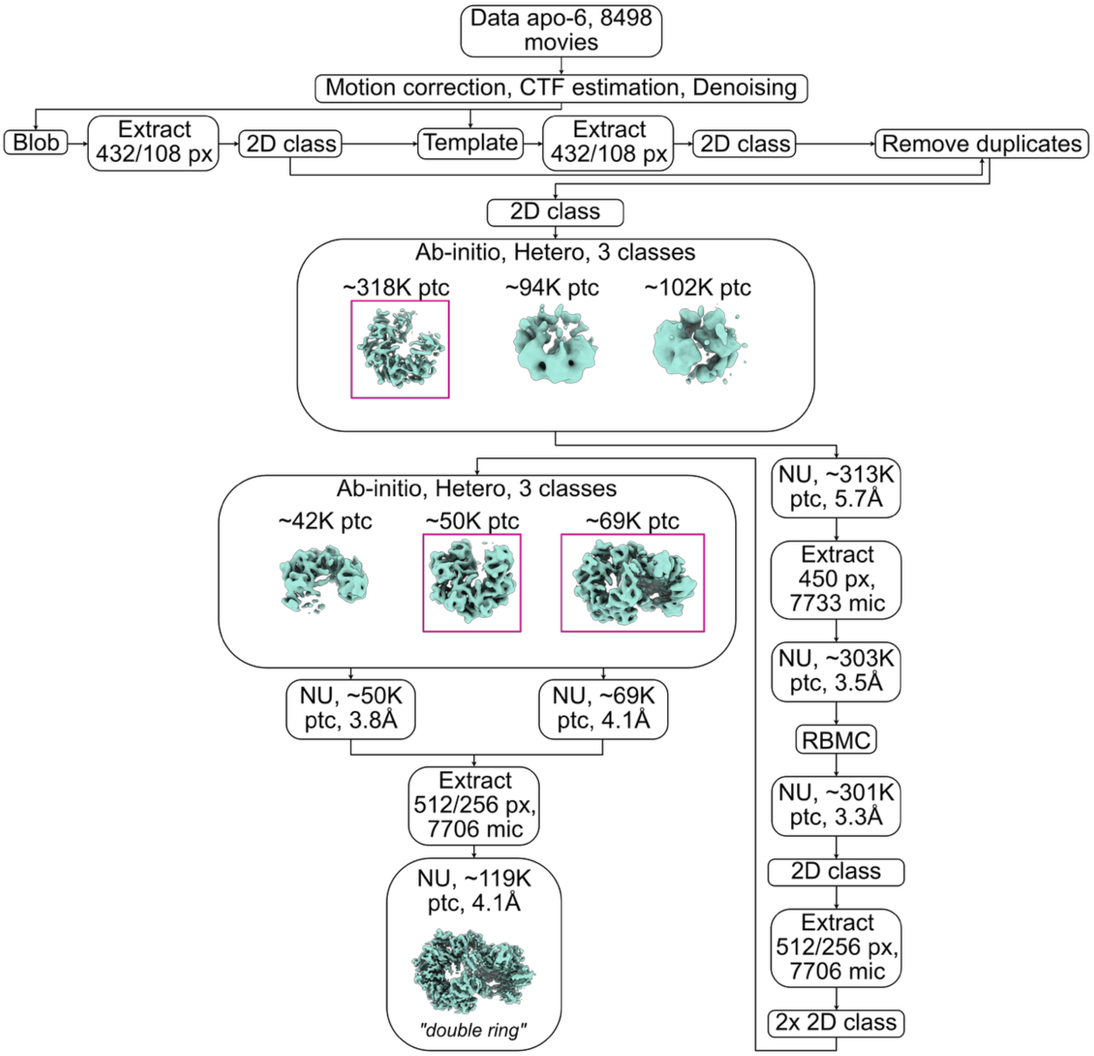


**Supplementary Fig. 3. Cryo-EM processing workflow for the $CobS^{Rc}$ apo sample at 0.75 mg/ml.** For Extract jobs, box size (full/Fourier-cropped) in pixels (px) and micrograph (mic) number are indicated. For Hetero- and NU-refinement jobs, the approximate number of used particles (ptc) and achieved resolution are indicated. Purple frames indicate Hetero-refinement reconstructions used for downstream processing. CTF, contrast transfer function; RBMC, reference-based motion correction.

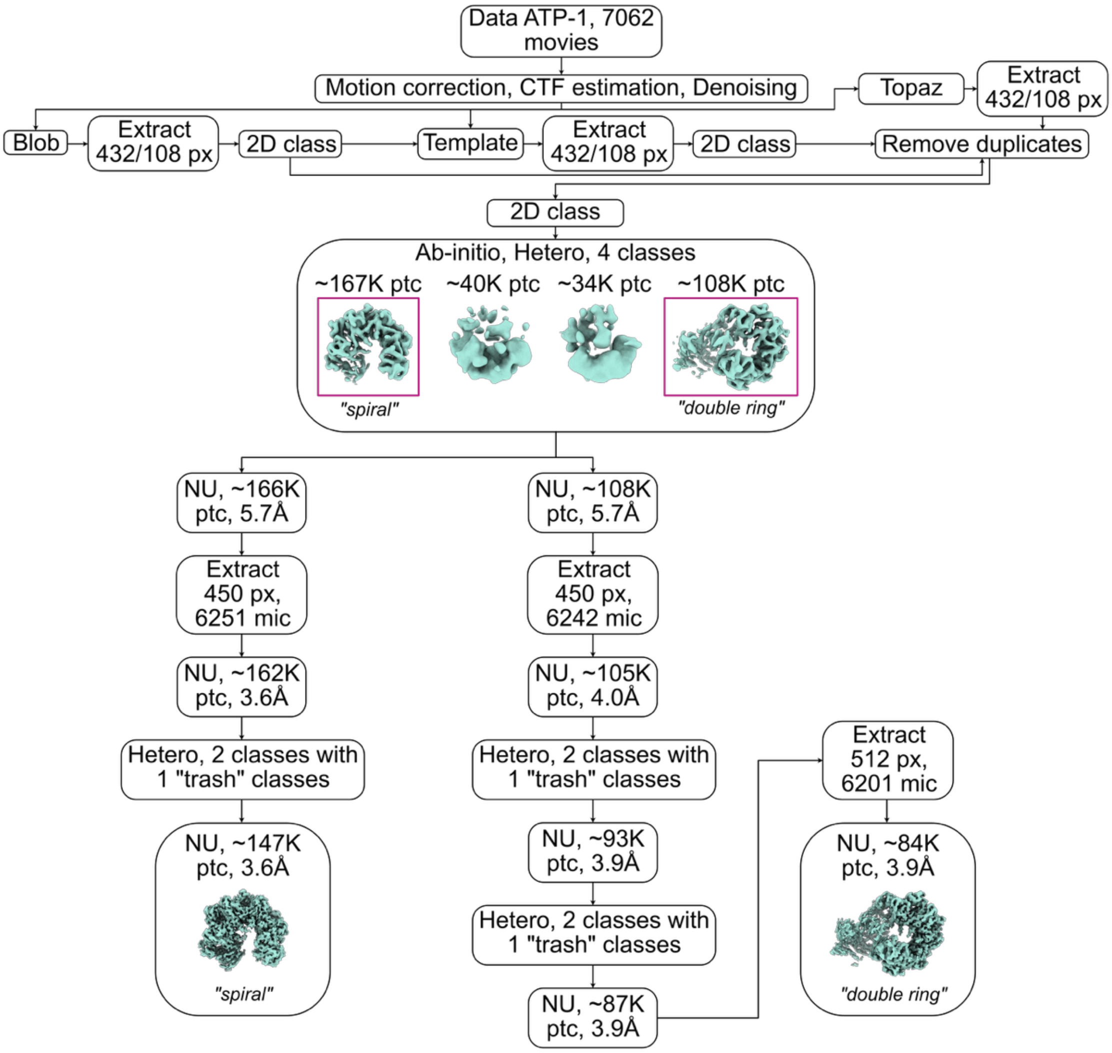


**Supplementary Fig. 4. Cryo-EM processing workflow for the CobS$^{Rc}$ sample incubated with ATP.** For Extract jobs, box size (full/Fourier-cropped) in pixels (px) and micrograph (mic) number are indicated. For Hetero- and NU-refinement jobs, the approximate number of used particles (ptc) and achieved resolution are indicated. Purple frames indicate Hetero-refinement reconstructions used for downstream processing. CTF, contrast transfer function.

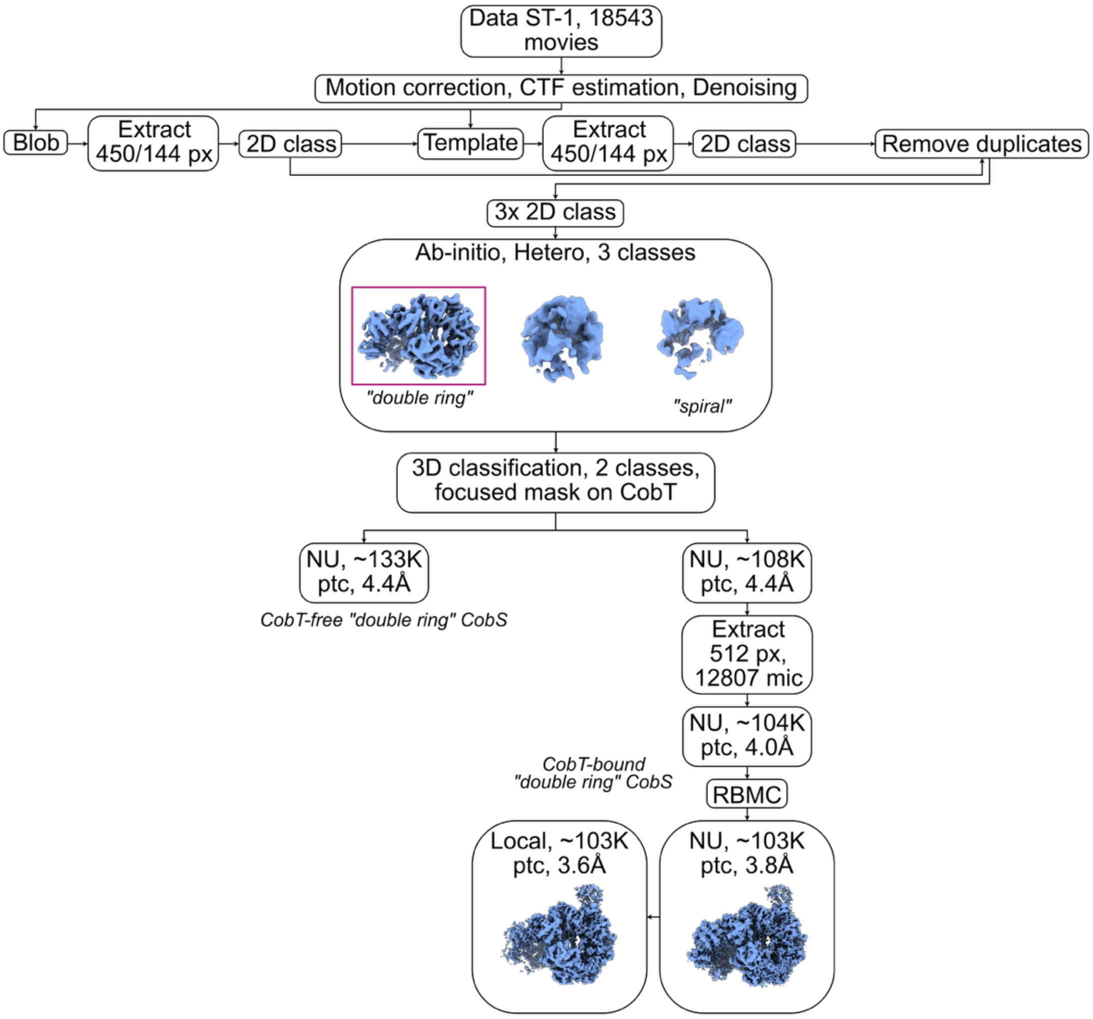


**Supplementary Fig. 5. Cryo-EM processing workflow for the CobST$^{Rc}$ apo sample.** For Extract jobs, box size (full/Fourier-cropped) in pixels (px) and micrograph (mic) number are indicated. For Hetero-, NU-, and Local-refinement jobs, the approximate number of used particles (ptc) and achieved resolution are indicated. Purple frames indicate Hetero-refinement reconstructions used for downstream processing. CTF, contrast transfer function; RBMC, reference-based motion correction.

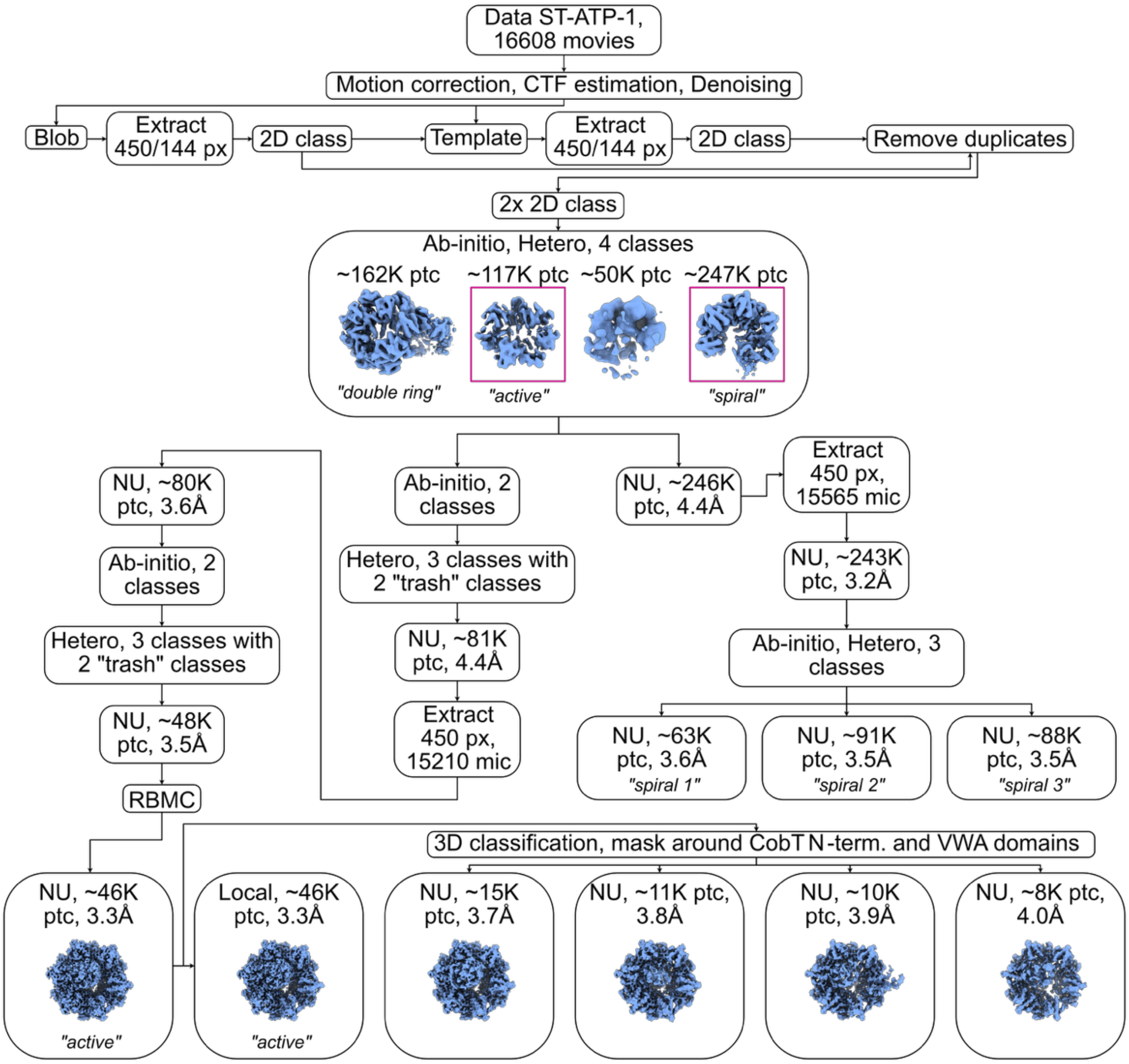


**Supplementary Fig. 6. Cryo-EM processing workflow for the $CobST^{Rc}$ sample incubated with ATP and $MgCl_2$.** For Extract jobs, box size (full/Fourier-cropped) in pixels (px) and micrograph (mic) number are indicated. For Hetero-, NU-, and Local-refinement jobs, the approximate number of used particles (ptc) and achieved resolution are indicated. Purple frames indicate Hetero-refinement reconstructions used for downstream processing. CTF, contrast transfer function; RBMC, reference-based motion correction.

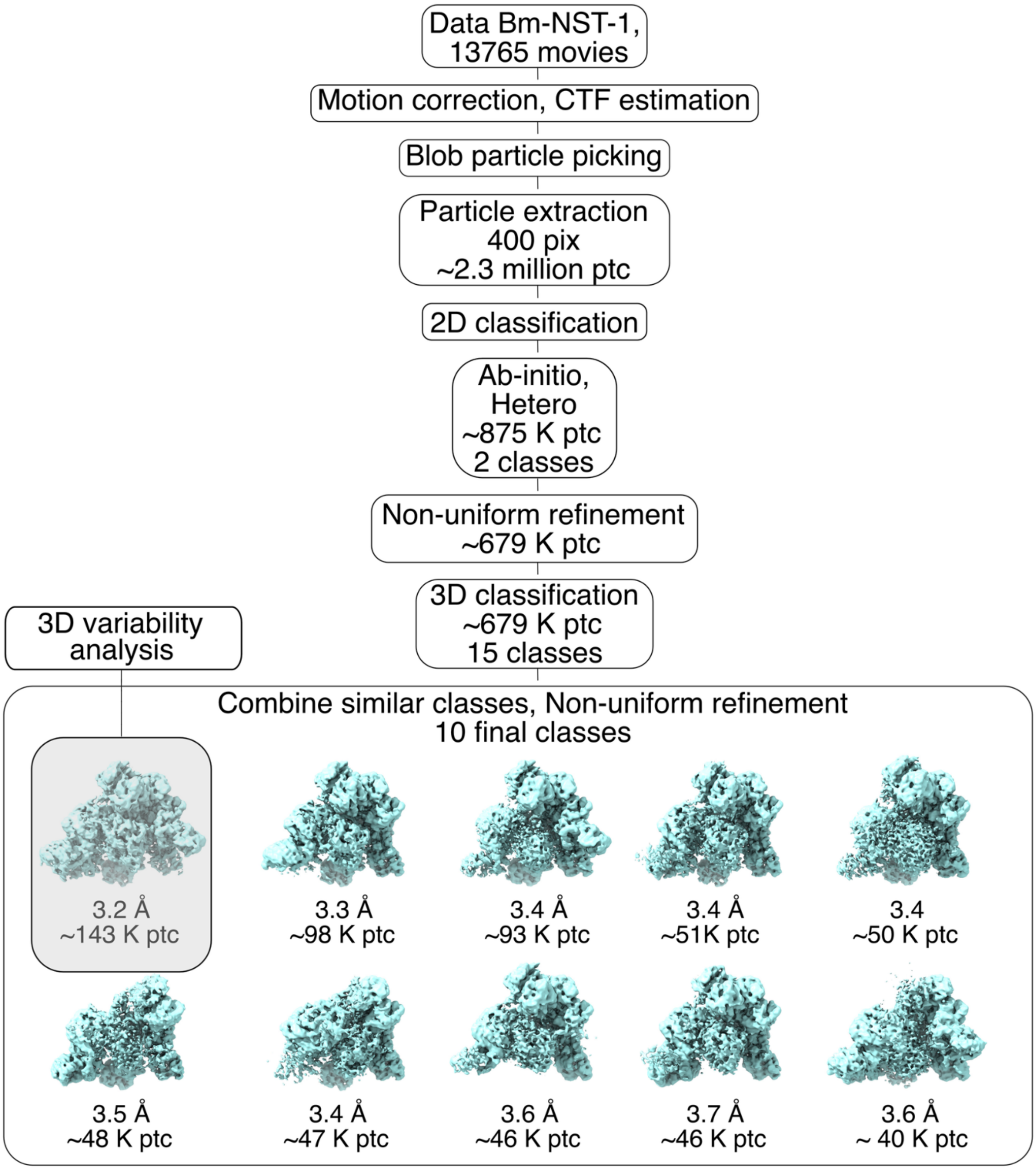


**Supplementary Fig. 7. Cryo-EM processing workflow for the CobST$^{Bm}$ sample.** For the Extract job, box size (full/Fourier-cropped) in pixels (pix) is indicated. For Extract, Hetero-, and NU-refinement jobs, the approximate number of used particles (ptc) and achieved resolution are indicated. The gray semi-transparent box indicates the final NU-refinement used for model building. CTF, contrast transfer function.

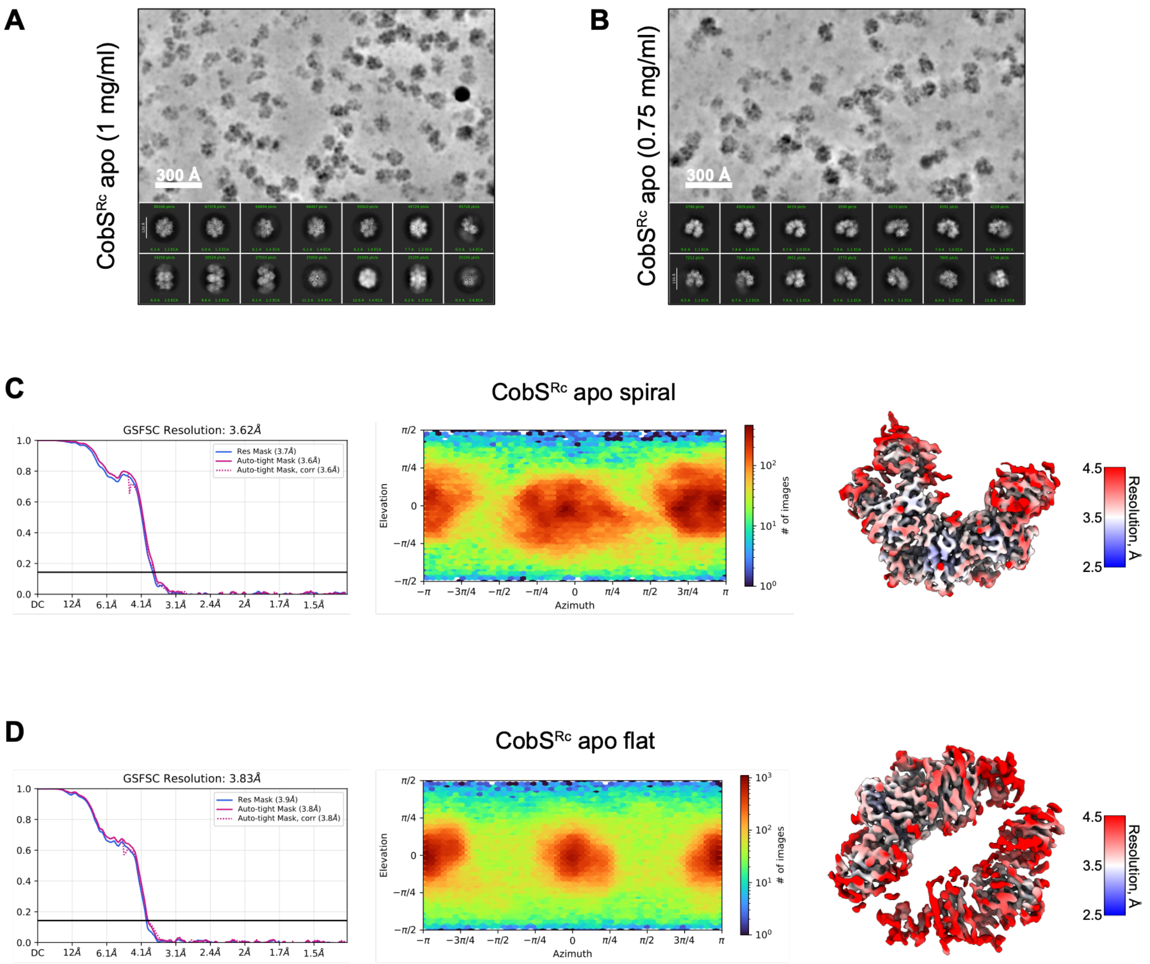


**Supplementary Fig. 8. Single-particle cryo-EM data analysis and validation for CobS$^{Rc}$.** **A, B.** Representative cryo-EM denoised micrograph (top) and 2D class averages (bottom) for the CobS$^{Rc}$ apo sample at 0.9-1 mg/ml (**A**) and at 0.75 mg/ml (**B**). **C, D.** Left, gold standard Fourier shell correlation (GSFSC) curves (generated in cryoSPARC), middle, angular distribution heatmap plots (generated in cryoSPARC), right, local resolution estimation map (generated in cryoSPARC, are colored by local resolution) for the CobS$^{Rc}$ apo spiral ring (conformation B) (**C**) and CobS$^{Rc}$ apo flat ring (conformation A) (**D**).

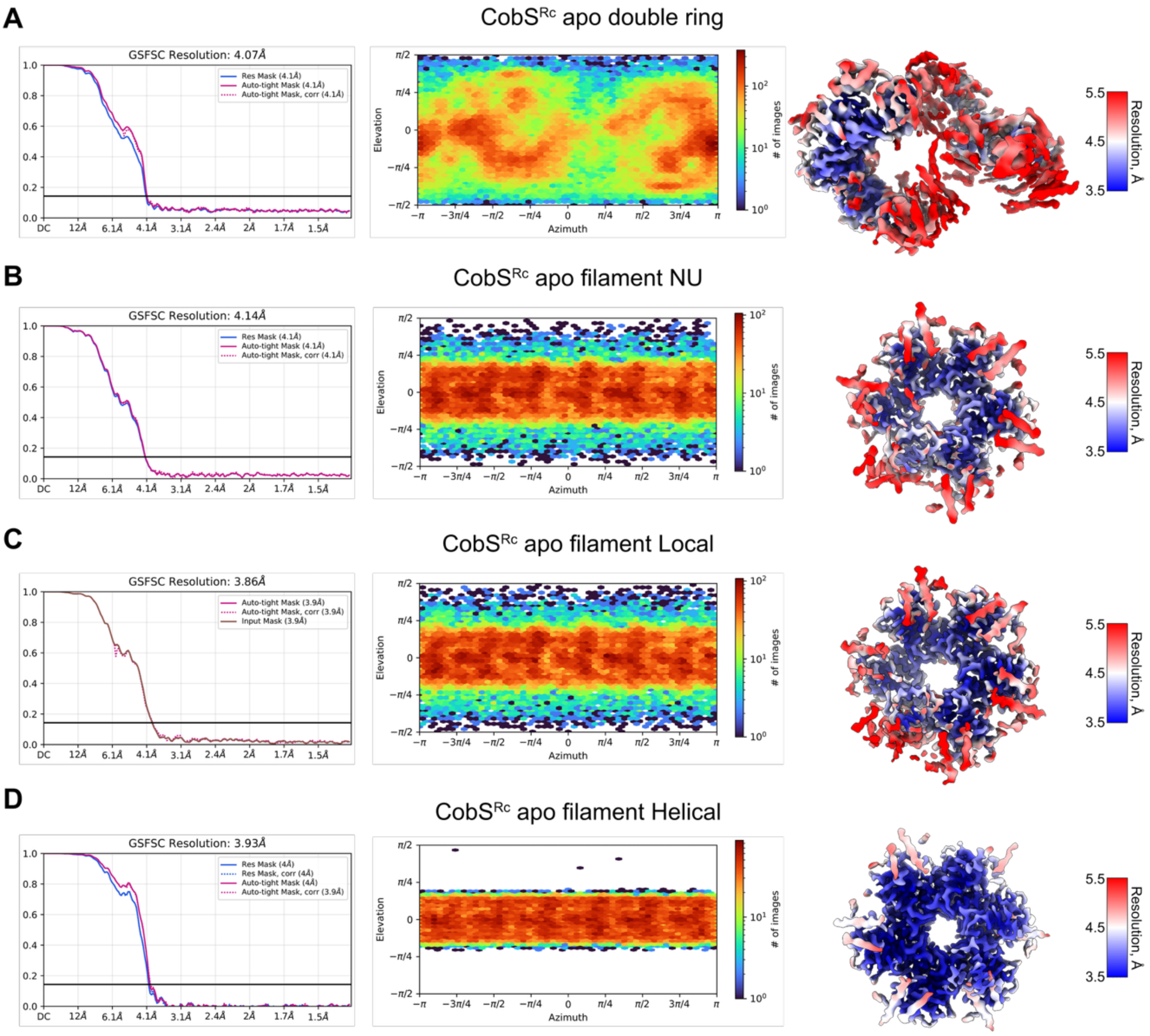


**Supplementary Fig. 9. Single-particle cryo-EM data analysis and validation for filament and infinite ring CobS$^{Rc}$. A-D.** Left, gold standard Fourier shell correlation (GSFSC) curves (generated in cryoSPARC), middle, angular distribution heatmap plots (generated in cryoSPARC), right, local resolution estimation map (generated in cryoSPARC, are colored by local resolution) for the CobS$^{Rc}$ apo conformation D double ring (**A**) and the CobS$^{Rc}$ apo filament (conformation C) (**B,** NU-refinement; **C,** Local refinement; **D**, Helical refinement).

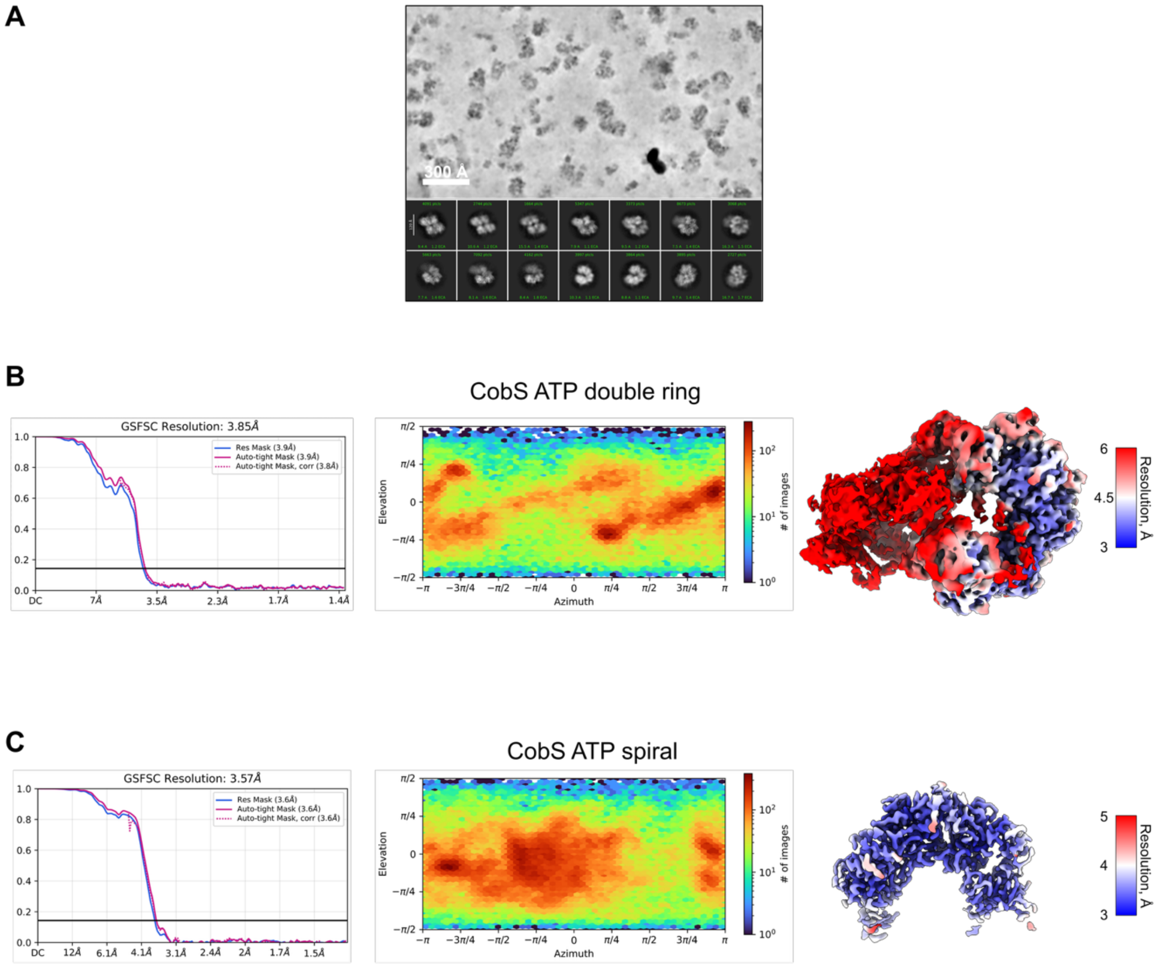


**Supplementary Fig. 10. Single-particle cryo-EM data analysis and validation of the CobS$^{Rc}$ sample incubated with ATP. A.** Representative cryo-EM denoised micrograph (top) and 2D class averages (bottom) for the CobS$^{Rc}$ sample incubated with ATP. **B,C.** Left, gold standard Fourier shell correlation (GSFSC) curves (generated in cryoSPARC), middle, angular distribution heatmap plots (generated in cryoSPARC), right, local resolution estimation map (generated in cryoSPARC, colored by local resolution) for the CobS$^{Rc}$ ATP-bound double ring conformation (**B**) and CobS$^{Rc}$ ATP-bound spiral ring conformation (**C**).

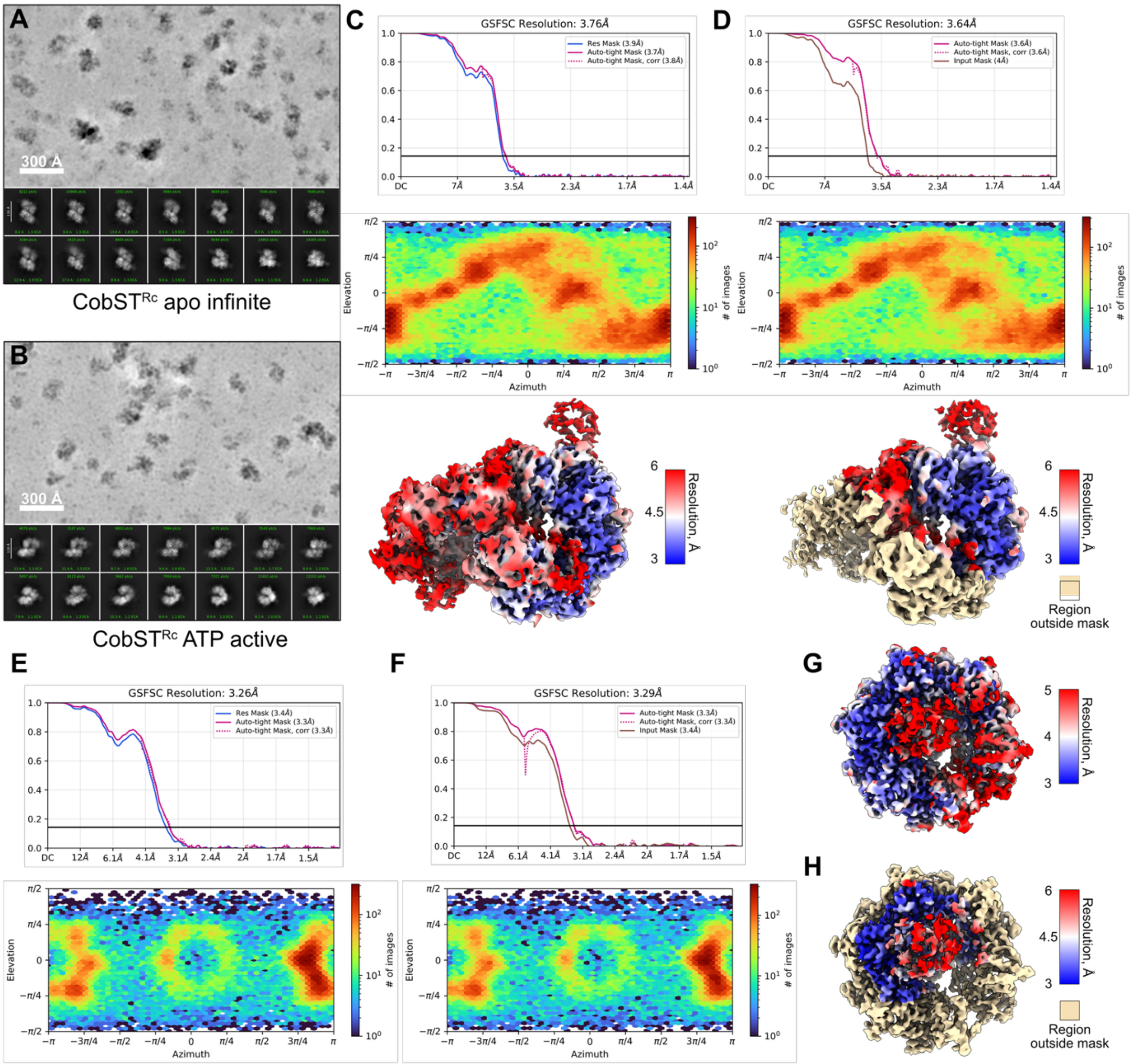


**Supplementary Fig. 11. Single-particle cryo-EM data analysis and validation of CobST$^{Rc}$.** **A,B.** Representative cryo-EM denoised micrograph (top) and 2D class averages (bottom) for the CobST$^{Rc}$ apo sample (**A**) and the sample incubated with ATP and $MgCl_2$ (**B**). **C,D.** Top, gold standard Fourier shell correlation (GSFSC) curves (generated in cryoSPARC), middle, angular distribution heatmap plots (generated in cryoSPARC), bottom, local resolution estimation map (generated in cryoSPARC, colored by local resolution) for the CobST$^{Rc}$ apo double ring conformation (**C,** NU-refinement; **D,** Local refinement). **E,F.** Top, gold standard Fourier shell correlation (GSFSC) curves (generated in cryoSPARC), bottom, angular distribution heatmap plots (generated in cryoSPARC) (**E,** NU-refinement; **F,** Local refinement) **G,H.** Local resolution estimation map (generated in cryoSPARC, colored by local resolution) for the CobST$^{Rc}$ ATP/ADP-bound active conformation (**G,** NU-refinement; **H,** Local refinement).

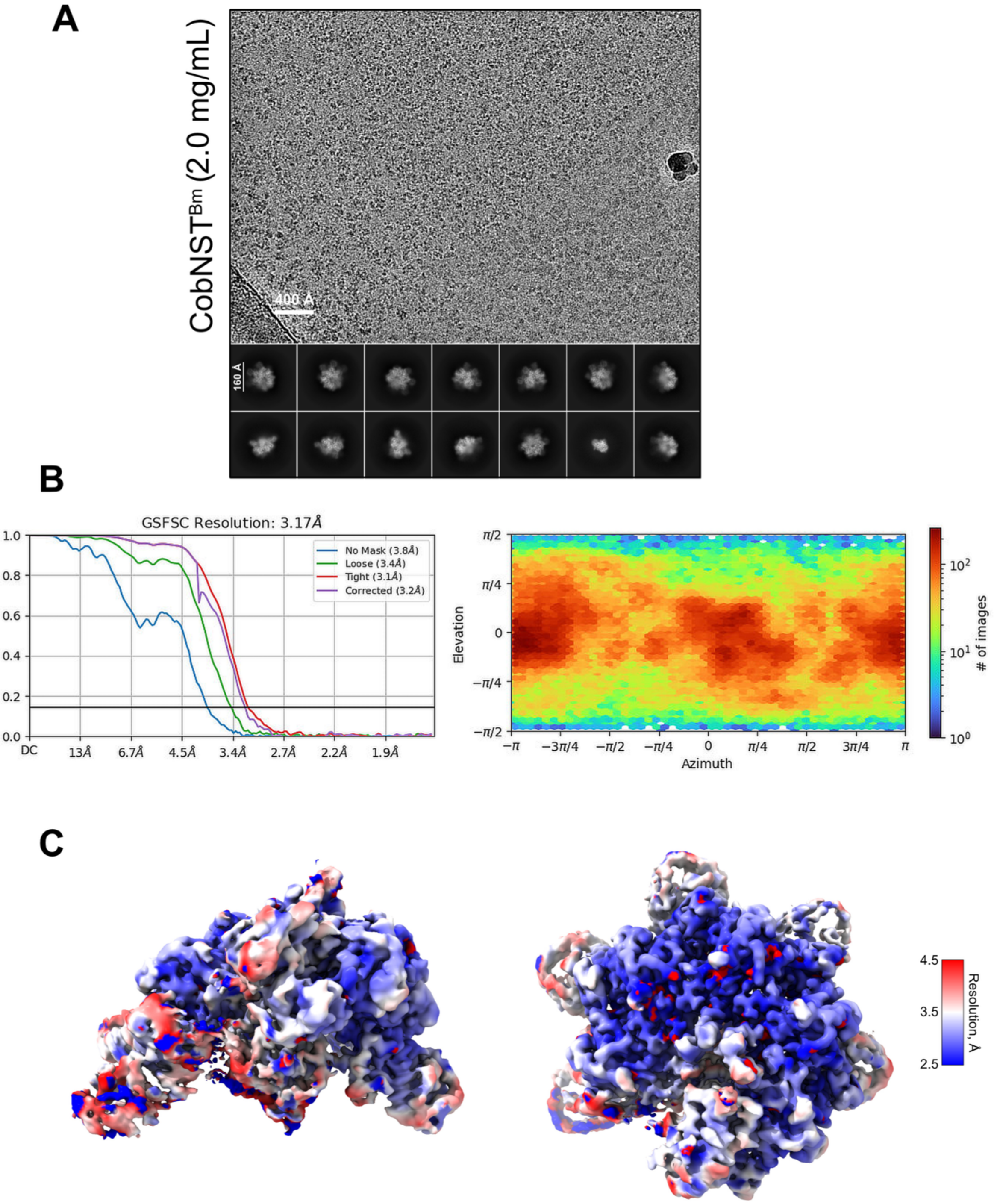


**Supplementary Fig. 12. Single-particle cryo-EM data analysis and validation of CobST$^{Bm}$. A.** Representative cryo-EM micrograph (top) and 2D class averages (bottom) for the CobST$^{Bm}$ sample. **B.** Left, gold standard Fourier shell correlation (GSFSC) curve (generated in cryoSPARC), right, angular distribution heatmap plot (generated in cryoSPARC). **C.** Local resolution estimation map (generated in cryoSPARC, colored by local resolution) for the CobST$^{Bm}$ complex (left, side view, right, top view).

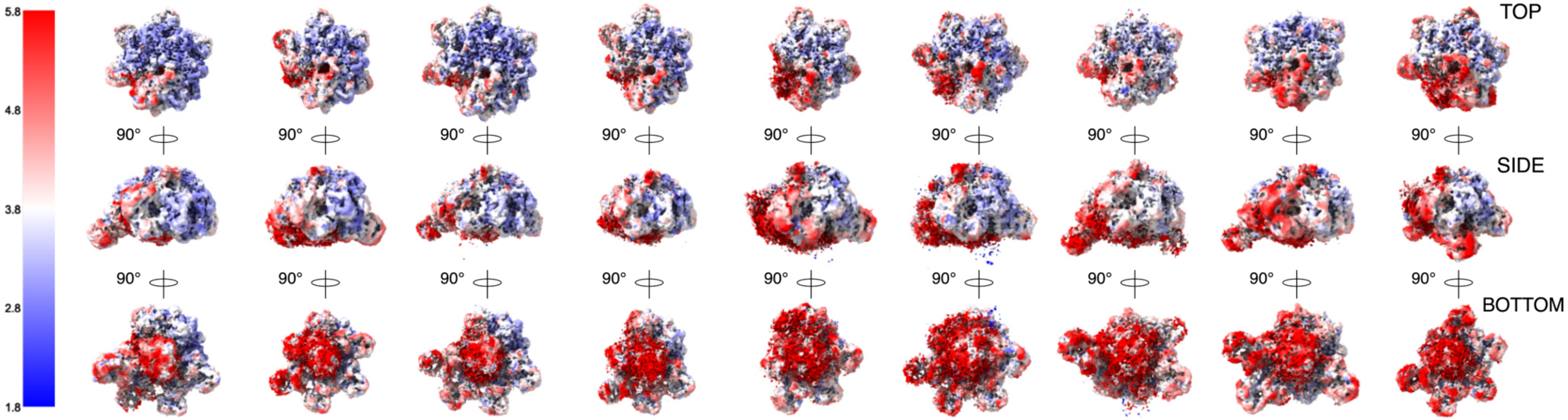


| | | | | | | | | | |
|---|---|---|---|---|---|---|---|---|---|
| particles (no.) | 98109 | 92836 | 51079 | 49975 | 48262 | 46777 | 46250 | 45965 | 39655 |
| resolution (Å) | 3.3 | 3.4 | 3.4 | 3.4 | 3.5 | 3.4 | 3.6 | 3.7 | 3.6 |
| composition ($S:T_N:T_C$) | 6:4:1 | 6:4:1 | 6:5:1 | 6:3:1 | 5:3:1 | 5:4:1 | 6:4:1 | 6:3:1 | 5:5:1 |

**Supplementary Fig. 13. Conformational variability of the CobST$^{Bm}$ complex in the left-handed spiral post-hydrolysis state.** Cryo-EM maps of CobST$^{Bm}$ complexes with different molecules of attached CobT$^{Bm}$ N-terminal domains, obtained by 3D classification of the consensus map but not used for model building (see Supplementary Fig. 7 for details of data processing), colored by local resolution. The numbers of observed CobS$^{Bm}$ subunits (S), CobT$^{Bm}$ N-terminal domains ($T_N$), and CobT$^{Bm}$ C-terminal VWA ($T_C$) domains are indicated below each map.

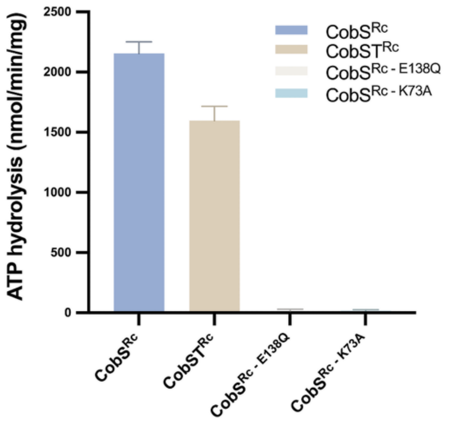


**Supplementary Fig. 14. ATPase activity of CobS$^{Rc}$ and CobST$^{Rc}$ proteins.** ATP hydrolysis was monitored as the decrease in NADH absorbance at 340 nm for 2 hours at 37 °C. The reactions contained 0.25 µg of CobS$^{Rc}$ protein in the presence of 10 mM ATP and 12 mM $MgSO_4$ (see Methods section for more details). The Walker A and Walker B motif mutants, CobS$^{Rc – K73A}$ and CobS$^{Rc – E138Q}$, respectively, were used as negative controls. ATP hydrolysis rates (nmol ATP per minute per milligram of protein) were calculated from the linear portion of the 340nm absorbance decrease. Data are shown as mean ± SD (CobS$^{Rc}$: 2155.9 ± 95.5 nmol/min/mg; CobST$^{Rc}$: 1596.6 ± 118.1 nmol/min/mg; CobS$^{Rc – E138Q}$: 12.4 ± 17.9 nmol/min/mg; CobS$^{Rc – K73A}$: 16.5 ± 8.9 nmol/min/mg).

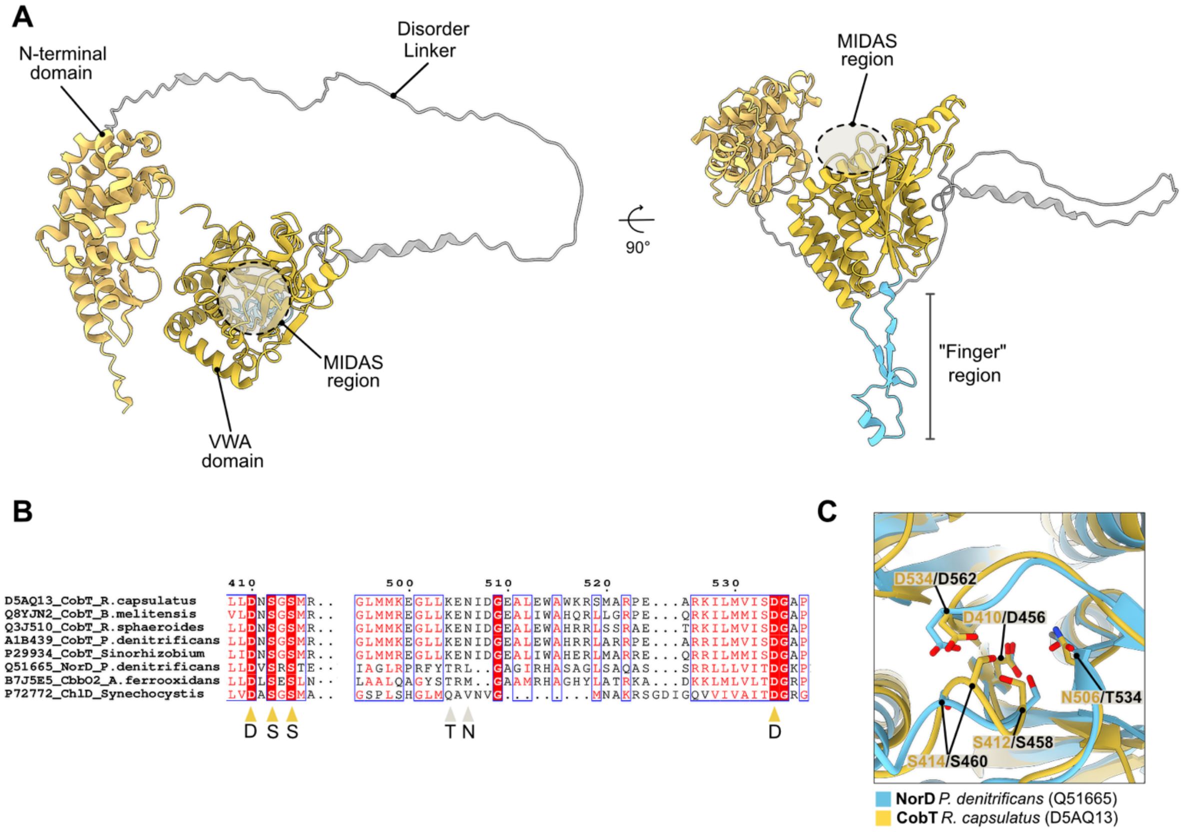


**Supplementary Fig. 15. Overview of the $\text{CobT}^{Rc}$ structure prediction. A.** Top (left) and side (right) views of the AlphaFold [1] prediction of $\text{CobT}^{Rc}$ (based on Uniprot entry: D5AQ13) represented as ribbons/slabs. N-terminal domain (light yellow), predicted disorder linker (grey) and VWA domain (darker yellow). The MIDAS putative region is highlighted as a circle, and the Finger-like region is coloured blue. **B.** Protein multiple sequence alignment of the putative MIDAS region (DXSXS…T…D) for different CobT proteins and other MoxR $\text{AAA}^{+}$ VWA-partner proteins (NorD, CbbO and ChlD). Conserved MIDAS residues are highlighted with yellow triangles and the variable residues are highlighted in grey triangles. **C.** Close-up view of the AlphaFold prediction of the putative $\text{CobT}^{Rc}$ and the previously characterized *P. denitrificans* NorD MIDAS regions [2,3]. Putative $\text{CobT}^{Rc}$ MIDAS residues highlighted in yellow.

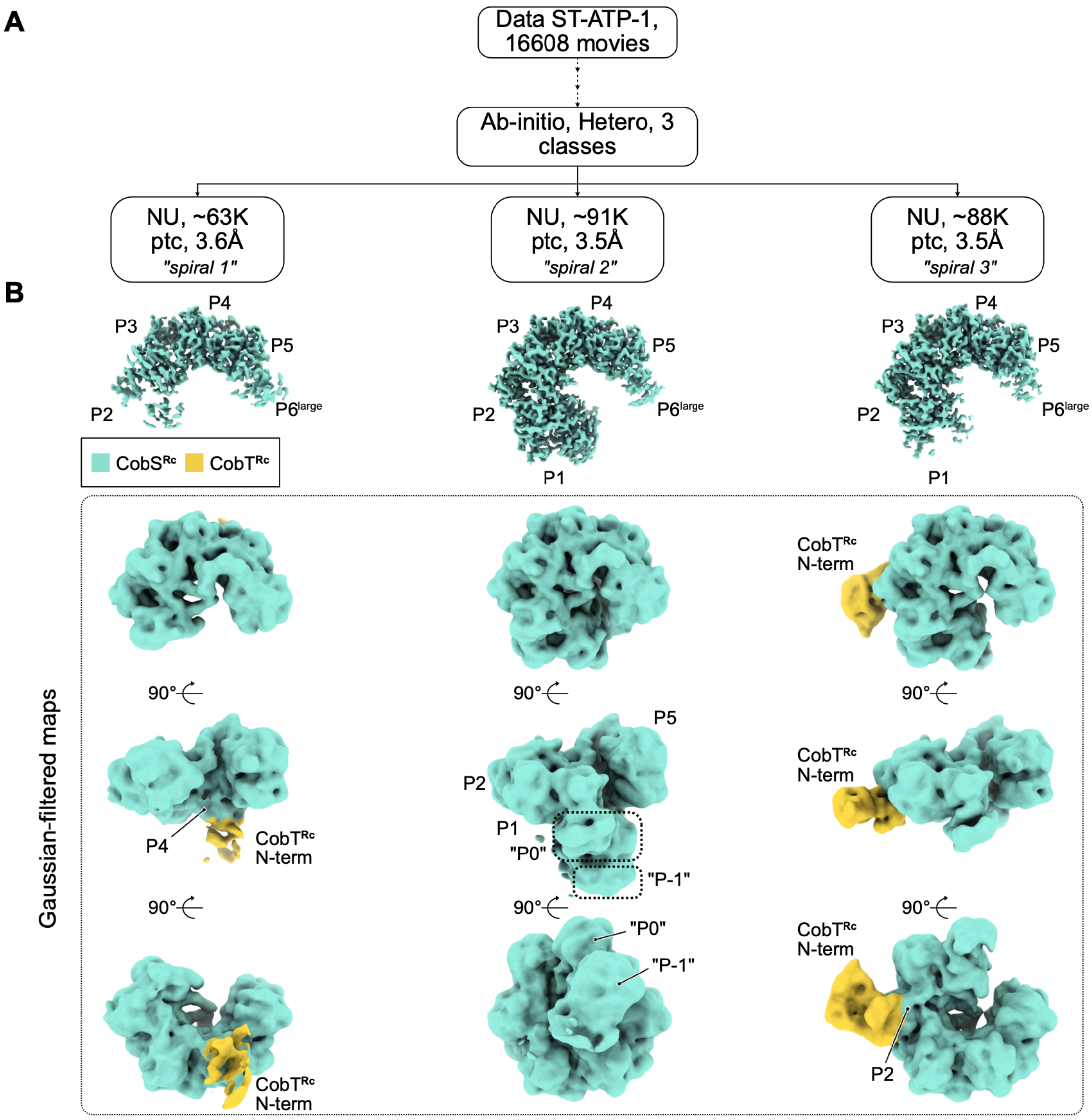


**Supplementary Fig. 16. Conformational variability of the CobST$^{Rc}$ complex in the left-handed spiral state from the sample with ATP and $MgCl_2$. A.** Cryo-EM processing workflow resulting in three maps (**B**) representing different spiral conformations of CobS$^{Rc}$ (teal), free or complexed with CobT$^{Rc}$ (gold); see **Supplementary Fig. 6** for processing details. **B.** Top row, maps representing the three spiral conformations from (**A**) shown at contour level of 0.1. CobS$^{Rc}$ protomers are labelled (P-1 to P6, corresponding to the labelling in **Figure 1**). Three lower rows (in box), Gaussian-filtered maps generated from the maps in the top row, viewed from different angles. CobT$^{Rc}$ N-terminal domains (gold) with their respective interacting CobS$^{Rc}$ protomers are labelled.

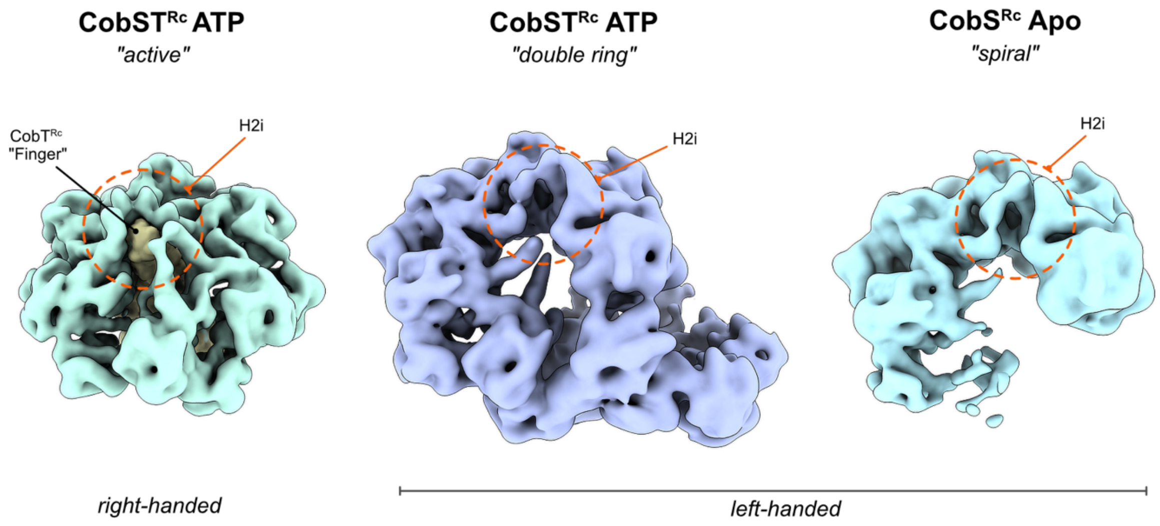


**Supplementary Fig. 17. Comparison of the handedness of CobS$^{Rc}$ helical conformations across different states.** Left, the active CobST$^{Rc}$ complex: CobS$^{Rc}$ H2i insertions (dashed circle, orange) displaying a right-handed arrangement (P2 to P4) engaging CobT$^{Rc}$ through the central pore. CobT$^{Rc}$ is represented as a volume generated using the molmap function in ChimeraX (gold, res. 4 Å) and CobS$^{Rc}$ as a Gaussian-filtered map (teal). Middle, Gaussian-filtered map of the double ring CobST$^{Rc}$ conformation displayed at a lower contour level (CobT$^{Rc}$ N-terminal not visible), highlighting H2i insertions (dashed circle, orange) showing a left-handed arrangement of CobS$^{Rc}$ subunits. Right, Gaussian-filtered map of the apo CobS$^{Rc}$ left-handed spiral ring. H2i inserts highlighted in a dashed circle, orange.

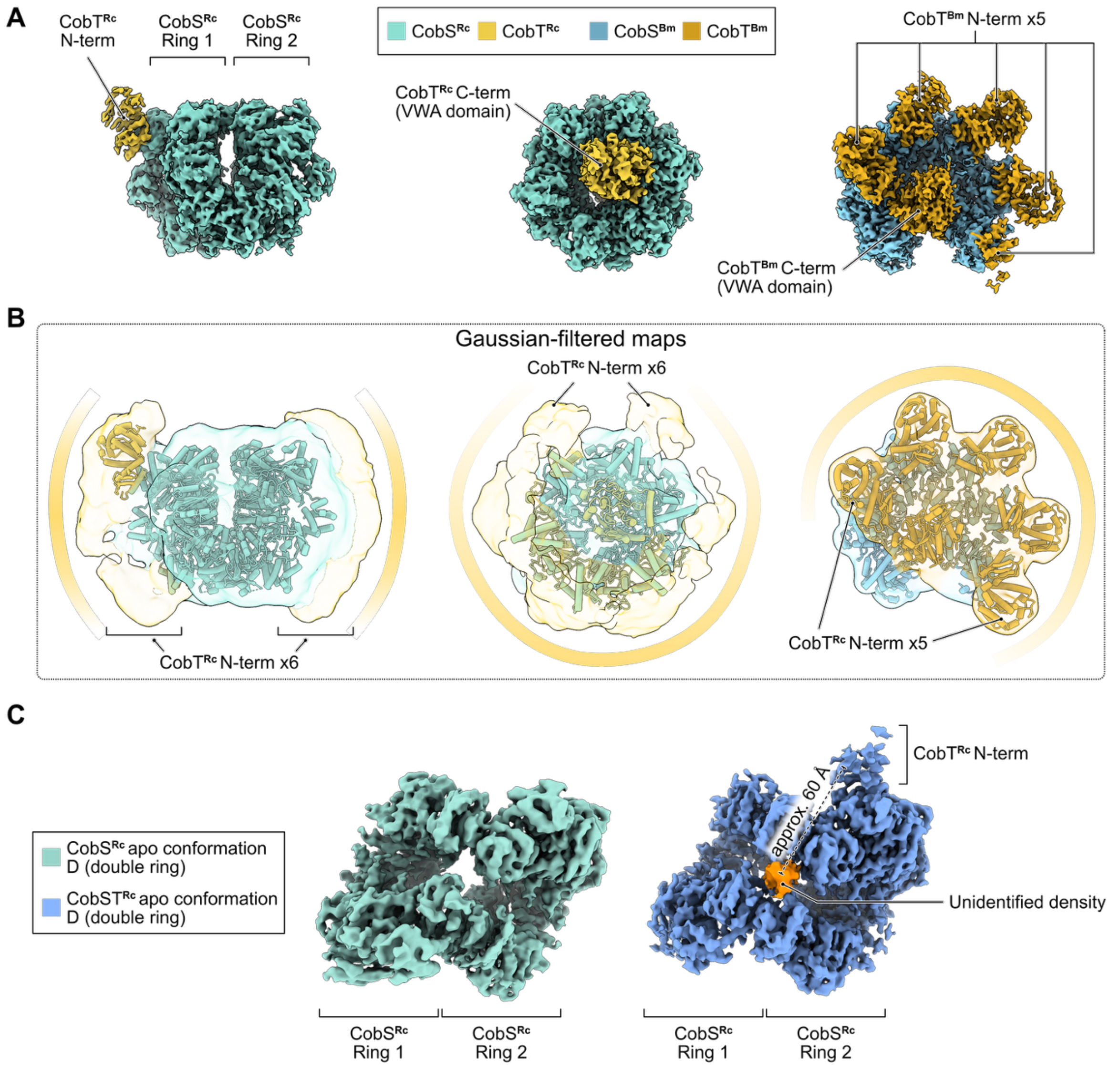


**Supplementary Fig. 18. Decoration of the CobS oligomers by CobT N-terminal domains. A.** Cryo-EM maps of the apo double ring CobST$^{\mathrm{Rc}}$ conformation (left), active right-handed helical closed CobST$^{\mathrm{Rc}}$ conformation (middle), and post-hydrolysis left-handed helical conformation of CobST$^{\mathrm{Bm}}$ (right). CobS$^{\mathrm{Rc}}$ and CobT$^{\mathrm{Rc}}$ are colored teal and gold, respectively; CobS$^{\mathrm{Bm}}$ and CobT$^{\mathrm{Bm}}$ are colored light blue and orange, respectively. **B.** Gaussian-filtered maps from **A** with fitted molecular models of the respective complexes shown as cylinders/stubs. Additional densities decorating the periphery of CobS$^{\mathrm{Rc}}$ rings, assigned to CobT$^{\mathrm{Rc}}$ N-terminal domains, are shown. **C.** Comparison of the double ring (conformation D) of CobS$^{\mathrm{Rc}}$ and CobST$^{\mathrm{Rc}}$ apo maps, highlighting the unidentified density within the CobST$^{\mathrm{Rc}}$ ring interface in orange. Distance calculated between the centre of this density and the density for the CobT$^{\mathrm{Rc}}$ N-terminal domain (~ 60 Å).

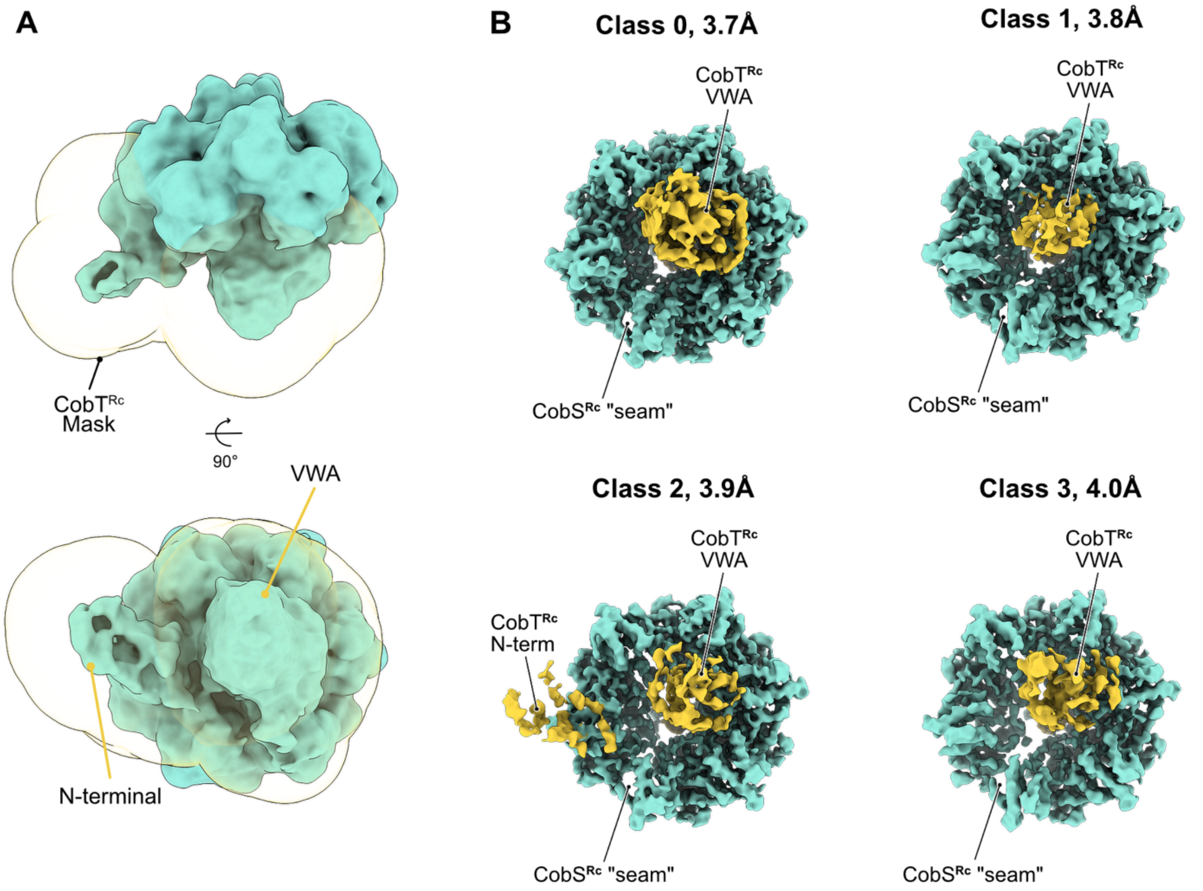


**Supplementary Fig. 19. Focused 3D classification analysis of CobST$^{Rc}$ highlights flexibility of the CobT$^{Rc}$ domains. A.** Gaussian-filtered consensus map of the active CobST$^{Rc}$ (teal, threshold: 0.00521), focused mask (threshold: 0.00521, dilation radius ~4.1 Å, soft padding width ~20.4 Å) used for 3D variability analysis displayed as semi-transparent (gold). The mask encompasses the CobT$^{Rc}$ N-terminal and C-terminal (VWA) domains (gold). **B.** Different class maps (class 0 – class 3) obtained in the 3D classification. Class 0 (~15K particles, 34.4%) displays a well-resolved VWA as seen in the consensus cryo-EM map (Fig. 4). Class 1 (11K particles, 24.5%) and class 3 (8K particles, 17.5%) show a partial occupancy of the CobT$^{Rc}$ VWA domain. Class 2 (10K particles, 23.5%) displays density for both CobT$^{Rc}$ VWA and N-terminal domains.

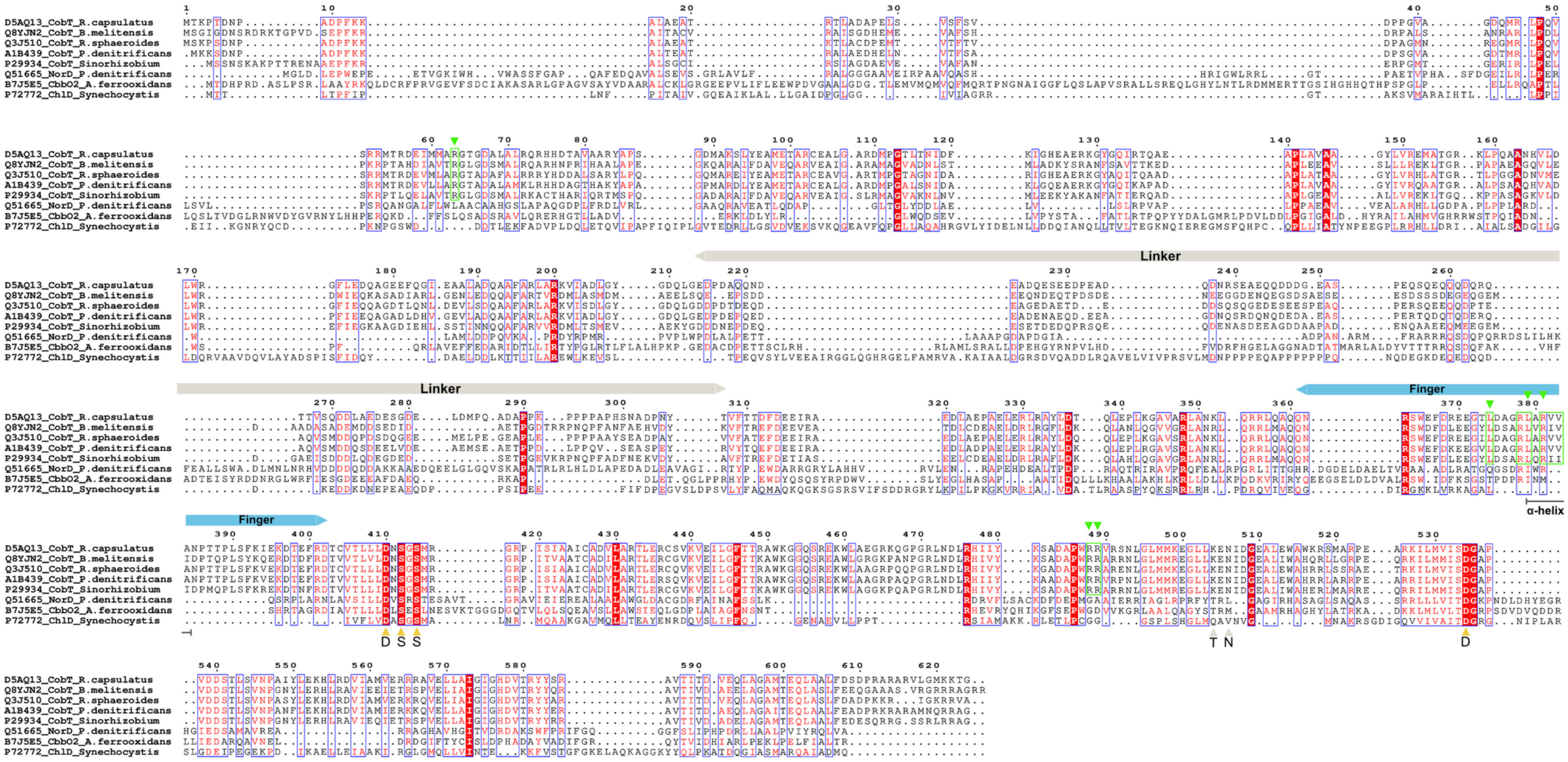

**Supplementary Fig. 20. Multiple protein sequence alignment of CobT from different organisms and other MoxR AAA$^{+}$ partner VWA proteins.** Each sequence is labelled and identified by its Uniprot entry. The CobT$^{Rc}$ linker region (beige) and finger region (blue) within the VWA domain are highlighted as boxes. The MIDAS residues are highlighted as triangles (yellow and grey, as in Supplementary Fig. 18). CobT conserved residues observed to be important for interacting with the AAA$^{+}$ CobS are highlighted as green boxes and triangles. Multiple sequence alignment was performed using MAFFT in SnapGene software (www.snapgene.com) and represented based on conservation in ESPript [4].

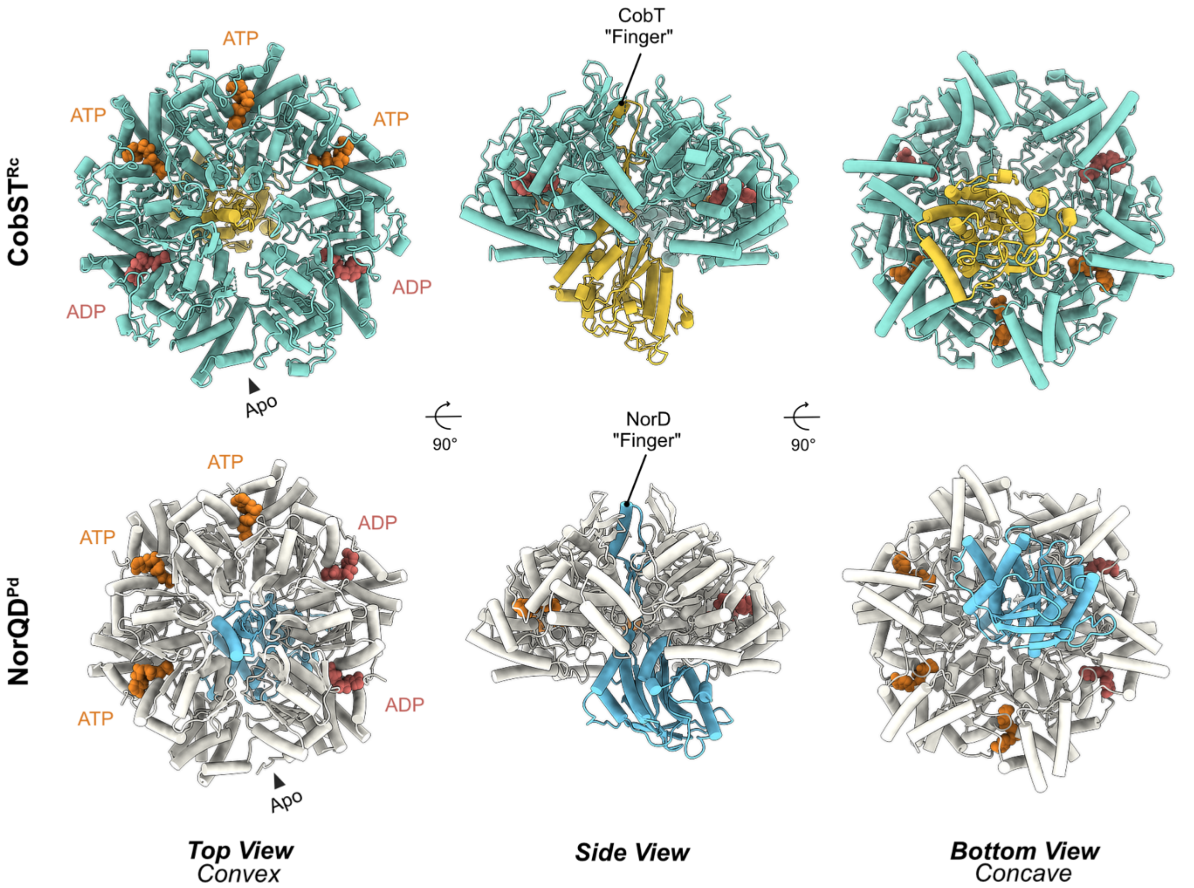


**Supplementary Fig. 21. Comparison of the CobST$^{Rc}$ (active) and NorQD$^{Pd}$ (ATPase-deficient) structures.** In the top row, the active *R. capsulatus* CobST$^{Rc}$ structure and in the lower row, the *P. denitrificans* Walker B NorQ mutant in complex with the VWA domain of NorD (PDB: 9QH6) [3]. Models are represented as cylinders/stubs. Left: top views (ring convex side). Middle: side views. Right: bottom views (ring concave side).

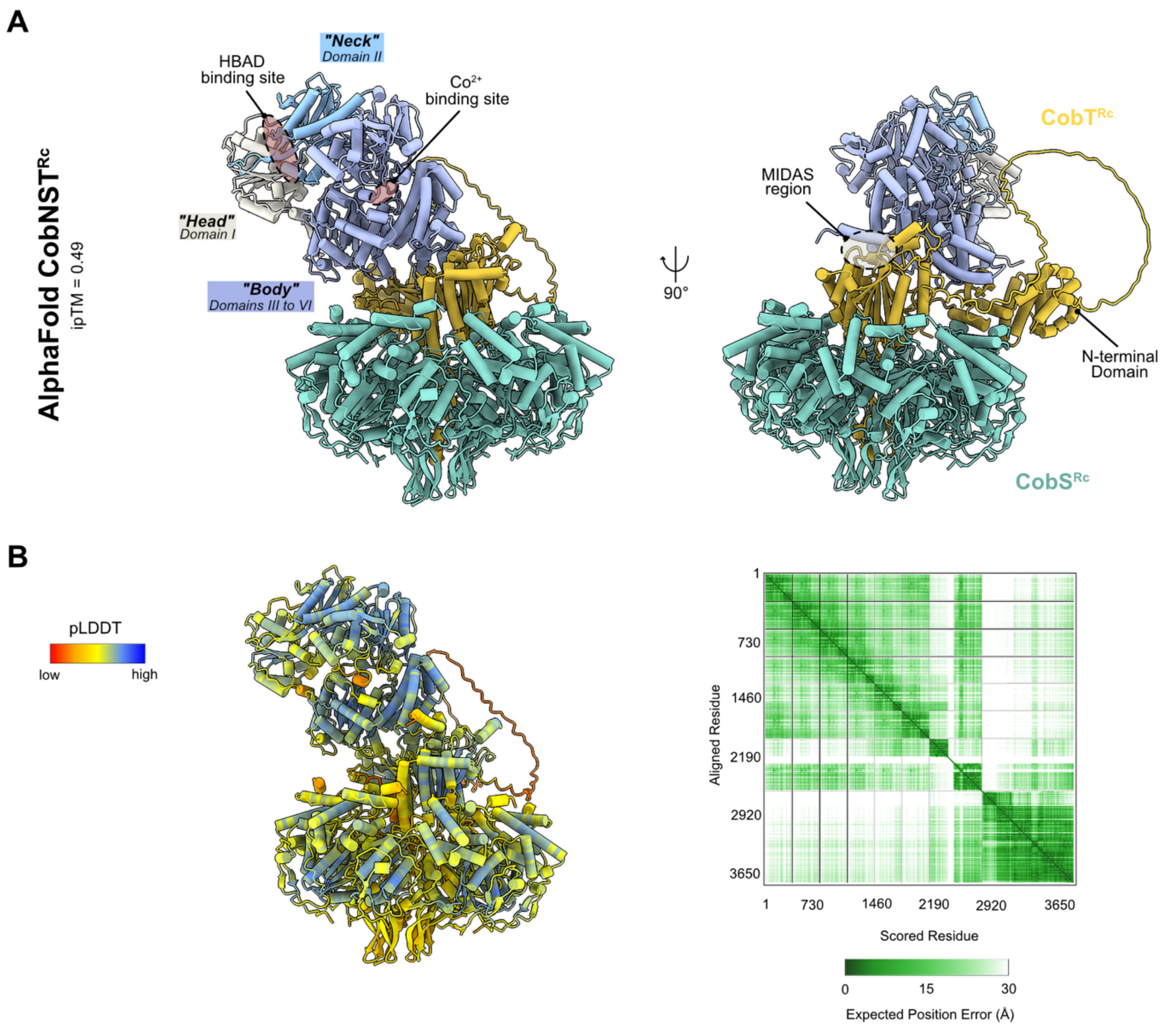


**Supplementary Fig. 22. AlphaFold-Multimer model of the overall *R. capsulatus* CobNST complex. A.** Side views of AlphaFold 3 [1] prediction of the CobNST$^{Rc}$ complex (6:1:1, CobS:CobT:CobN) in the absence of nucleotides and ions (interface-predicted template modeling, ipTM = 0.49). Model represented as cylinders/stubs. CobS subunits are colored in teal and CobT is colored in gold. CobN domain I ("head") is colored in beige, domain II ("neck") is colored in blue and domains III to VI ("body") are colored in purple. HBAD substrate binding site ("head"/"neck" interface) and cobalt binding site are highlighted as red circles. MIDAS motif region is highlighted as a grey circle. **B.** Confidence assessments of the prediction. Left, predicted CobNST$^{Rc}$ structure (view as in **A.** left) colored based on the per-residue predicted local distance difference test (pLDDT) score; red indicates low-confidence regions and blue indicates high-confidence regions. Right, predicted alignment error (PAE) plot showing the expected error in the relative positions of residues, with lower values indicating higher confidence. The representations and plots were generated with ChimeraX [5] software.

### Description of Additional Supplementary Files

**Supplementary Movie 1. Rotation of the small AAA$^+$ domain of the subunit P1 from the CobS$^{Rc}$ flat ring conformation.** The movie shows the superposition of subunits P1, P2 and P6 from the CobS$^{Rc}$ flat ring conformation (see also Fig. 2D) and the rotation of the small AAA$^+$ domain of P1 relative to the other represented subunits, illustrated by morphing between subunits P1 and P6. Models are represented as licorice/ovals. Colored as in Fig.1.

**Supplementary Movie 2. CobS$^{Rc}$ flat ring opening to spiral ring.** The movie shows the morph between CobS$^{Rc}$ flat and spiral ring conformations, illustrating the possible transition between them. Models are represented as licorice/ovals. Colored as in Fig.1.

**Supplementary Movie 3. Rotation of the small AAA$^+$ domain of the subunit P6 from the CobS$^{Rc}$ double ring conformation.** The movie shows the superposition of subunits P1, P2 and P6 from the CobS$^{Rc}$ double ring conformation (see Fig. 2E and 2F) and the rotation of the small AAA$^+$ domain of P6 relative to the other represented subunits, illustrated by morphing between subunits P6 and P1. Models are represented as licorice/ovals. Colored as in Fig.1.

**Supplementary Movie 4. Structural transitions between CobST active and CobST post-hydrolysis conformations.** The movie represents the morph between the active CobST$^{Rc}$ structure (see Fig. 4) and a hypothetical post-hydrolysis model of CobST$^{Rc}$ generated by aligning of CobST$^{Rc}$ subunits to the post-hydrolysis CobST$^{Bm}$ complex (see Fig. 6) using the matchmaker function in ChimeraX. The morph highlights the possible movement of the CobT VWA domain. Molecular models are represented as licorice/ovals and colored as in Fig. 1 and Fig. 4. CobT finger region and CobS P6 small domain are omitted to avoid clashes.

**Supplementary Movie 5. 3D Variability analysis of CobST$^{Bm}$.** The movie displays the structural dynamics of the post-hydrolysis CobST$^{Bm}$ complex.

**Supplementary Table 1.** List of primers and DNA fragments used in this work.

| Gene | Direction | Sequence (5' – 3') | Source |
|---|---|---|---|
| ***cobS***$^{Rc}$ | Forward | gtacatatgctcgaccagaccgccaagc | This study |
| | Reverse | gtactcgaggcgccccgcgatgacg | This study |
| ***cobS***$^{Bm}$ | Forward | cagcatatgatgaacaaggttgagc | [6] |
| | Reverse | cagactagtctcatcaggcgagaacg | [6] |
| ***cobT***$^{Bm}$ | Forward | catcatatgtcgggccaaatgtcag | [6] |
| | Reverse | catactagtgctctatcttcgaccg | [6] |
| ***cobN***$^{Bm}$ | Forward | cgaattaatatgcatctcctgcttgcccag | [6] |
| | Reverse | cgaactagttctcagccatttatcgc | [6] |
| ***cobS***$^{Rc}$ **- E138Q** | Forward | gttcgatcagtacgacgccggcc | This study |
| | Reverse | acaatcgcggtcgggttgcgca | This study |
| ***cobS***$^{Rc}$ **- K73A** | Forward | gcaccggcgcgtcgacccatatc | This study |
| | Reverse | cgtgatagccctgaatcatcacgcg | This study |
| ***cobS***$^{Rc}$ (co-expression) | Forward | gttaagtataagaaggagatatacatatgctcgaccagaccgccaag | This study |
| | Reverse 1 | atgatgtggaccttgaaacaaaacttctaagcgccccgcgatgacggc | This study |
| | Reverse 2 | cgcagcagcggtttctttaccagacttagtgatgatgatgatgatgtggaccttgaaacaaaacttctaagc | This study |
| ***cobT***$^{Rc}$ (co-expression) | Forward | gttagaagttttgtttcaaggtccaacgaagcccactgacaacc | This study |
| | Reverse | ctttctgttcgacttaagcattatgcttaccccgtctttttcatccc | This study |
| | dsDNA | gtttaactttaagaaggagatatacatgagtgcttggagtcatccacaattcgaaaaaggtggaggttccggaggtggatcgggaggttctgcctggtctcatccacaatttgagaagttagaagttttgtttcaaggtcca | This study |

### Supplementary Table 2. Cryo-EM data collection, refinement and validation statistics for CobS$^{Rc}$ and CobST$^{Rc}$ datasets.

| | Apo CobS conf. A (flat ring) PDB XXXX EMDB-XXXX | Apo CobS conf. B (spiral ring) PDB XXXX EMDB-XXXX | Apo CobS conf. C (filament), NU-refinement) PDB XXXX EMDB-XXXX | Apo CobS conf. C (filament), Local refinement) EMDB-XXXX | Apo CobS conf. C (filament), Helical refinement EMDB-XXXX | Apo CobS conf. D (double ring) PDB XXXX EMDB-XXXX | Nucleotide-bound CobS conf. B (spiral ring) EMDB-XXXX | Nucleotide-bound CobS conf. D (double ring) EMDB-XXXX | Apo CobST conf. D (double ring) NU refinement PDB XXXX EMDB-XXXX | Apo CobST conf. D (double ring) Local refinement EMDB-XXXX | Nucleotide-bound active CobST NU refinement PDB XXXX EMDB-XXXX | Nucleotide-bound active CobST Local refinement EMDB-XXXX |
|---|---|---|---|---|---|---|---|---|---|---|---|---|
| **Data collection and processing** | | | | | | | | | | | | |
| Magnification | 165000× | 165000× | 165000× | 165000× | 165000× | 165000× | 165000× | 165000× | 165000× | 165000× | 165000× | 165000× |
| Voltage (kV) | 200 | 200 | 200 | 200 | 200 | 200 | 200 | 200 | 200 | 200 | 200 | 200 |
| Electron exposure (e$^{-}$/Å$^2$) | 50 | 50 | 50 | 50 | 50 | 50 | 50 | 50 | 50 | 50 | 50 | 50 |
| Defocus range (μm) | 0.8 - 1.8 | 0.8 - 1.8 | 0.8 - 1.8 | 0.8 - 1.8 | 0.8 - 1.8 | 0.8 - 1.8 | 0.8 - 1.8 | 0.8 - 1.8 | 0.8 - 1.8 | 0.8 - 1.8 | 0.8 - 1.8 | 0.8 - 1.8 |
| Pixel size (Å) | 0.68 | 0.68 | 0.68 | 0.68 | 0.68 | 0.68 | 0.68 | 0.68 | 0.68 | 0.68 | 0.68 | 0.68 |
| Extraction box size (pixels) | 432/450 | 432/450 | 432/450 | 432/450 | 432/450 | 432/450/512 | 432/450 | 432/450/512 | 450/512 | 450/512 | 450 | 450 |
| Symmetry imposed | C1 | C1 | C1 | C1 | Helical symmetry Rise: 10.11Å; Twist: -61.78Å, symmetry order: 5 | C1 | C1 | C1 | C1 | C1 | C1 | C1 |
| Initial particle images (no.) | 242007 | 242007 | 3084614 | 3084614 | 3084614 | 2323800 | 662599 | 662599 | 832457 | 832457 | 1114860 | 1114860 |
| Final particle images (no.) | 231064 | 191066 | 60101 | 60101 | 60017 | 119435 | 147276 | 84444 | 103511 | 103511 | 46304 | 46304 |
| Map resolution (Å) | 3.8 | 3.6 | 4.1 | 3.9 | 3.9 | 4.1 | 3.6 | 3.9 | 3.8 | 3.6 | 3.3 | 3.3 |
| FSC threshold | 0.143 | 0.143 | 0.143 | 0.143 | 0.143 | 0.143 | 0.143 | 0.143 | 0.143 | 0.143 | 0.143 | 0.143 |
| Map resolution range (Å) | 1.4-57.7 | 3.1-56.5 | 3.6-64.2 | 3.4-35.4 | 1.4-54.2 | 1.4-64.2 | 3.1-56.4 | 3.4-63.3 | 1.4-60.9 | 1.4-65.2 | 2.9-54.3 | 1.4-52.5 |
| **Model Building and refinement** | | | | | | | | | | | | |
| Initial model used (Uniprot code) | D5AQ14 (CobS$^{Rc}$) | D5AQ14 (CobS$^{Rc}$) | D5AQ14 (CobS$^{Rc}$) | - | - | D5AQ14 (CobS$^{Rc}$) | D5AQ14 (CobS$^{Rc}$) | D5AQ14 (CobS$^{Rc}$) | D5AQ14 (CobS$^{Rc}$) and D5AQ13 (CobT$^{Rc}$) | - | D5AQ14 (CobS$^{Rc}$) and D5AQ13 (CobT$^{Rc}$) | - |
| Map sharpening *B* factor (Å$^2$) | 130.5 | 117.1 | 94.6 | 92.7 | 111.3 | 113.8 | 112.5 | 83.9 | 102.4 | 88.0 | 73.3 | 70.5 |
| Model composition | | | | - | - | | - | - | | - | | - |
| Non-hydrogen atoms (no.) | 22429 | 19682 | 4257 | | | 48819 | | | 50930 | | 33244 | |
| Protein residues (no.) | 1390 | 1224 | 266 | | | 3055 | | | 3296 | | 2086 | |
| Ligands (no.) | 0 | 0 | 0 | | | 0 | | | 0 | | MG: 2<br>ATP: 3<br>ADP: 2 | |
| Mean *B* factors (Å$^2$) | | | | - | - | | - | - | | - | | - |
| Protein | 114.16 | 73.90 | 114.54 | | | 194.97 | | | 95.06 | | 81.95 | |
| Ligand | - | - | - | | | - | | | - | | 55.08 | |
| R.m.s. deviations | | | | - | - | | - | - | | - | | - |
| Bond lengths (Å) | 0.003 | 0.003 | 0.002 | | | 0.004 | | | 0.01 | | 0.002 | |
| Bond angles (°) | 0.549 | 0.551 | 0.453 | | | 0.903 | | | 0.722 | | 0.555 | |
| Validation | | | | - | - | | - | - | | - | | - |
| MolProbity score | 2.66 | 2.49 | 2.31 | | | 2.14 | | | 2.43 | | 2.59 | |
| Clashscore | 12.33 | 10.49 | 9.41 | | | 12.07 | | | 18.06 | | 8.42 | |
| Poor rotamers (%) | 5.36 | 4 | 2.63 | | | 0.00 | | | 1.65 | | 5.59 | |
| Ramachandran plot | | | | - | - | | - | - | | - | | - |
| Favored (%) | 91.65 | 91.70 | 91.80 | | | 90.32 | | | 91.17 | | 89.31 | |
| Allowed (%) | 8.11 | 7.87 | 7.81 | | | 9.40 | | | 8.54 | | 10.40 | |
| Disallowed (%) | 0.24 | 0.44 | 0.39 | | | 0.28 | | | 0.29 | | 0.29 | |

**Supplementary Table 3. Cryo-EM data collection, refinement and validation statistics for the CobST$^{Bm}$ dataset.**

| | Class 1 EMDB PDB | Class 2 EMDB | Class 3 EMDB | Class 4 EMDB | Class 5 EMDB | Class 6 EMDB | Class 7 EMDB | Class 8 EMDB | Class 9 EMDB | Class 10 EMDB |
|---|---|---|---|---|---|---|---|---|---|---|
| **Data collection and processing** | | | | | | | | | | |
| Magnification | 105000 × | 105000 × | 105000 × | 105000 × | 105000 × | 105000 × | 105000 × | 105000 × | 105000 × | 105000 × |
| Voltage (kV) | 300 | 300 | 300 | 300 | 300 | 300 | 300 | 300 | 300 | 300 |
| Electron exposure ($e^{-}$/Å$^2$) | 48 | 48 | 48 | 48 | 48 | 48 | 48 | 48 | 48 | 48 |
| Defocus range (μm) | 0.6 - 2.0 | 0.6 - 2.0 | 0.6 - 2.0 | 0.6 - 2.0 | 0.6 - 2.0 | 0.6 - 2.0 | 0.6 - 2.0 | 0.6 - 2.0 | 0.6 - 2.0 | 0.6 - 2.0 |
| Pixel size (Å) | 0.84 | 0.84 | 0.84 | 0.84 | 0.84 | 0.84 | 0.84 | 0.84 | 0.84 | 0.84 |
| Extraction box size (pixels) | 400 | 400 | 400 | 400 | 400 | 400 | 400 | 400 | 400 | 400 |
| Symmetry imposed | C1 | C1 | C1 | C1 | C1 | C1 | C1 | C1 | C1 | C1 |
| Initial particle images (no.) | 678562 | 678562 | 678562 | 678562 | 678562 | 678562 | 678562 | 678562 | 678562 | 678562 |
| Final particle images (no.) | 142788 | 98109 | 92836 | 51079 | 49975 | 48262 | 46777 | 46250 | 45965 | 39655 |
| Map resolution (Å) | 3.2 | 3.3 | 3.4 | 3.4 | 3.4 | 3.5 | 3.4 | 3.6 | 3.7 | 3.6 |
| FSC threshold | 0.143 | 0.143 | 0.143 | 0.143 | 0.143 | 0.143 | 0.143 | 0.143 | 0.143 | 0.143 |
| Map resolution range (Å) | 1.8 - 28 | 2.0 - 32 | 2.0 - 33 | 2.1 - 33 | 2.2 - 32 | 2.2 - 38 | 2.0 - 35 | 2.1 - 39 | 2.2 - 39 | 2.2 - 43 |
| **Map composition** | | | | | | | | | | |
| CobS:CobT-Nterm:CobT-Cterm | 6:5:1 | 6:4:1 | 6:4:1 | 6:5:1 | 6:3:1 | 5:3:1 | 5:4:1 | 6:4:1 | 6:3:1 | 5:5:1 |
| **Model building and refinement** | | | | | | | | | | |
| Initial model used (PDB code) | - | - | - | - | - | - | - | - | - | - |
| Model resolution (Å) | 3.3 | - | - | - | - | - | - | - | - | - |
| FSC threshold | 0.5 | - | - | - | - | - | - | - | - | - |
| Model resolution range (Å) | 1.8 - 5.9 | - | - | - | - | - | - | - | - | - |
| Map sharpening *B* factor (Å$^2$) | 98.7 | 96.4 | 96.9 | 81.4 | 84.6 | 86.7 | 81.5 | 89.5 | 86.2 | 84.2 |
| Model composition | | | | | | | | | | |
| Non-hydrogen atoms (no.) | 24590 | - | - | - | - | - | - | - | - | - |
| Protein residues (no.) | 3125 | - | - | - | - | - | - | - | - | - |
| Ligands (no.) | 5 | - | - | - | - | - | - | - | - | - |
| *B* factors (Å$^2$) | | | | | | | | | | |
| Protein | 212.8 | - | - | - | - | - | - | - | - | - |
| Ligand | 112.4 | - | - | - | - | - | - | - | - | - |
| R.m.s. deviations | | | | | | | | | | |
| Bond lengths (Å) | 0.003 | - | - | - | - | - | - | - | - | - |
| Bond angles (°) | 0.552 | - | - | - | - | - | - | - | - | - |
| Validation | | | | | | | | | | |
| MolProbity score | 1.72 | - | - | - | - | - | - | - | - | - |
| Clashscore | 8.86 | - | - | - | - | - | - | - | - | - |
| Poor rotamers (%) | 1.13 | - | - | - | - | - | - | - | - | - |
| Ramachandran plot | | | | | | | | | | |
| Favored (%) | 96.76 | - | - | - | - | - | - | - | - | - |
| Allowed (%) | 3.24 | - | - | - | - | - | - | - | - | - |
| Disallowed (%) | 0.00 | - | - | - | - | - | - | - | - | - |

### References


(1) Abramson, J.; Adler, J.; Dunger, J.; Evans, R.; Green, T.; Pritzel, A.; Ronneberger, O.; Willmore, L.; Ballard, A. J.; Bambrick, J.; Bodenstein, S. W.; Evans, D. A.; Hung, C.-C.; O'Neill, M.; Reiman, D.; Tunyasuvunakool, K.; Wu, Z.; Žemgulytė, A.; Arvaniti, E.; Beattie, C.; Bertolli, O.; Bridgland, A.; Cherepanov, A.; Congreve, M.; Cowen-Rivers, A. I.; Cowie, A.; Figurnov, M.; Fuchs, F. B.; Gladman, H.; Jain, R.; Khan, Y. A.; Low, C. M. R.; Perlin, K.; Potapenko, A.; Savy, P.; Singh, S.; Stecula, A.; Thillaisundaram, A.; Tong, C.; Yakneen, S.; Zhong, E. D.; Zielinski, M.; Žídek, A.; Bapst, V.; Kohli, P.; Jaderberg, M.; Hassabis, D.; Jumper, J. M. Accurate Structure Prediction of Biomolecular Interactions with AlphaFold 3. *Nature* **2024**, *630* (8016), 493–500. https://doi.org/10.1038/s41586-024-07487-w

(2) Kahle, M.; Appelgren, S.; Elofsson, A.; Carroni, M.; Ädelroth, P. Insights into the Structure-Function Relationship of the NorQ/NorD Chaperones from Paracoccus Denitrificans Reveal Shared Principles of Interacting MoxR AAA+/VWA Domain Proteins. *BMC Biol* **2023**, *21* (1), 47. https://doi.org/10.1186/s12915-023-01546-w

(3) Kahle, M.; Appelgren, S.; König, F.; Carroni, M.; Ädelroth, P.; Wendler, P. NorQD AAA+ Complex Drives Metal Insertion by a Twisting Mechanism. *Nat Commun* **2026**, *17* (1), 3032. https://doi.org/10.1038/s41467-026-71044-4

(4) Robert, X.; Guillon, C.; Gouet, P. FoldScript: A Web Server for the Efficient Analysis of AI-Generated 3D Protein Models. *Nucleic Acids Research* **2025**, *53* (W1), W277–W282. https://doi.org/10.1093/nar/gkaf326

(5) Pettersen, E. F.; Goddard, T. D.; Huang, C. C.; Meng, E. C.; Couch, G. S.; Croll, T. I.; Morris, J. H.; Ferrin, T. E. UCSF ChimeraX: Structure Visualization for Researchers, Educators, and Developers. *Protein Science* **2021**, *30* (1), 70–82. https://doi.org/10.1002/pro.3943

(6) Lundqvist, J.; Elmlund, D.; Heldt, D.; Deery, E.; Söderberg, C. A. G.; Hansson, M.; Warren, M.; Al-Karadaghi, S. The AAA+ Motor Complex of Subunits CobS and CobT of Cobaltochelatase Visualized by Single Particle Electron Microscopy. *Journal of Structural Biology* **2009**, *167* (3), 227–234. https://doi.org/10.1016/j.jsb.2009.06.013
